# The expanded immunoregulatory protease network in mosquitoes is governed by gene co-expression

**DOI:** 10.1101/2024.06.18.599423

**Authors:** Bianca Morejon, Kristin Michel

## Abstract

Serine protease cascades regulate key innate immune responses. In mosquitoes, these cascades involve clip-domain serine proteases and their non-catalytic homologs (CLIPs), forming a complex network whose make-up and structural organization is not fully understood. This study assessed the impact of 85 CLIPs on humoral immunity in *Anopheles gambiae*. By coupling RNAi with assays measuring antimicrobial activity and melanization, we identified 27 CLIPs as immunoregulators that together form two distinct subnetworks. CLIPs regulating antimicrobial activity were found to control infection resistance, as knockdowns reduced bacterial load and improved survival. Furthermore, our analysis of CLIP gene expression unveiled a novel immunoregulatory mechanism reliant on protease baseline co-expression rather than infection-induced upregulation. These findings underscore that despite its complexity mosquito immune regulation may be targeted for malaria interventions.

---

The Afrotropical mosquito *Anopheles gambiae* is the primary vector responsible for the transmission of malaria parasites in most of sub-Saharan Africa. As malaria prevention relies on controlling mosquito populations, novel approaches hinge on a detailed molecular grasp of mosquito physiology, including innate immunity and its regulation (*1–3*). This knowledge is crucial for our fundamental understanding of vector competence in this most important human vector-borne disease (*4–6*).

Serine protease cascades play essential roles in a wide range of innate defense responses in animals, including blood clotting in mammals and horseshoe crab (*7, 8*), and complement in mammals (*9*). In insects, these cascades consist of clip-serine proteases (cSPs) and their non-catalytic homologs (cSPHs), together known as CLIPs (*10*). Immune cascades involving CLIPs lead to the specific activation of prophenoloxidase (proPO), crucial for the melanization response (*11–13*). Melanization is a fast-acting immune response deployed against microbial infections, including those caused by bacteria, ciliates, oomycetes, fungi, filarial worms, and malaria parasites (*14*). The process involves synthesizing melanin and cross-linking it with molecules on microbial surfaces, effectively killing the invading pathogens (*12*). Given its broad-spectrum effectiveness, melanization is a potential target for novel control strategies, including the development of parasite transmission blockers and late-life acting (LLA) insecticides (*15, 16*). Furthermore, CLIP cascades can also result in the proteolytic activation of the cytokine Spätzle (SPZ) (*17–19*), ultimately leading to the expression of Toll-regulated genes, including well-characterized antimicrobial peptides (AMPs). These small molecules are rapidly secreted into the hemolymph, where they disrupt microbial cell membranes or interfere with microbial physiological processes (*20*). Endogenous and exogenous AMPs have been expressed as gene drive effectors in transgenic mosquitoes, representing promising control tools to block parasite development (*2, 21*).

Genetic and biochemical studies in *An. gambiae* have identified multiple cSPs and cSPHs regulating proPO activity (*13, 22–32*). These are organized into two functional modules: the cSP module, which comprises one or more interacting protease cascades, and the cSPH module, which consists of a hierarchical arrangement of protease homologs (*25*). However, the identity and structural organization of these modules governing melanization are only partially defined, and the CLIP cascade leading to Toll pathway activation, and thus, antimicrobial activity, remains uncharacterized. To address these knowledge gaps, we developed a functional genetic screen to assess the impact of CLIPs on hemolymph antimicrobial activity and melanization in *An. gambiae* mosquitoes.

## An RNAi screen identifies 27 CLIPs as immunoregulators in *An. gambiae*

The *An. gambiae* genome contains 110 CLIP genes, which are divided into 5 subfamilies (A-E), forming one of the largest protein families in mosquitoes (*33*). To specifically test the contribution of CLIPs to melanization and hemolymph antimicrobial activity, we coupled RNAi with the Melanization Associated Spot Assay (MelASA) and the Zone of Inhibition (ZOI) assay in our screen strategy (Fig. 1A) (*27, 34*). RNAi was employed to individually silence 85 out of the 110 CLIP genes across all subfamilies (table S1), with each individual CLIP gene silenced efficiently and specifically (fig. S1 and table S2). Mosquitoes injected with dsRNA were challenged with the non-pathogenic and highly immunogenic *Micrococcus luteus* and evaluated for their immune response. We validated this approach by silencing established positive and negative immunoregulators. Specifically, we targeted the proPO activation pathway components *CLIPB4* and *Serine protease inhibitor 2* (*SRPN2*), known to be required and inhibitory for melanization, respectively (*5, 32*). Likewise, we silenced the Toll pathway components *REL1* and *CACTUS*, both known as necessary and inhibitory factors for antimicrobial activity, respectively (*34*).

**Fig. 1.**
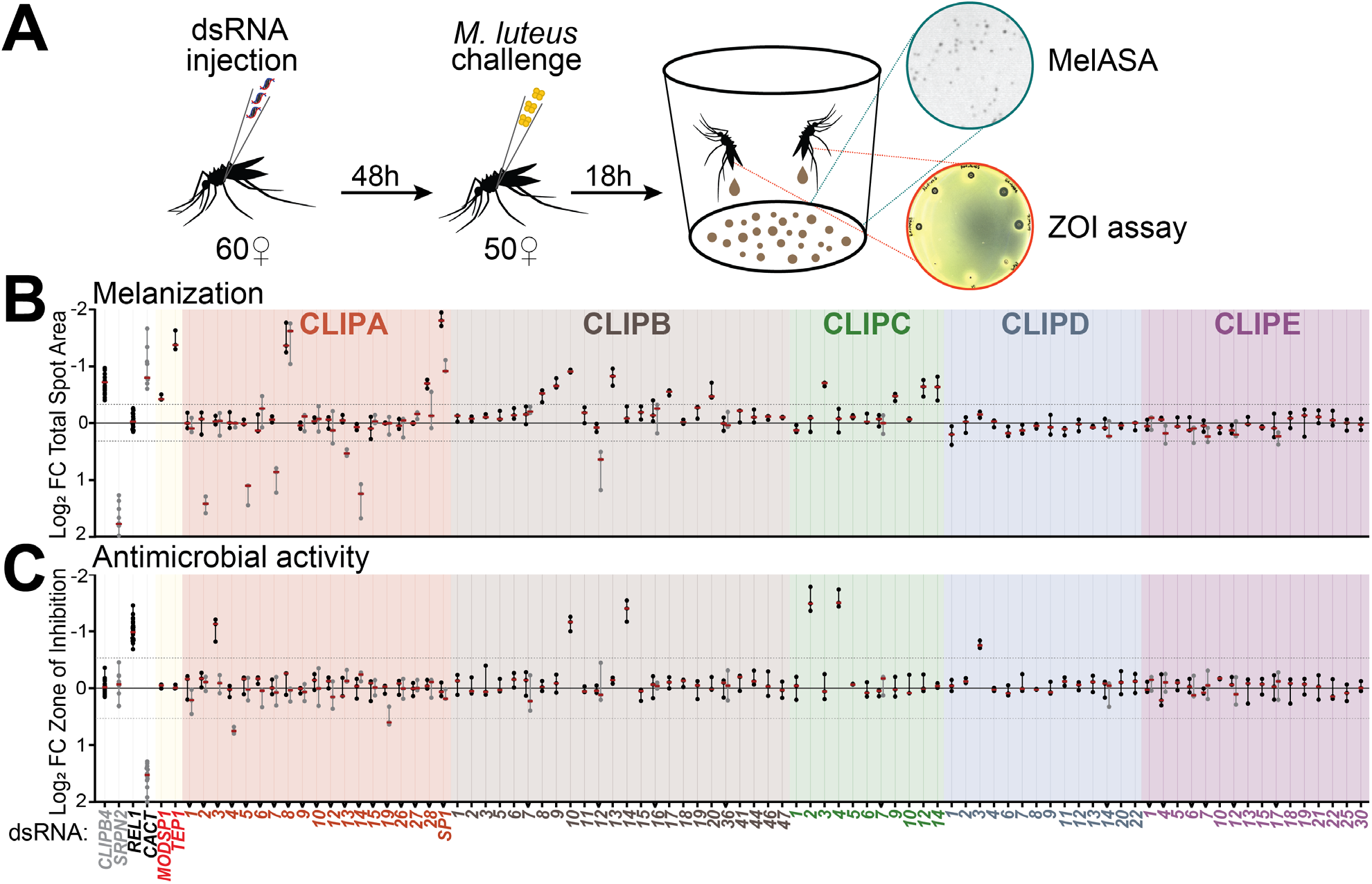
CLIPs act as positive and negative regulators of melanization and antimicrobial activity. (**A**) Schematic of screen strategy. Batches of 60 female mosquitoes were injected with dsRNA to individually silence 85 CLIP genes. DsRNA injected mosquitoes were challenged with 50.6 nL of resuspended lyophilized *M. luteus* at OD_600_ = 5 and 0.1 for putative positive and negative regulators, respectively. The MelASA and ZOI assays were performed 18 h after challenge using the same mosquitoes. (**B**) MelASA RNAi screen. Data shown as log2 fold change (FC) Total spot area relative to ds*GFP* injected mosquitoes challenged with lyophilized *M. luteus* suspension (n=50). *CLIPB4* and *SRPN2* kds were used as positive controls for positive and negative regulators, respectively. (**C**) ZOI RNAi screen. Data shown as log2 FC ZOI relative to ds*GFP* injected mosquitoes challenged with lyophilized *M. luteus* suspension (n=40). *REL1* and *CACT* kds were used as positive controls for positive and negative regulators, respectively. Batches of 7-10 candidate genes were tested in independent groups using their own positive and negative controls. Dashed lines represent the thresholds established to identify candidate regulators (defined in ‘Methods’ section). Red line and error bars represent median ± interquartile range from three biological replicates.

The RNAi screen identified 27 out of the 85 *CLIP* genes as immunoregulators, with 20 genes regulating melanization (Fig. 1B, table S3 and S4), and eight genes regulating antimicrobial activity (Fig. 1B, table S5 and S6). The MelASA phenotypes were found within the CLIPA (8), B (6), and C (4) subfamilies, with no identified phenotypes in subfamilies D or E. Similarly, the ZOI phenotypes belonged mostly to the CLIPA (3), B (2) and C (2) subfamilies. However, unlike melanization, we identified one CLIP D as a regulator of antimicrobial activity. Notably, only *CLIPB10* kd induced both MelASA and ZOI phenotypes (Fig. 1B and C), revealing minimal overlap between the cascades regulating melanization and antimicrobial activity.

To delve deeper into the potential common immune factors between melanization and antimicrobial activity, we expanded our screen to include components recognized as upstream regulators of CLIP cascades. Specifically, we tested the impact of two opsonins, the *Thioester-containing protein 1* (*TEP1*) and the ortholog of the *Drosophila melanogaster* modular serine protease (hereby named ‘*MODSP1*’), in these reactions. We observed that kds of *MODSP1* and *TEP1* met the threshold for MelASA positive regulator phenotypes, but not for ZOI phenotypes (Fig. 1B and C, table S3 and S5), suggesting that these do not contribute to hemolymph antimicrobial activity.

Overall, these results demonstrate that the set of proteases regulating humoral immunity in *An. gambiae* is much larger than previously understood. The limited overlap among CLIPs and upstream immune factors in the regulation of melanization and antimicrobial activity supports the presence of largely distinct regulatory modules in these immune reactions. This is in clear contrast to other insects such as *D. melanogaster* and *Manduca sexta*, where some of the cascade members involved in Toll pathway activation also participate in the melanization response (*35, 36*). These findings suggest that in mosquitoes, the regulation of these two immune responses has significantly diverged, and does not involve tightly integrated protease cascades.

## An expanded repertoire of CLIPs regulates melanization

Out of the 20 CLIPs that showed a MelASA phenotype, we identified *CLIPB13* and *B20*, as well as *C3, C12, C14* as novel positive regulators (Fig. 2A and table S7), and *A5, A7, A13* and *B12* as novel negative regulators of melanization (Fig. 2A and table S8). Phylogenetic analyses of the *An. gambiae* CLIP family identified *CLIPC12* and *C14* as part of the CLIPB subfamily (*33*), suggesting a downstream role in proPO activation. While *CLIPC12* and *C14* are close paralogs (*33*), we did not observe cross-silencing, confirming that the obtained phenotypes as positive regulators are due to specific silencing by their corresponding dsRNAs (fig. S1B).

**Fig. 2.**
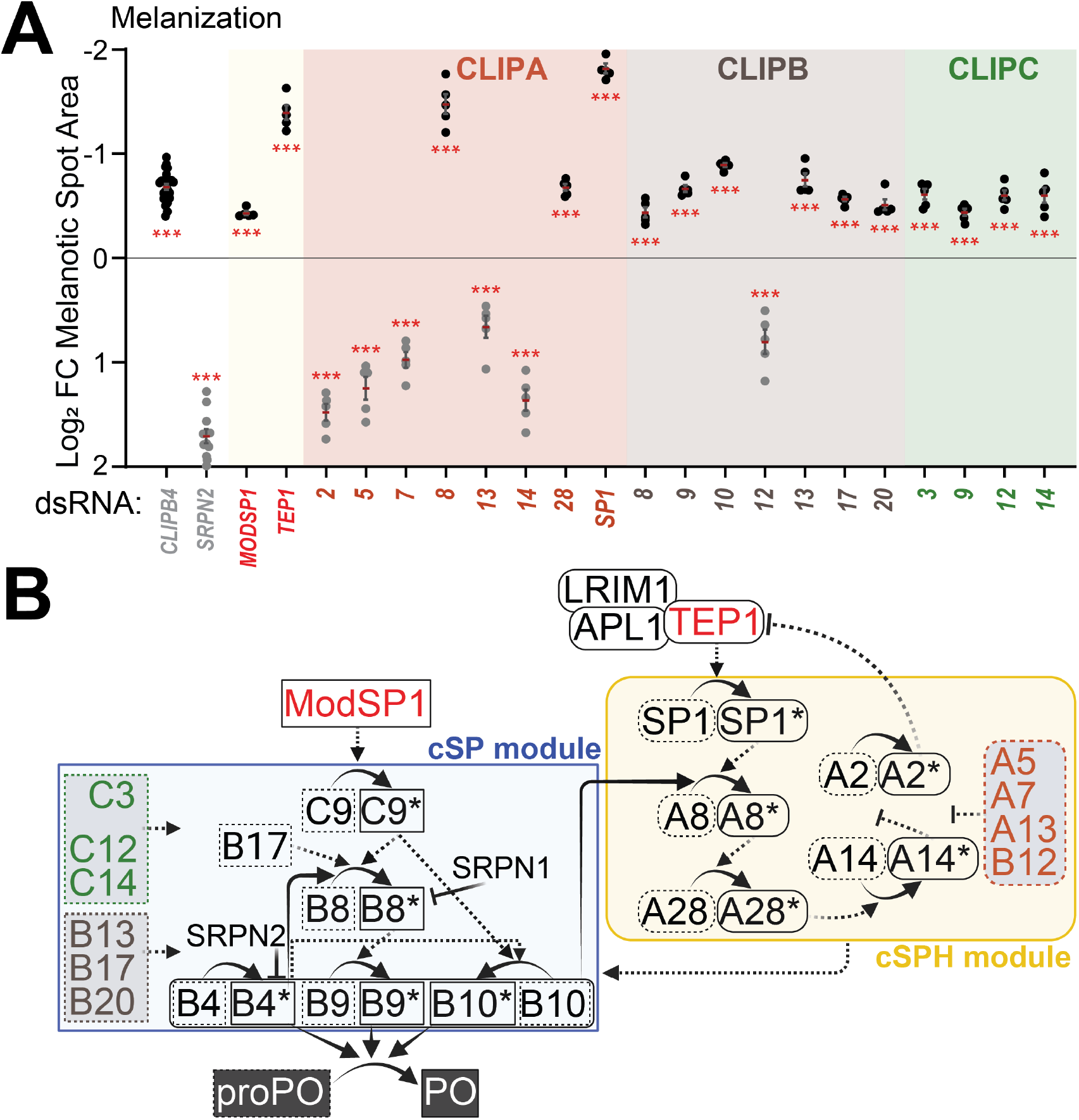
An unprecedented number of CLIPs contribute to melanization in *Anopheles gambiae*. (**A**) MelASA RNAi screen identified 20 CLIPs, MODSP1 and TEP1 as regulators of melanization. Data shown as log2 fold change (FC) Total spot area relative to ds*GFP* injected mosquitoes challenged with lyophilized *M. luteus* suspension (n=50). Knockdown of *CLIPB4* and *SRPN2* were used as positive controls for positive and negative regulators, respectively. Red line and error bars represent mean ± SE from five biological replicates. One-way ANOVA followed by Bonferroni’s post-tests were performed on ZOI data to calculate statistical significance (*** P < 0.0001). (**B**) Proposed model of the *An. gambiae* proPO activation cascades, including the interactions and hierarchies within and across the cSP (blue) and cSPH (yellow) modules. We identified 6 new cSPs that promote melanization (shown in green and brown), and 4 new cSPHs that inhibit melanization (shown in orange). Their placement within their respective modules is currently unknown. Dashed lines indicate interactions inferred from *in vivo* genetic studies. Solid lines indicate direct interactions inferred from *in vitro* biochemical assays. *Active cSP or cSPH. Arrows and lines with perpendicular bars indicate direction of positive and negative interactions, respectively.

Based on these results, combined with the current understanding of the *An. gambiae* proPO activation cascades (*13, 22–25, 27, 29, 31, 32, 37*), we propose a model depicting the potential structural organization of these immunoregulatory cascades, and the interactions between their cSP and cSPH modules (Fig. 2B). Compared to other serine protease cascades with regulatory roles in animals (*7–9*), the *An. gambiae* melanization cascades feature an expanded number of proteases and form a complex regulatory network. Within this network, cSPs and cSPHs interact positively and negatively at multiple levels linking pathogen recognition with recruitment of proPO activity to pathogen surfaces.

## Regulators of humoral antimicrobial activity control resistance to bacterial infection

Our results identified *CLIPA3, B10, B14, C2, C4* and *D3* as positive regulators and *CLIPA4* and *A19* as negative of antimicrobial activity (Fig. 3A, tables S9 and S10). These findings represent the first report of CLIPs regulating antimicrobial activity in *An. gambiae* mosquitoes. This also marks the first identification of a CLIPD as a regulator of humoral immunity in *An. gambiae*, and only the second such instance across all insects (*38*). Importantly, this is the first description of cSPHs as regulators of hemolymph antimicrobial activity, suggesting a novel mechanism to control humoral immunity in insects.

**Fig. 3.**
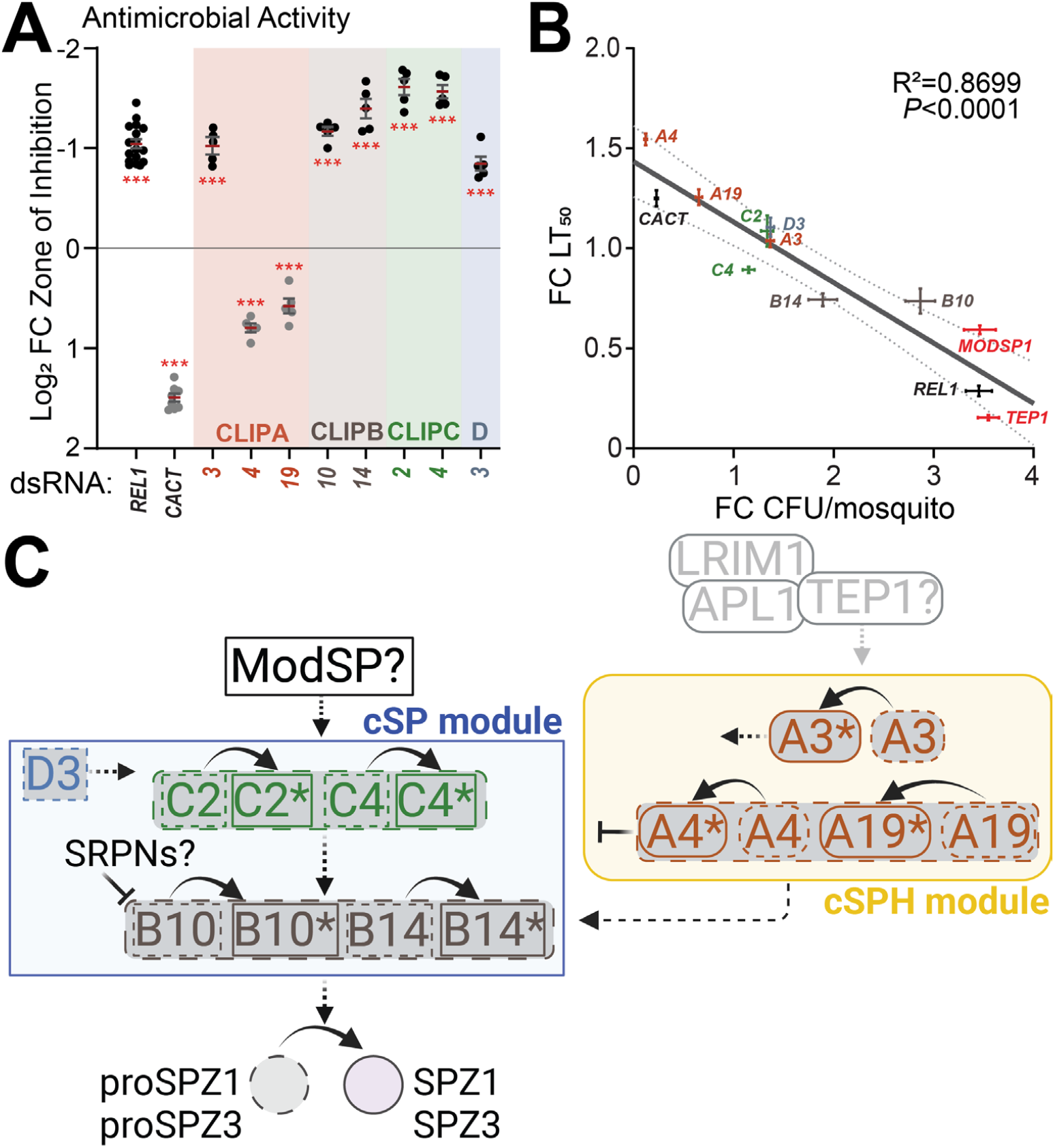
A CLIPs cascade orchestrates antimicrobial defenses that are crucial for resistance to bacterial infection in *Anopheles gambiae*. (**A**) ZOI RNAi screen identified 8 CLIPs as regulators of antimicrobial activity. Data shown as log2 fold change (FC) ZOI relative to ds*GFP* injected mosquitoes challenged with *M. luteus* suspension (n=40). *REL1* and *CACT* kds were used as positive controls for positive and negative regulators, respectively. Red line and error bars represent mean ± SE from five biological replicates. One-way ANOVA followed by Bonferroni’s post-tests were performed on ZOI data to calculate statistical significance (*** P < 0.0001). (**B**) Higher bacterial load after silencing regulators of antimicrobial activity correlated with lower median survival. Data shown as mean log2 FC Colony Forming Units (CFU)/mosquito and Lethal Time 50 (LT_50_) ± SE (n=5) plotted on linear x and y-axes, respectively. Solid line represents linear regression, dashed lines show 95% confidence bands. Pearson correlation coefficient= -0.9327. (**C**) Proposed model of the *An. gambiae* SPZ activation cascades, including hierarchies within and across the cSP (blue) and cSPH (yellow) modules. We identified 6 cSPs that promote, and 2 cSPHs that inhibit antimicrobial activity. *Active cSP or cSPH.

CLIP serine proteases regulating insect antimicrobial activity are primarily recognized for cleaving SPZ proteins (*17, 35*). As the SPZ ligand for TOLL in mosquitoes remains unidentified, we assessed the impact of the six annotated *SPZ* genes in the *An. gambiae* genome on antimicrobial activity. Kd of *SPZ1* and *SPZ3* produced a significant reduction in ZOI (fig. S2A and table S11), phenocopying the antimicrobial activity observed after silencing *REL1* and *MYD88*, which are required intracellular components of the Toll signal transduction pathway (*34*). Phenotypes were attributed to the individual silencing of *SPZ1* and *SPZ3*, as no evidence of cross-silencing between them was detected (fig S2B). In addition, from the six annotated *SPZ* genes, only *SPZ1* and *SPZ3* were expressed in adult female mosquitoes (table S12), as shown in our previously published RNASeq data (*39*). Therefore, we incorporated SPZ1 and SPZ3 as putative TOLL ligands in the SPZ activation cascade in *An. gambiae*.

Multiple studies have emphasized the function of antimicrobial factors in resistance and tolerance defense mechanisms (*40, 41*). Therefore, we asked whether immunoregulatory CLIPs alter the outcome of *An. gambiae* infection with its opportunistic pathogen *Enterococcus faecalis*. We found a negative correlation between bacterial load and median survival after kd of positive and negative regulators of antimicrobial activity (Fig. 3B). Silencing the negative regulators *CLIPA4* and *A19* reduced bacterial load, and significantly increased median survival (fig. S3, table S13 and S14) compared to ds*GFP* controls. In turn, kd of the positive regulators *CLIPA3, C2, B10, B14*, and *D3* increased bacterial load. Among these, *CLIPB10* and *B14* kds reduced median survival, while no difference was observed after *CLIPA3, C2* and *D3* kds. Neither median survival nor bacterial load were impacted after kd of the positive regulator *CLIPC4*. Kd of *MODSP1* and *TEP1* also produced an increase in bacterial load and reduction in median survival (Fig. 3B, table S13 and S14).

Overall, our results showed that higher bacterial load after silencing immunoregulatory CLIPs was correlated with lower median survival, indicating that the regulators of humoral antimicrobial activity contribute to resistance to infection. Based on the current knowledge of the SPZ activation cascades in other insects, we propose for the first time a layout of the potential structural organization of these cascades in *An. gambiae* mosquitoes, and the interactions between their cSP and cSPH components (Fig. 3C).

## Humoral immune regulators are characterized by co-expression

Genes that encode CLIP family of proteases and their homologs are upregulated transcriptionally after infection with pathogenic and non-pathogenic bacteria; indeed, *CLIP*s are often among those genes identified in RNASeq experiments that display a strong transcriptional response to environmental stimuli (*42–44*). Using our previously published RNASeq data (*39*), we therefore tested whether gene expression levels or changes upon infection were able to discriminate between immunoregulatory and non-phenotypic CLIPs.

The normalized expression levels in unchallenged (UC) females were on average significantly higher for CLIPs that regulate humoral immunity, as compared to those that did not (Fig. 4A, table S15). We also observed that 15% of non-phenotypic CLIPs were not expressed in adult females (table S15). When we removed all CLIP genes that were not expressed from the analysis, expression levels of immunoregulatory CLIPs remained elevated as compared to those without such function (fig. S4A and B). We obtained the same result when comparing expression levels in mosquitoes challenged with pathogenic (*E. faecalis*) and non-pathogenic (*M. luteus*) bacteria (fig. S4C-F). While 66% of CLIPs were positively regulated by infection (fig. S4G-H, J-K), no difference was observed on the average fold change between immunoregulatory and non-phenotypic CLIPs after challenge (Fig 4B, fig. S4L, table S15). This suggested that at least for CLIPs, upregulation after infection is not a predictor for function in humoral immunity in *An. gambiae*.

**Fig. 4.**
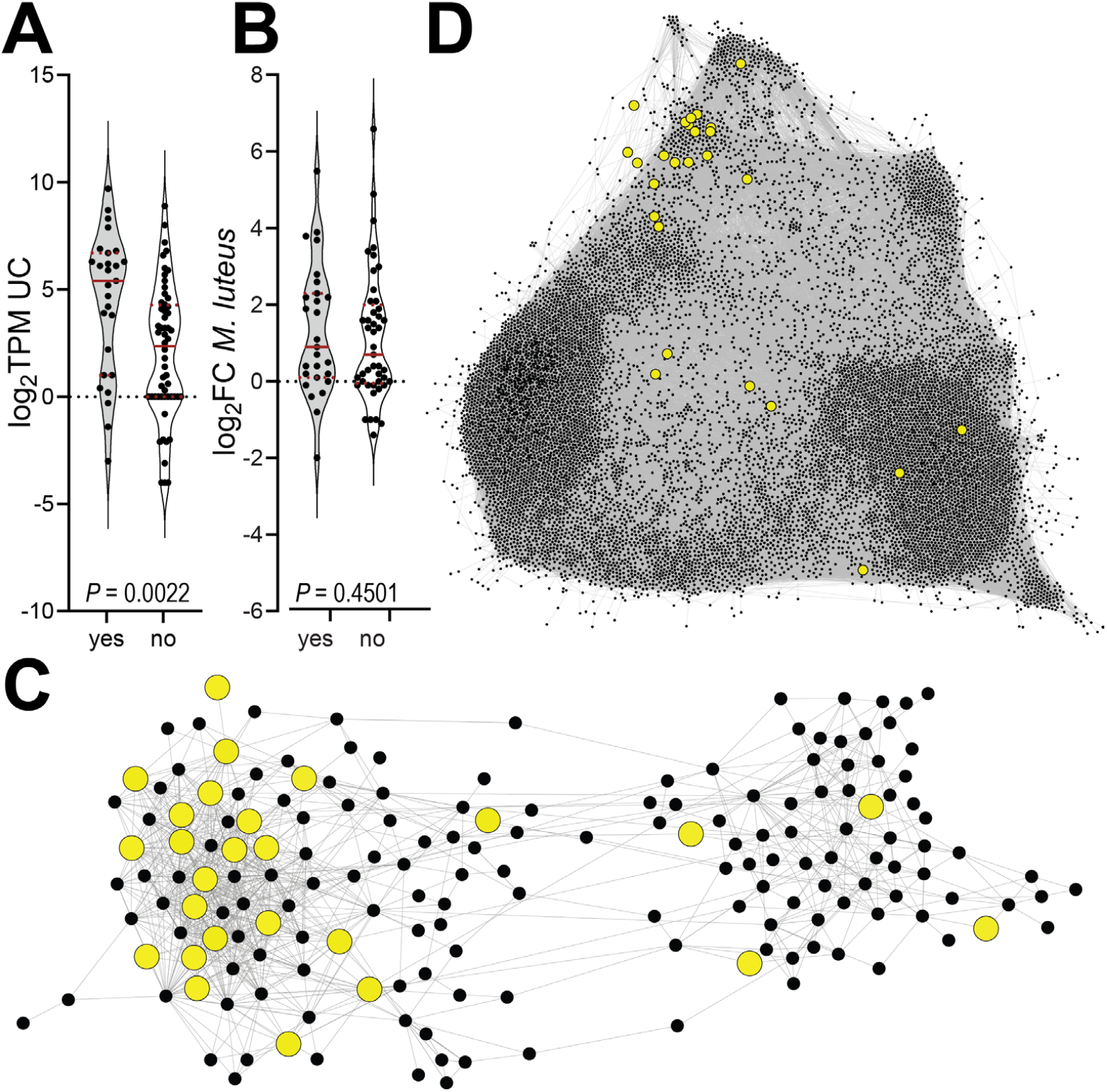
Immunoregulatory CLIPs are globally co-expressed, but not regulated by infection. (**A**) Expression levels of immunoregulatory CLIPs (yes), as measured by log2 Transcripts per million (TPM), are significantly higher in naïve adult female mosquitoes as compared non-phenotypic CLIPs (no). (**B**) However, no difference is observed in fold change after *M. luteus* challenge between these two. Violin plots show the comparisons indicated in each panel, with red solid and dashed lines representing median and interquartile range limits, respectively. *P* values of unpaired two-tailed t-tests are shown in each panel. (**C, D**) The AgMelGCN2.0 (**C**) and the AgGCN1.0 networks (**D**). Positive and negative immunoregulatory CLIPs cluster mostly in one community of both networks, AgMelGCN (Community 2) and AgGCN1.0 (Community 7). Network images generated in Gephi 0.9.2 (Forceatlas 2, settings: Thread s number, 19; Tolerance, 1.0; approximate repulsion, checked; Scaling, 5.0; Gravity, 1.0; Prevent Overlap, checked; Edge Weight influence, 1.0)).

By examining node centrality measures in gene co-expression networks established in *An. gambiae*, we recently identified CLIPB4 as a hub in a gene co-expression network, called AgMelGCN (*32*). To determine whether high centrality is a general feature of immunoregulatory CLIPs, we updated the existing network and built AgMelGCN2.0 using our recently published pipeline (table S16) (*45*). We then reanalyzed node centralities for all CLIPs in the network (table S17). Immunoregulatory CLIPs, on average, exhibited higher strength, eigenvector, and closeness centrality in AgMelGCN2.0 compared to non-phenotypic CLIPs, while betweenness remained similar (fig. S5A-D). These data showed that immunoregulatory CLIPs are co-expressed with members of the genes families that control melanization, and suggested that immunoregulatory CLIPs are co-expressed on a transcriptional level. To test this directly, we examined the distribution of immunoregulatory CLIPs in the network, and observed them clustering significantly in community 2 (Fig. 4C, table S18). In contrast, non-phenotypic CLIPs were enriched in community 4 (fig. S6A-C, table S18), demonstrating distinct co-expression patterns and transcriptional regulation between immunoregulatory and non-phenotypic CLIPs.

To determine whether this observed co-expression was an artifact of the gene selection used to build AgMelGCN2.0, we performed the same analyses in the previously published transcriptome-wide AgGCN1.0, built on the same 256 experimental conditions as AgMelGCN2.0 (*45*). Overall, the CLIP gene family was nearly 10-fold enriched in community 7, as compared to all genes in the network (Fig. 4D, table S19). This enrichment was largely driven by immunoregulatory and less so by non-phenotypic CLIPs (15-fold vs. 8-fold enrichment in community 7, fig. S6D-F). Non-phenotypic CLIPs also clustered in community 2 (2-fold enrichment) (Fig. 4D). Together, these data confirmed that transcriptional regulation differs between immunoregulatory and non-phenotypic CLIPs, and that immunoregulatory CLIPs are globally co-expressed.

## Discussion

Our comprehensive genetic screen revealed a complex network of immunoregulatory proteases playing pivotal roles in defense mechanisms. By integrating our genetic approach with gene expression and co-expression analyses, our findings underscore the complexity and unique features of humoral immunoregulation in mosquitoes.

Our MelASA screen recapitulated the phenotypes of most of the previously characterized regulators of melanization (Fig. 2A and table S7 and S8) (*13, 22–25, 27, 29, 31, 32, 37*). Nevertheless, we noticed some variability between our screen results and published melanization phenotypes (*23, 30*) or *in vitro* biochemical activities of cSPHs (*46*). The evident variability may be the result of the same cSPH performing distinct roles upstream and downstream of the core proteolytic activity resulting in activation cleavage of proPO. Immunoregulatory cSPHs can serve as positive and negative immunoregulators of TEP1-mediated opsonization and recruitment of cSPs to the pathogen surface (*24, 27, 47*) and as efficient cofactors of proPO activity (*46*). Downstream functions may be masked by phenotypes caused due to upstream functions and vice versa. In addition, the differences between the regulatory roles of cSPHs could stem from microbial class-specific functions linked to specific components of the melanization network (*29, 30*), which could be captured by repeated cSPH-focused screens using an expanded repertoire of microbial challenges.

Our findings have potential practical applications for developing new mosquito control strategies, as a dysregulated immune response can reduce vectorial capacity by shortening the lifespan of adult females (*5, 47*). In addition, recent data suggest that immunoregulatory CLIPs such as CLIPB4 are in part responsible for the wAlbB-mediated suppression of human malaria parasite infection in this *Anopheles stephensi* mosquito strain (*48*). Immunoregulatory CLIPs could also be directly exploited to block parasite transmission at both early (*13, 22, 24, 29*) and late stages of *P. berghei* and *P. falciparum* (*49*). Since these CLIPs regulate the melanization response, it would be interesting to investigate whether CLIPs involved in antimicrobial activity can also be targeted, given their crucial roles in mosquito fitness and infection resistance.

## Acknowledgments

We would like to thank all members of the Michel laboratory for mosquito rearing. We thank Dr. Lynn Hancock, University of Kansas for the kind gift of the *Enterococcus faecalis* strain. We gratefully acknowledge Dr. Ed Brokesh and the Intelligent Agar Solutions team (Ryan Cunningham, Seth Hoffman, Dugan Hult, Elizabeth Seidl) Kansas State University, Reno, for the design of the agar stencil. Figure panels 1A, 2B, and 3B were generated with Biorender.com. Finally, sincere thanks go to Dr. Michael R. Kanost, Kansas State University, for his continued support.

## Funding

National Institutes of Health R01AI140760 (KM)

USDA National Institute of Food and Agriculture Hatch project 1021223 (KM)

Kansas State GRIP Seed grant (KM)

This is contribution No. 24-249-J from the Kansas Agricultural Experiment Station.

The contents of this article are solely the responsibility of the authors and do not necessarily represent the official views of the funding agencies.

## Author contributions

Conceptualization: BM, KM

Methodology: BM, KM

Investigation: BM, KM

Visualization: BM, KM

Funding acquisition: KM

Project administration: KM

Supervision: KM

Writing – original draft: BM, KM

Writing – review & editing: BM, KM

## Competing interests

Authors declare that they have no competing interests.

## Data and materials availability

All data are available in the main text or the supplementary materials.

## Supplementary Materials

Materials and Methods

Supplementary Text

Figs. S1 to S6

Tables S1 to S19

References (*1-59*)

## Notes

### Competing Interest Statement

The authors have declared no competing interest.

### Summary of Updates

Revised manuscript has a separate discussion section.

## References and Notes

1. A. Ickowicz, S. D. Foster, G. R. Hosack, K. R. Hayes, Predicting the spread and persistence of genetically modified dominant sterile male mosquitoes. Parasit. Vectors 14, 480 (2021).

2. T. Nolan, Control of malaria-transmitting mosquitoes using gene drives. Philos. Trans. R. Soc. Lond. B. Biol. Sci. 376, 20190803 (2021).

3. M. Sicard, M. Bonneau, M. Weill, Wolbachia prevalence, diversity, and ability to induce cytoplasmic incompatibility in mosquitoes. Curr. Opin. Insect Sci. 34, 12–20 (2019).

4. S. Blandin, S.-H. Shiao, L. F. Moita, C. J. Janse, A. P. Waters, F. C. Kafatos, E. A. Levashina, Complement-like protein TEP1 is a determinant of vectorial capacity in the malaria vector Anopheles gambiae. Cell 116, 661–670 (2004).

5. K. Michel, A. Budd, S. Pinto, T. J. Gibson, F. C. Kafatos, Anopheles gambiae SRPN2 facilitates midgut invasion by the malaria parasite Plasmodium berghei. EMBO Rep. 6, 891–897 (2005).

6. E. L. G. Pryzdial, A. Leatherdale, E. M. Conway, Coagulation and complement: Key innate defense participants in a seamless web. Front. Immunol. 13, 918775 (2022).

7. L. Cerenius, K. Söderhäll, Coagulation in Invertebrates. J. Innate Immun. 3, 3–8 (2010).

8. K. J. Kearney, J. Butler, O. M. Posada, C. Wilson, S. Heal, M. Ali, L. Hardy, J. Ahnström, D. Gailani, R. Foster, E. Hethershaw, C. Longstaff, H. Philippou, Kallikrein directly interacts with and activates Factor IX, resulting in thrombin generation and fibrin formation independent of Factor XI. Proc. Natl. Acad. Sci. 118, e2014810118 (2021).

9. D. Ricklin, G. Hajishengallis, K. Yang, J. D. Lambris, Complement: a key system for immune surveillance and homeostasis. Nat. Immunol. 11, 785–797 (2010).

10. M. R. Kanost, H. Jiang, Clip-domain serine proteases as immune factors in insect hemolymph. Curr. Opin. Insect Sci. 11, 47–55 (2015).

11. C. An, M. Zhang, Y. Chu, Z. Zhao, Serine protease MP2 activates prophenoloxidase in the melanization immune response of Drosophila melanogaster. PloS One 8, e79533 (2013).

12. H. Kan, C.-H. Kim, H.-M. Kwon, J.-W. Park, K.-B. Roh, H. Lee, B.-J. Park, R. Zhang, J. Zhang, K. Söderhäll, N.-C. Ha, B. L. Lee, Molecular control of phenoloxidase-induced melanin synthesis in an insect. J. Biol. Chem. 283, 25316–25323 (2008).

13. X. Zhang, M. Li, L. El Moussawi, S. Saab, S. Zhang, M. A. Osta, K. Michel, CLIPB10 is a Terminal Protease in the Regulatory Network That Controls Melanization in the African Malaria Mosquito Anopheles gambiae. Front. Cell. Infect. Microbiol. 10, 585986 (2020).

14. L. C. Bartholomay, K. Michel, Mosquito Immunobiology: The Intersection of Vector Health and Vector Competence. Annu. Rev. Entomol. 63, 145–167 (2018).

15. M. L. Simões, Y. Dong, G. Mlambo, G. Dimopoulos, C-type lectin 4 regulates broadspectrum melanization-based refractoriness to malaria parasites. PLoS Biol. 20, e3001515 (2022).

16. A. F. Read, P. A. Lynch, M. B. Thomas, How to Make Evolution-Proof Insecticides for Malaria Control. PLOS Biol. 7, e1000058 (2009).

17. I.-H. Jang, N. Chosa, S.-H. Kim, H.-J. Nam, B. Lemaitre, M. Ochiai, Z. Kambris, S. Brun, C. Hashimoto, M. Ashida, P. T. Brey, W.-J. Lee, A Spätzle-processing enzyme required for toll signaling activation in Drosophila innate immunity. Dev. Cell 10, 45–55 (2006).

18. Z. Kambris, S. Brun, I.-H. Jang, H.-J. Nam, Y. Romeo, K. Takahashi, W.-J. Lee, R. Ueda, B. Lemaitre, Drosophila immunity: a large-scale in vivo RNAi screen identifies five serine proteases required for Toll activation. Curr. Biol. CB 16, 808–813 (2006).

19. K.-B. Roh, C.-H. Kim, H. Lee, H.-M. Kwon, J.-W. Park, J.-H. Ryu, K. Kurokawa, N.-C. Ha, W.-J. Lee, B. Lemaitre, K. Söderhäll, B.-L. Lee, Proteolytic cascade for the activation of the insect toll pathway induced by the fungal cell wall component. J. Biol. Chem. 284, 19474–19481 (2009).

20. Y. Shai, Mode of action of membrane active antimicrobial peptides. Pept. Sci. 66, 236–248 (2002).

21. W. Kim, H. Koo, A. M. Richman, D. Seeley, J. Vizioli, A. D. Klocko, D. A. O’Brochta, Ectopic Expression of a Cecropin Transgene in the Human Malaria Vector Mosquito Anopheles gambiae (Diptera: Culicidae): Effects on Susceptibility to Plasmodium. J. Med. Entomol. 41, 447–455 (2004).

22. L. El Moussawi, J. Nakhleh, L. Kamareddine, M. A. Osta, The mosquito melanization response requires hierarchical activation of non-catalytic clip domain serine protease homologs. PLoS Pathog. 15, e1008194 (2019).

23. J. Nakhleh, G. K. Christophides, M. A. Osta, The serine protease homolog CLIPA14 modulates the intensity of the immune response in the mosquito Anopheles gambiae. J. Biol. Chem. 292, 18217–18226 (2017).

24. M. Povelones, L. Bhagavatula, H. Yassine, L. A. Tan, L. M. Upton, M. A. Osta, G. K. Christophides, The CLIP-domain serine protease homolog SPCLIP1 regulates complement recruitment to microbial surfaces in the malaria mosquito Anopheles gambiae. PLoS Pathog. 9, e1003623 (2013).

25. S. A. Saab, X. Zhang, S. Zeineddine, B. Morejon, K. Michel, M. A. Osta, Insight into the structural hierarchy of the protease cascade that regulates the mosquito melanization response. Microbes Infect. 26, 105245 (2024).

26. A. K. D. Schnitger, F. C. Kafatos, M. A. Osta, The Melanization Reaction Is Not Required for Survival of Anopheles gambiae Mosquitoes after Bacterial Infections. J. Biol. Chem. 282, 21884–21888 (2007).

27. G. L. Sousa, R. Bishnoi, R. H. G. Baxter, M. Povelones, The CLIP-domain serine protease CLIPC9 regulates melanization downstream of SPCLIP1, CLIPA8, and CLIPA28 in the malaria vector Anopheles gambiae. PLOS Pathog. 16, e1008985 (2020).

28. J. Volz, M. A. Osta, F. C. Kafatos, H.-M. Müller, The Roles of Two Clip Domain Serine Proteases in Innate Immune Responses of the Malaria Vector Anopheles gambiae. J. Biol. Chem. 280, 40161–40168 (2005).

29. J. Volz, H.-M. Muller, A. Zdanowicz, F. C. Kafatos, M. A. Osta, A genetic module regulates the melanization response of Anopheles to Plasmodium. Cell. Microbiol. 8, 1392–1405 (2006).

30. R. Zakhia, M. A. Osta, CLIPA7 Exhibits Pleiotropic Roles in the Anopheles gambiae Immune Response. J. Innate Immun. 15, 317–332 (2023).

31. X. Zhang, C. An, K. Sprigg, K. Michel, CLIPB8 is part of the prophenoloxidase activation system in Anopheles gambiae mosquitoes. Insect Biochem. Mol. Biol. 71, 106–115 (2016).

32. X. Zhang, S. Zhang, J. Kuang, K. A. Sellens, B. Morejon, S. A. Saab, M. Li, E. C. Metto, C. An, C. T. Culbertson, M. A. Osta, C. Scoglio, K. Michel, CLIPB4 Is a Central Node in the Protease Network that Regulates Humoral Immunity in Anopheles gambiae Mosquitoes. J. Innate Immun. 15, 680–696 (2023).

33. X. Cao, M. Gulati, H. Jiang, Serine protease-related proteins in the malaria mosquito, Anopheles gambiae. Insect Biochem Mol Biol 88, 48–62 (2017).

34. B. Morejon, K. Michel, A zone-of-inhibition assay to screen for humoral antimicrobial activity in mosquito hemolymph. Front. Cell. Infect. Microbiol. 13, 891577 (2023).

35. C. An, J. Ishibashi, E. J. Ragan, H. Jiang, M. R. Kanost, Functions of Manduca sexta Hemolymph Proteinases HP6 and HP8 in Two Innate Immune Pathways. J. Biol. Chem. 284, 19716–19726 (2009).

36. T. Shan, Y. Wang, K. Bhattarai, H. Jiang, An evolutionarily conserved serine protease network mediates melanization and Toll activation in Drosophila. Sci. Adv. 9, eadk2756 (2023).

37. C. An, A. Budd, M. R. Kanost, K. Michel, Characterization of a regulatory unit that controls melanization and affects longevity of mosquitoes. Cell. Mol. Life Sci. 68, 1929–1939 (2011).

38. X. Cao, H. Jiang, Building a platform for predicting functions of serine protease-related proteins in Drosophila melanogaster and other insects. Insect Biochem. Mol. Biol. 103, 53–69 (2018).

39. B. Hixson, L. Huot, B. Morejon, X. Yang, P. Nagy, K. Michel, N. Buchon, The transcriptional response in mosquitoes distinguishes between fungi and bacteria but not Gram types. BMC Genomics 25, 353 (2024).

40. J. Huang, Y. Lou, J. Liu, P. Bulet, C. Cai, K. Ma, R. Jiao, J. A. Hoffmann, S. Liégeois, Z. Li, D. Ferrandon, A Toll pathway effector protects Drosophila specifically from distinct toxins secreted by a fungus or a bacterium. Proc. Natl. Acad. Sci. U. S. A. 120, e2205140120 (2023).

41. K. Troha, J. H. Im, J. Revah, B. P. Lazzaro, N. Buchon, Comparative transcriptomics reveals CrebA as a novel regulator of infection tolerance in D. melanogaster. PLOS Pathog. 14, e1006847 (2018).

42. M. Bonizzoni, W. A. Dunn, C. L. Campbell, K. E. Olson, M. T. Dimon, O. Marinotti, A. A. James, RNA-seq analyses of blood-induced changes in gene expression in the mosquito vector species, Aedes aegypti. BMC Genomics 12, 82 (2011).

43. K. Chen, T. Tang, Q. Song, Z. Wang, K. He, X. Liu, J. Song, L. Wang, Y. Yang, C. Feng, Transcription Analysis of the Stress and Immune Response Genes to Temperature Stress in Ostrinia furnacalis. Front. Physiol. 10 (2019).

44. Z. Luo, Y. Yu, Q. Zhang, Z. Bao, J. Xiang, F. Li, Comparative Transcriptome Analysis Reveals the Adaptation Mechanism to High Salinity in Litopenaeus vannamei. Front. Mar. Sci. 9 (2022).

45. J. Kuang, N. Buchon, K. Michel, C. Scoglio, A global Anopheles gambiae gene co-expression network constructed from hundreds of experimental conditions with missing values. BMC Bioinformatics 23, 170 (2022).

46. Q. Jin, Y. Wang, Y. Hu, Y. He, C. Xiong, H. Jiang, Serine protease homolog pairs CLIPA4-A6, A4-A7Δ, and A4-A12 act as cofactors for proteolytic activation of prophenoloxidase-2 and -7 in Anopheles gambiae. Insect Biochem. Mol. Biol. 164, 104048 (2024).

47. H. Yassine, L. Kamareddine, S. Chamat, G. K. Christophides, M. A. Osta, A Serine Protease Homolog Negatively Regulates TEP1 Consumption in Systemic Infections of the Malaria Vector Anopheles gambiae. J. Innate Immun. 6, 806–818 (2014).

48. V. Vandana, S. Dong, T. Sheth, Q. Sun, H. Wen, A. Maldonado, Z. Xi, G. Dimopoulos, Wolbachia infection-responsive immune genes suppress Plasmodium falciparum infection in Anopheles stephensi. PLOS Pathog. 20, e1012145 (2024).

49. S. Zeineddine, S. Jaber, S. A. Saab, J. Nakhleh, G. Dimopoulos, M. A. Osta, Late sporogonic stages of Plasmodium parasites are susceptible to the melanization response in Anopheles gambiae mosquitoes. bioRxiv [Preprint] (2024). 10.1101/2024.05.31.596773.

50. T. Horn, M. Boutros, E-RNAi: a web application for the multi-species design of RNAi reagents—2010 update. Nucleic Acids Res. 38, W332–W339 (2010).

51. V. L. Rhodes, M. B. Thomas, K. Michel, The interplay between dose and immune system activation determines fungal infection outcome in the African malaria mosquito, Anopheles gambiae. Dev. Comp. Immunol. 85, 125–133 (2018).

52. M. W. Pfaffl, A new mathematical model for relative quantification in real-time RT-PCR. Nucleic Acids Res. 29, 45e–445 (2001).

53. D. F. Sahm, J. Kissinger, M. S. Gilmore, P. R. Murray, R. Mulder, J. Solliday, B. Clarke, In vitro susceptibility studies of vancomycin-resistant Enterococcus faecalis. Antimicrob Agents Chemother 33, 1588–1591 (1989).

54. B. D. Jett, K. L. Hatter, M. M. Huycke, M. S. Gilmore, Simplified agar plate method for quantifying viable bacteria. BioTechniques 23, 648–650 (1997).

55. J. Kuang, K. Michel, C. Scoglio, GeCoNet-Tool: a software package for gene co-expression network construction and analysis. BMC Bioinformatics 24, 281 (2023).

56. T. M. Therneau, A. Elizabeth, C. Cynthia, survival: Survival Analysis, version 3. 5–7 (2023); https://lib.stat.cmu.edu/R/CRAN/web/packages/survival/index.html.

57. A. Kassambara, M. Kosinski, P. Biecek, S. Fabian, survminer: Drawing Survival Curves using “ggplot2,” version 0.4.9 (2021); https://lib.stat.cmu.edu/R/CRAN/web/packages/survminer/index.html.

58. T. L. Pedersen, patchwork: The Composer of Plots, version 1.2.0 (2024); https://lib.stat.cmu.edu/R/CRAN/web/packages/patchwork/index.html.

59. T. Graeber, Hypergeometric p-value calculator, Hypergeometric p-value calculator (2019). https://systems.crump.ucla.edu/hypergeometric/index.php.

