## Supplementary Materials for "The expanded immunoregulatory protease network in mosquitoes is governed by gene co-expression"

**The PDF file includes:**

Materials and Methods
Figs. S1 to S6
Tables S1 to S19
References (*50*-*59*)

### Materials and Methods

Mosquito rearing

*Anopheles gambiae* (s.l) of the G3 strain (BEI Resources Accession # MRA-112) were reared as described previously (*37*). Briefly, freshly hatched larvae were fed with a 2% (w/v) suspension of baker's yeast (Fleischmann's Active Dry Yeast, AB Mauri, St. Louis, MO, USA) for 48 h, followed by a slurry of 2% powdered fish food (TetraMin® Tropical Flakes, Tetra, Melle, Germany) and baker's yeast at a 2:1 ratio. Upon emergence, adult mosquitoes were provided with 8% fructose (Emprove Chemicals, Millipore Sigma, MO, USA) *ad libitum*. Female mosquitoes were blood fed using commercial heparinized horse blood (PlasVacc, Templeton, CA, USA) provided through an artificial membrane feeding system. All stages were housed in an environmental chamber with conditions set at 27°C, 80% humidity, and a 12:12-h light:dark cycle.

Double stranded RNA synthesis and *in vivo* gene silencing in female mosquitoes

DsRNA synthesis and adult female mosquito injections were performed as described previously (*37*). PCR amplification of the genes of interest was performed using the T7-tagged primers listed in table S1. Primer pair sequences for dsRNA templates were either obtained from former publications or generated using the webtool for design of RNAi constructs E-RNAi (*50*). Target-specific designs were obtained for 85 out of 110 CLIP genes annotated in the *Anopheles gambiae* reference genome (PEST 4.11). For the remaining 25 CLIPs (annotations based on (*33*)), either a dsRNA with a single gene target could not be designed, or amplification of a dsRNA construct was unsuccessful due to lack of expression in adult mosquitoes. In all experiments, 2 day old adult female mosquitoes were injected intrathoracically with 69 nL of 3 ug/uL dsRNA solution using a nanoinjector (Nanoject III, Drummond Scientific, Broomall, PA, USA). For kd control, mosquitoes were injected with an equivalent amount of the non-targeting dsRNA of Green Fluorescent Protein (ds*GFP*). Mosquitoes were allowed 48 h to recover before further manipulation.

Real Time-quantitative Polymerase Chain Reaction (RT-qPCR)

Efficiency of gene silencing was measured by RT-qPCR in mosquitoes collected 48 h after dsRNA injection. Total RNA was extracted from 8 females per treatment and replicate using TRIzol reagent (Invitrogen), and subsequently purified using an RNeasy mini kit (Qiagen, Valencia, CA, USA). cDNA was synthesized using 100 ng of purified total RNA in a 20 μL volume reaction with an iScript cDNA synthesis kit (Bio-Rad, Hercules, CA, USA), as described in (*51*). RT-qPCR was performed using 2 μL of 1:5 diluted cDNA in a 20 μL volume reaction with a PowerUp SYBR Green Master Mix (Thermo Fisher Scientific, Waltham, MA, USA), as described previously (*34*). Primers used in RT-qPCR are listed in table S2. Relative gene expression values were calculated using the method described in (*52*), taking into account the primer efficiencies (table S2). Ribosomal protein S7 (*RpS7*) expression was used as reference and ds*GFP* treatments as calibrator conditions. RT-qPCRs were performed in triplicate for each sample and primer pair from three biological replicates.

RNAi screen

A functional genetic analysis pipeline was developed to assess the impact of CLIPs on humoral immunity after bacterial challenge. The pipeline incorporated a Melanization Associated Spot Assay (MelASA) and a Zone of Inhibition (ZOI) assay to test CLIPs contribution to melanization and hemolymph antimicrobial activity in the same mosquitoes. After 48 h of dsRNA treatment, 50 female mosquitoes were challenged by injection with 50.6 nL of resuspended lyophilized *Micrococcus luteus* (MilliporeSigma, Burlington, MA, USA). To prepare the inoculum, lyophilized *M. luteus* was resuspended in 1x phosphate-buffered saline (PBS) and diluted to high ($OD_{600}$ = 5) and low ($OD_{600}$ = 0.1) doses. To test for CLIPs and immune factors acting as positive regulators of humoral immunity, dsRNA-injected mosquitoes were challenged with *M. luteus* at the high dose, while negative regulators were tested by challenging with the low dose. Sterile 1x PBS injections were used as negative controls. Following bacterial challenge, dsRNA-injected mosquitoes were housed for 18 h in experimental cups containing P8-grade white filter papers (Fisher Scientific, Pittsburgh, PA, USA) at the bottom. Batches of 7-10 candidate genes were tested in independent groups using their own positive and negative controls. Three independent biological replicates were performed using different mosquito generations. For treatments where the median of the response variable surpassed the predefined phenotype cutoff values for MelASA and ZOI assays, two extra biological replicates to enable statistical evaluation of the screen results.

Melanization phenotype analysis

To assess CLIPs impact on melanization, MelASA was performed as described previously (*27*). Briefly, the filter papers containing melanotic excreta were removed from the experimental cups 18 h post infection. Collected filter papers were imaged under white epi-illumination and without filters in an Alpha Imager 2200 system (Alpha Innotech, San Leandro, CA, USA). Image processing and total spot area measurements were performed as described by (*27*) with the modifications described in (*32*). Positive controls for melanization regulation were established by injecting mosquitoes with dsRNA targeting the prophenoloxidase-activating protease *CLIPB4* as a positive regulator, and the serine-protease inhibitor *SRPN2* as a negative regulator (*25*, *27*, *32*). Positive regulation phenotypes were defined by a log2 fold change (FC) of the median total spot area <-0.25, and negative regulation phenotypes by a log2 FC of the median >0.25. Upon the addition of two biological replicates for treatments that exceeded the phenotype cutoffs, differences in total melanotic spot area were analyzed using One-Way ANOVA, followed by Bonferroni's multiple comparisons post-test ($P < 0.05$).

Hemolymph antimicrobial activity phenotype analysis

To measure CLIPs impact on hemolymph antimicrobial activity, hemolymph collection and ZOI assays were performed as described previously (*34*). Briefly, hemolymph from 40 dsRNA-injected mosquitoes per treatment was collected and flash-frozen 18 h post infection. To prepare ZOI plates, a 1:10 mixture of live *M. luteus* (ATCC, No. 4698) at an $OD_{600}$ = 10 and 1% Luria-Bertani (LB) agar was poured onto a 9 cm diameter Petri dish (Fisher Scientific, Waltham, MA, USA). While the agar solidified, equidistant wells were created using a 3D-printed stencil with metal pins, positioned 1.5 cm from the outer edge of the petri dish. After 4 minutes, the stencil was removed, and each well was loaded with 1 µL of hemolymph per treatment. Plates were incubated upside down for 16 h at 37 °C, and subsequently imaged under white epi-illumination and without filters using the Azure 300 imaging system (Azure Biosystems Inc., Dublin, CA, USA). Image processing and ZOI measurements were performed as described in (*34*). Positive controls for antimicrobial activity regulation were included by injecting mosquitoes with dsRNA targeting the NF-κB transcription factor *REL1* as a positive regulator, and the inhibitor of κB *CACT* as a negative regulator. Positive regulation phenotypes were assigned if the median log2FC in ZOI <-0.50, while negative regulation phenotypes were assigned for median log2FC >0.50. After adding two replicates for treatments exceeding phenotype cutoffs, ZOI differences were analyzed using One-Way ANOVA with Bonferroni's post-test ($P< 0.05$).

Bacterial Proliferation and Mosquito Survival Assays

To determine CLIPs impact on resistance to bacterial infections, microbial proliferation and survival assays were adapted from the protocols developed by (*22*). Briefly, 35-50 dsRNA-injected mosquitoes were injected intrathoracically with 50.6 nL of GFP-expressing, tetracycline-resistant *Enterococcus faecalis* (strain V583, (*53*)). To prepare the bacterial challenge, 2 mL of Todd-Hewitt broth were inoculated with a single colony of *E. faecalis*, then incubated for 16 h at 37 °C. Cultures were centrifuged 20 mins at 2000 rpm, washed with 1 x PBS, resuspended in 1 ml sterile 1x PBS and diluted to an $OD_{600}= 0.08$ (corresponding to an average dose of 5.3 x $10^7$ CFUs/mL).

Batches of 8 mosquitoes per treatment were ground in 400 µL Todd-Hewitt broth 18 h post-challenge, then adjusted to a final volume of 1 mL with the same medium. To determine the number of viable *E. faecalis* cells in whole mosquito homogenates, the track dilution bacteria enumeration technique was used as described in (*54*). Homogenates were diluted $10^3$-fold, and 20 µL samples were spotted in a row onto tetracycline-selective Todd Hewitt agar (15 µg/mL) on 100×15 mm square plates (Fisher Scientific, Pittsburgh, PA, USA). Plates were tilted to a 45°–75° angle immediately after inoculation, allowing the diluted homogenate spots to migrate in parallel tracks across the agar surface to the opposite side. Once the tracks were dried, usually in less than a minute, the plates were incubated upside down for 16 h at 37 °C.

GFP-positive CFUs were scored under a fluorescence microscope (Leica EL6000, Leica Microsystems, Wetzlar, Germany) for each treatment. Ds*GFP*-injected mosquitoes were used as a negative control, while ds*REL1* and ds*CACT*-treated mosquitoes acted as positive controls for positive and negative regulation of antimicrobial activity, respectively. Candidate genes were tested in two independent groups using their own positive and negative controls. Potential differences in CFU/mosquito from five independent biological replicates were analyzed using One-way ANOVA followed by Bonferroni's multiple comparisons post-test ($P< 0.05$).

Survival of the remaining 35-40 mosquitoes per treatment was monitored daily until mortality reached 100%. Survival curves were generated with the obtained data using the Kaplan–Meier method, and statistically compared using the Log-rank (Mantel-Cox) test. Lethal time 50 ($LT_{50}$, the time point after challenge where 50 % of mosquitoes in a given treatment had died) was statistically evaluated using One-way ANOVA followed by Bonferroni's multiple comparisons post-test. Correlation between FC CFU/mosquito and FC $LT_{50}$ was determined using Pearson correlation.

CLIP gene expression analysis

Gene expression levels for the 85 CLIPs analyzed in the RNAi screen were derived from a prior RNAseq study examining the transcriptomic response to various microbial systemic infections in *An. gambiae* (*39*). Expression data normalized as log2 transcripts per million (TPM) was extracted for unchallenged (UC) mosquitoes, as well as mosquitoes at 12 h post challenge with either *M. luteus* or *E. faecalis.* Expression data for immunoregulatory vs. non-phenotypic CLIPs exhibiting a normal distribution were analyzed with an unpaired two-tailed t-test, while those not meeting the normal distribution criteria were assessed using a two-tailed Mann-Whitney test.

Gene-co expression analysis

To test for gene co-expression among CLIPs we rebuilt a gene co-expression network (GCN) based on the z-score normalized gene co-expression matrix we had assembled previously (*32*) using the GeCoNet-Tool (*55*) with a bin size of 10 and a 0.005 cutoff. The resulting undirected weighted network, which we named AgMelGCN2.0, contained 177 nodes and 958 edges. The edge list for this network can be accessed in table S16. We then used the GeCoNet-Tool to define communities based on the Louvain algorithm, the network core, and node centralities as described previously (*55*). Characteristics of each node in the AgMelGCN2.0 are summarized in table S17. Node centralities and community membership were used as measures to assess potential differences in gene co-expression between immunoregulatory vs. non-phenotypic CLIPs. Differences in node centralities in AgMelGCN2.0 were analyzed using a two-tailed Mann-Whitney test. Enrichment in community membership was assessed in the AgMelGCN2.0 as well as in the transcriptome-wide AgGCN1.0 network (*45*) using a hypergeometric test.

Statistical analyses

Statistical analyses were conducted using GraphPad Prism 10 Software (GraphPad Software, USA) unless specified otherwise. Normality of the data was assessed using Shapiro-Wilk tests. Parametric tests were applied to normally distributed data, and non-parametric tests were used for non-normally distributed data, as indicated for each assay. Survival curves were plotted, and log-rank pairwise comparisons were performed in R (version 4.2.0) using the packages *survival* (*56*), *survminer* (*57*), and *patchwork* (*58*). Node enrichments in network communities were assessed by calculating the *P* value for under- or over-enrichment based on the cumulative distribution function of the hypergeometric distribution in Excel according to (*59*).

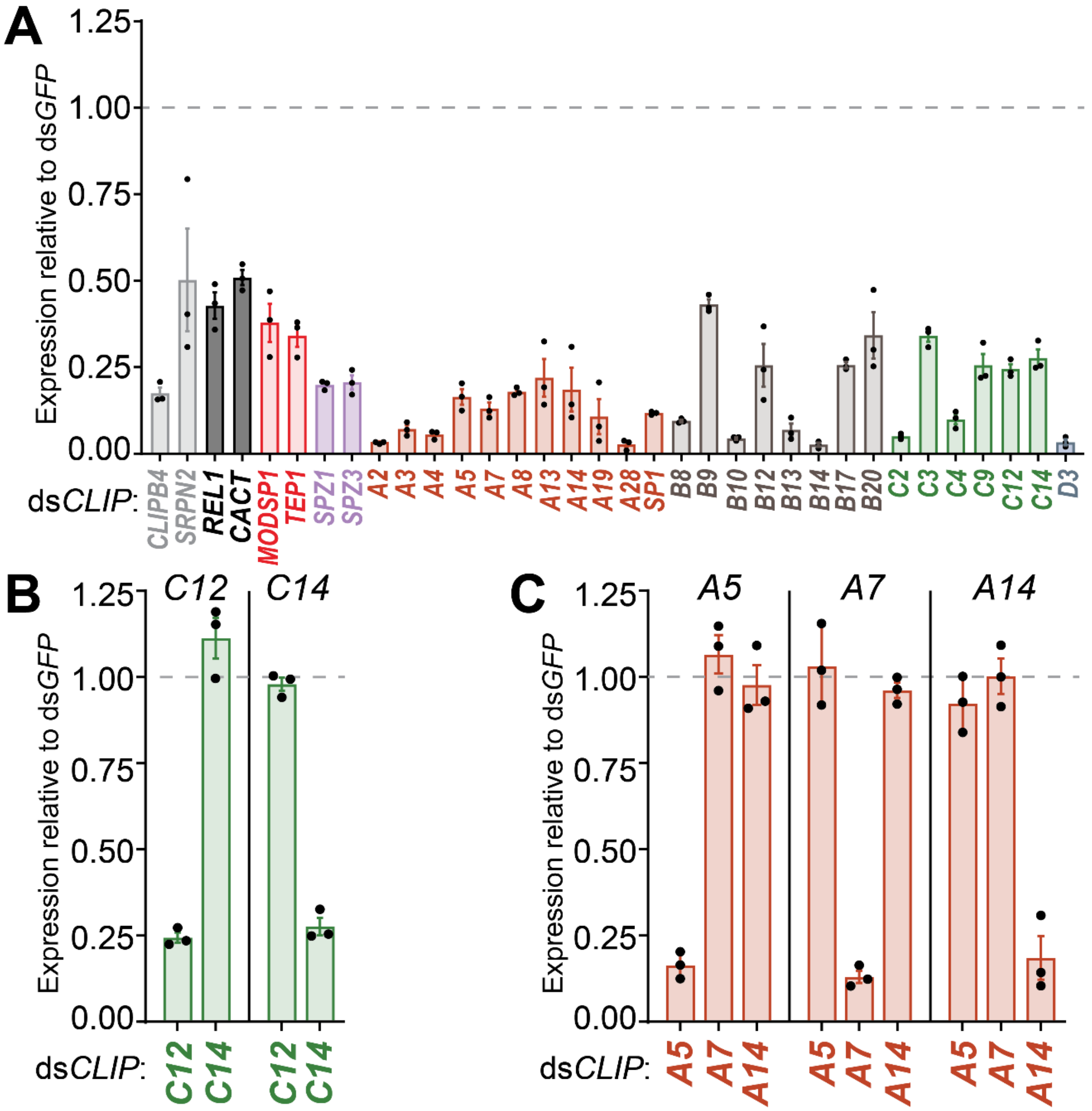


**Fig. S1. RNAi silencing efficiency and specificity**. (**A**) Silencing efficiencies for dsRNA injections of *CLIP* genes and other genes of interest that have immunoregulatory functions. Average knockdown efficiency for each individual target was ≥ 50%. (**B-C**) Silencing specificity of dsRNA targeting *CLIPC12* and *C14* (**B**), *A5*, *A7*, *A14* (**C**). No evidence for cross-silencing was detected. Relative expression measured by RT-qPCR in whole female mosquitoes collected 48 h after dsRNA injection (n=8). Bars represent mean ± SE. Transcript levels were normalized to the housekeeping ribosomal gene *RpS7* and calculated as the expression of the target gene relative to the ds*GFP* control. RT-qPCRs were performed in triplicate for each sample and primer pair from three biological replicates.

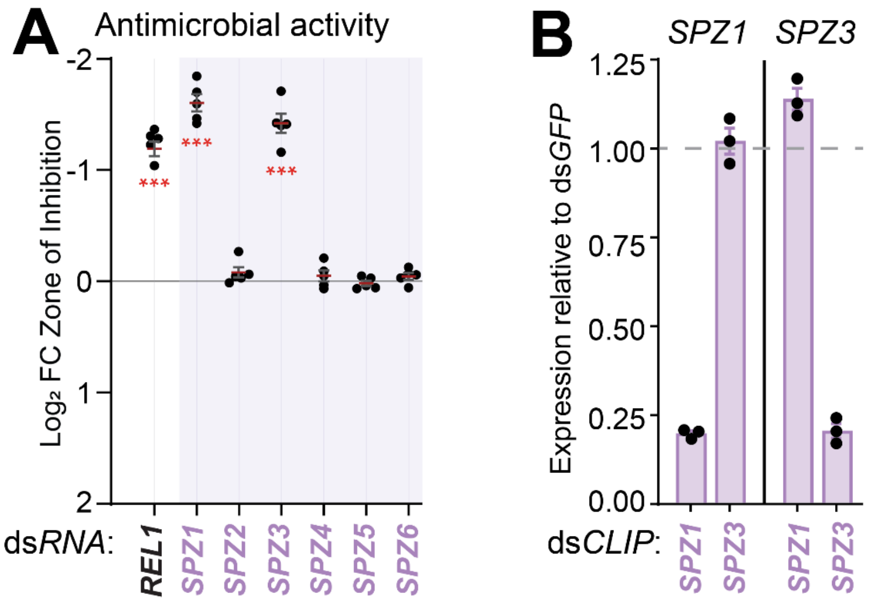


**Fig S2. SPZ1 and SPZ3 are putative TOLL ligands**. (**A**) Hemolymph antimicrobial activity after dsRNA injection targeting *Spätzle* genes (*SPZ1-6*). *SPZ1* and *SPZ3* kd significantly reduced antimicrobial activity compared to ds*GFP*-injected control mosquitoes. Data shown as log2 fold change (FC) Zone of Inhibition (ZOI) relative to ds*GFP* injected mosquitoes challenged with 50.6 nL of lyophilized *Micrococcus luteus* suspension at an $OD_{600}$=5. *REL1* kd was used as a positive control for positive regulation of antimicrobial activity. Red line and error bars represent mean ± SE (n=5; 40 female mosquitoes per replicate). One-way ANOVA followed by Bonferroni's post-tests were performed on ZOI data to calculate statistical significance (*** $P < 0.0001$). (**B**) Silencing efficiency and specificity for dsRNA injections of *SPZ1* and *SPZ3*. No evidence for cross-silencing was detected. Relative expression measured by RT-qPCR in whole female mosquitoes collected 48 h after dsRNA injection (n=8). Bars represent mean ± SE. Transcript levels were normalized to the housekeeping ribosomal gene *RpS7* and calculated as the expression of the target gene relative to the ds*GFP* control. RT-qPCRs were performed in triplicate for each sample and primer pair from three biological replicates.

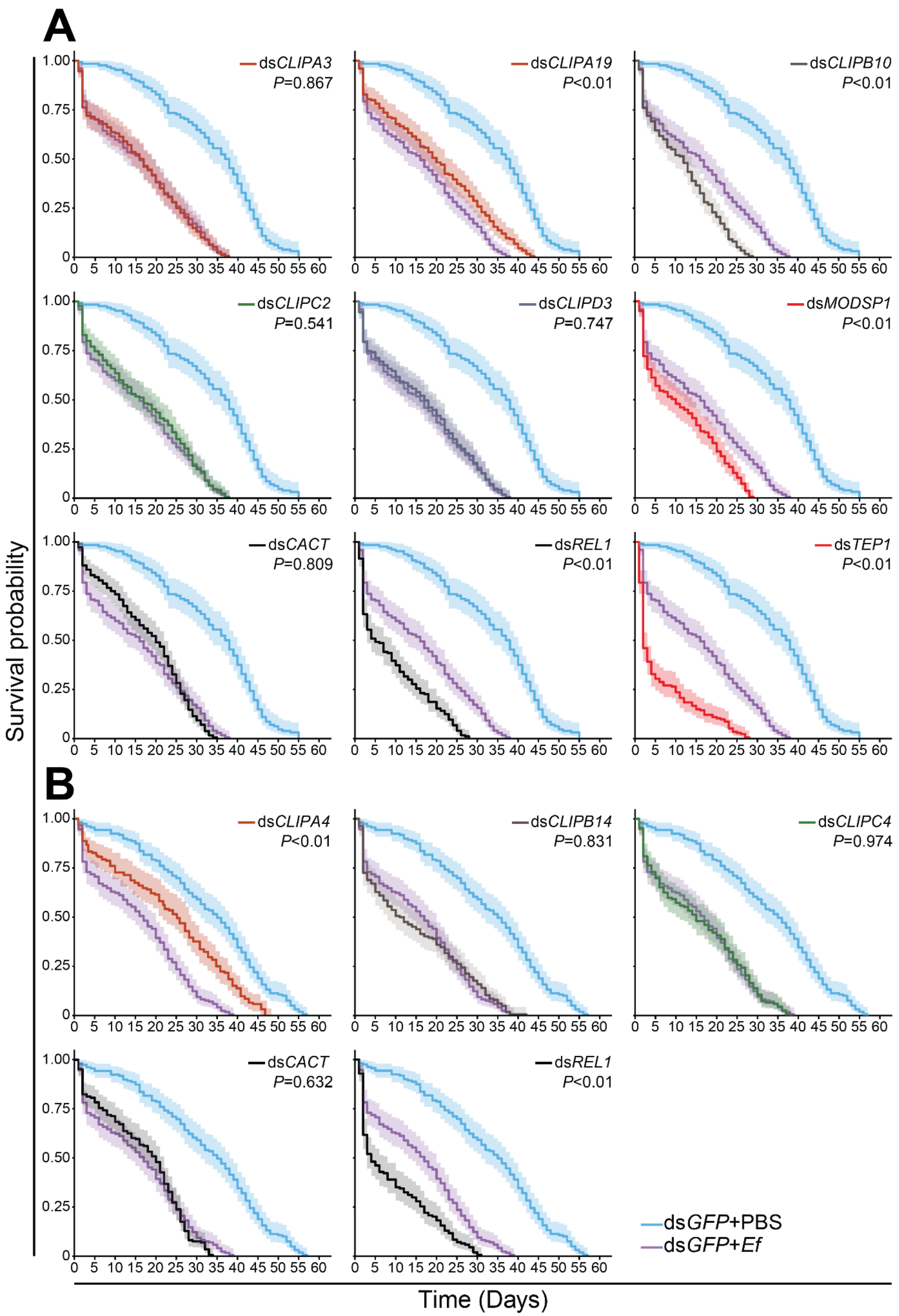
A
B
Survival probability
Time (Days)
dsCLIPA3 P=0.867
dsCLIPA19 P<0.01
dsCLIPB10 P<0.01
dsCLIPC2 P=0.541
dsCLIPD3 P=0.747
dsMODSP1 P<0.01
dsCACT P=0.809
dsREL1 P<0.01
dsTEP1 P<0.01
dsCLIPA4 P<0.01
dsCLIPB14 P=0.831
dsCLIPC4 P=0.974
dsCACT P=0.632
dsREL1 P<0.01
dsGFP+PBS
dsGFP+Ef

**Fig. S3. Impact of antimicrobial activity regulators on mosquito survival to infection.** Survival curves of dsRNA injected mosquitoes challenged with GFP-expressing, tetracycline-resistant *Enterococcus faecalis.* Candidate genes were tested in two independent groups (A and B) using their own positive (ds*CACT* -negative regulator; ds*REL1*-positive regulator) and negative (ds*GFP* prior to phosphate-buffered saline (PBS) injection) controls. Survival curves represent mean cumulative survival probability and 95% confidence intervals (CI) using 35-40 female mosquitoes per treatment per replicate (n=5). Statistical comparisons of survival curves after each dsRNA treatment relative to *E. faecalis*-challenged ds*GFP*-injected mosquitoes were assessed by Log-rank test, with resulting *P* values shown in each survival plot.

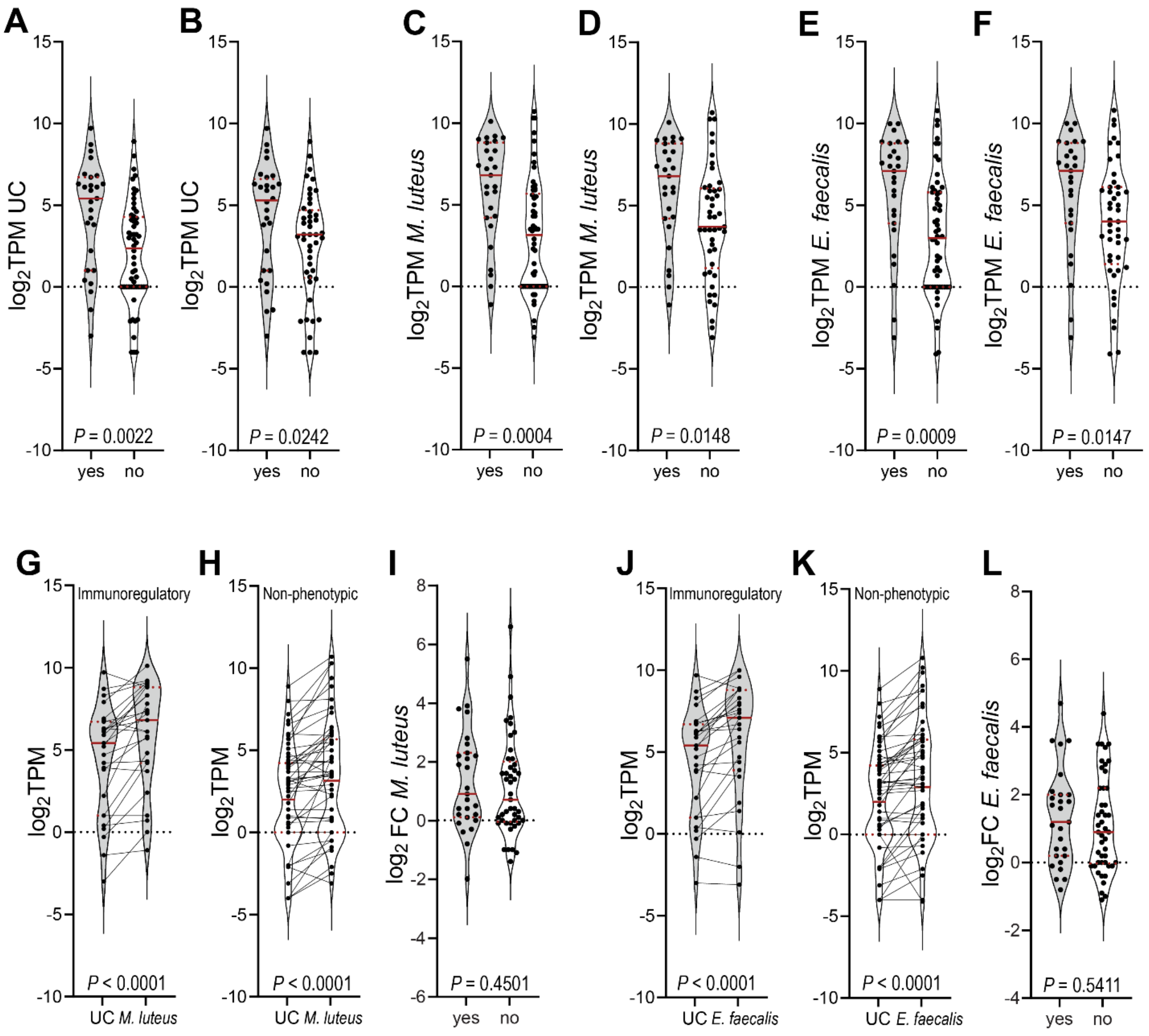


**Fig. S4. Clip gene expression patterns.** Expression of immunoregulatory CLIPs in naïve adult female (UC-unchallenged) mosquitoes is higher than that of non-phenotypic CLIPs (**A-F**). Data normalized as log2 transcripts per million (TPM) for immunoregulatory (yes) vs. non-phenotypic (no) CLIP genes. CLIP gene expression in UC mosquitoes (**A**, **B**) 12 h after *Micrococcus luteus* challenge (**C**, **D**) and 12 h after *Enterococcus faecalis* challenge (**E, F**). Panels **A, C, E** show expression data for all CLIP genes analyzed in the screen, while data for genes not expressed in adult females are omitted in panels **B, D, F**. In response to bacterial challenges, gene expression levels increase for both, immunoregulatory and non-phenotypic CLIPs (**G-L**). Lines connect the expression level of an individual CLIP gene in UC and *M. luteus* (**G, H**) or *E. faecalis* (**J, K**) infection conditions. Log2 fold change (FC) in CLIPs gene expression relative to *M. luteus* (**I**) and *E. faecalis* (**L**) challenges. Violin plots show the comparisons indicated in each panel, with red solid and dashed lines representing median and interquartile range, respectively. *P* values of unpaired two-tailed t-tests (**A-D, I**), Mann-Whitney U tests (**E-F, L**) and Wilcoxon matched-pairs sign tests **(G, H, J, K**) are shown in each panel.

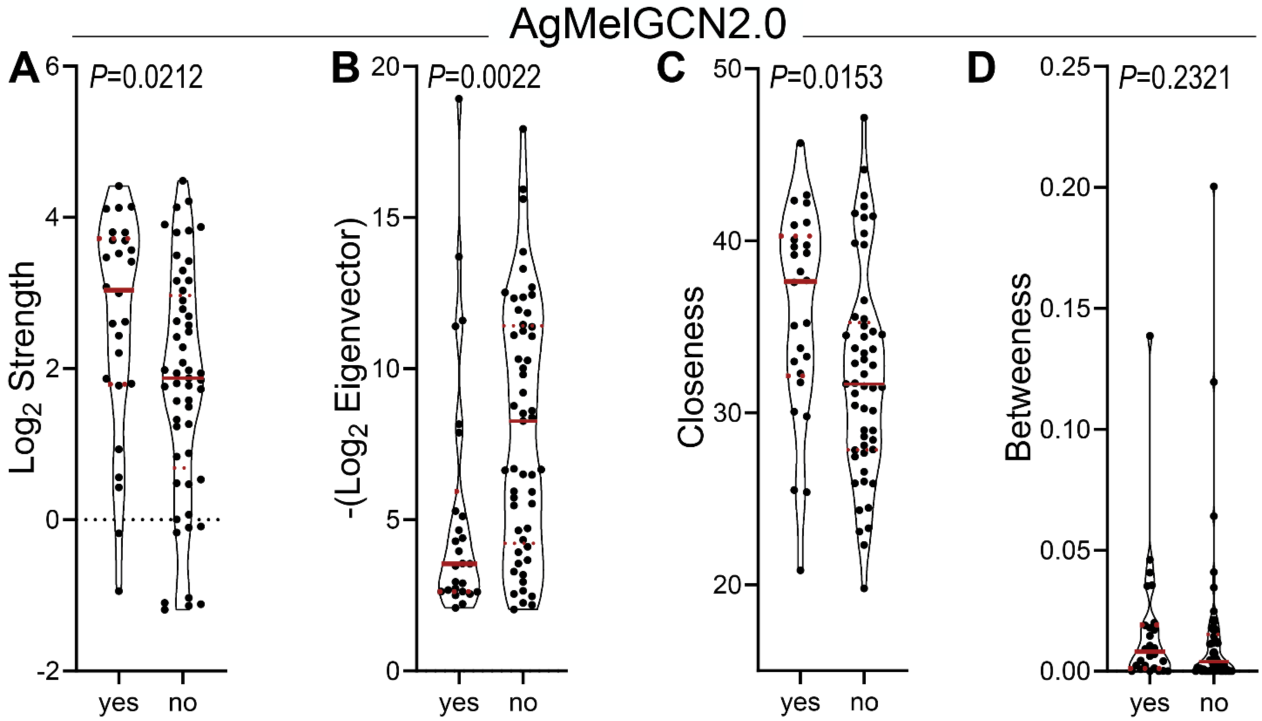


**Fig. S5. CLIPs centrality measures within the gene co-expression network of genes with putative roles in melanization (AgMelGCN2.0).** Immunoregulatory CLIPs (yes) showed significantly higher strength (**A**), eigenvector (**B**), and closeness (**C**) centralities compared to non-phenotypic CLIPs (no). Betweenness (**D**) centrality remained similar between the two groups. Strength and eigenvector measures were log2 and -log2 transformed, respectively, for CLIP genes in AgMelGCN2.0. Violin plots show the comparisons indicated in each panel, with red solid and dashed lines representing median and interquartile range (IR) limits, respectively. *P* values of unpaired two-tailed t-tests (**C**) and Mann-Whitney U tests (**A, B, D**) are shown in each panel.

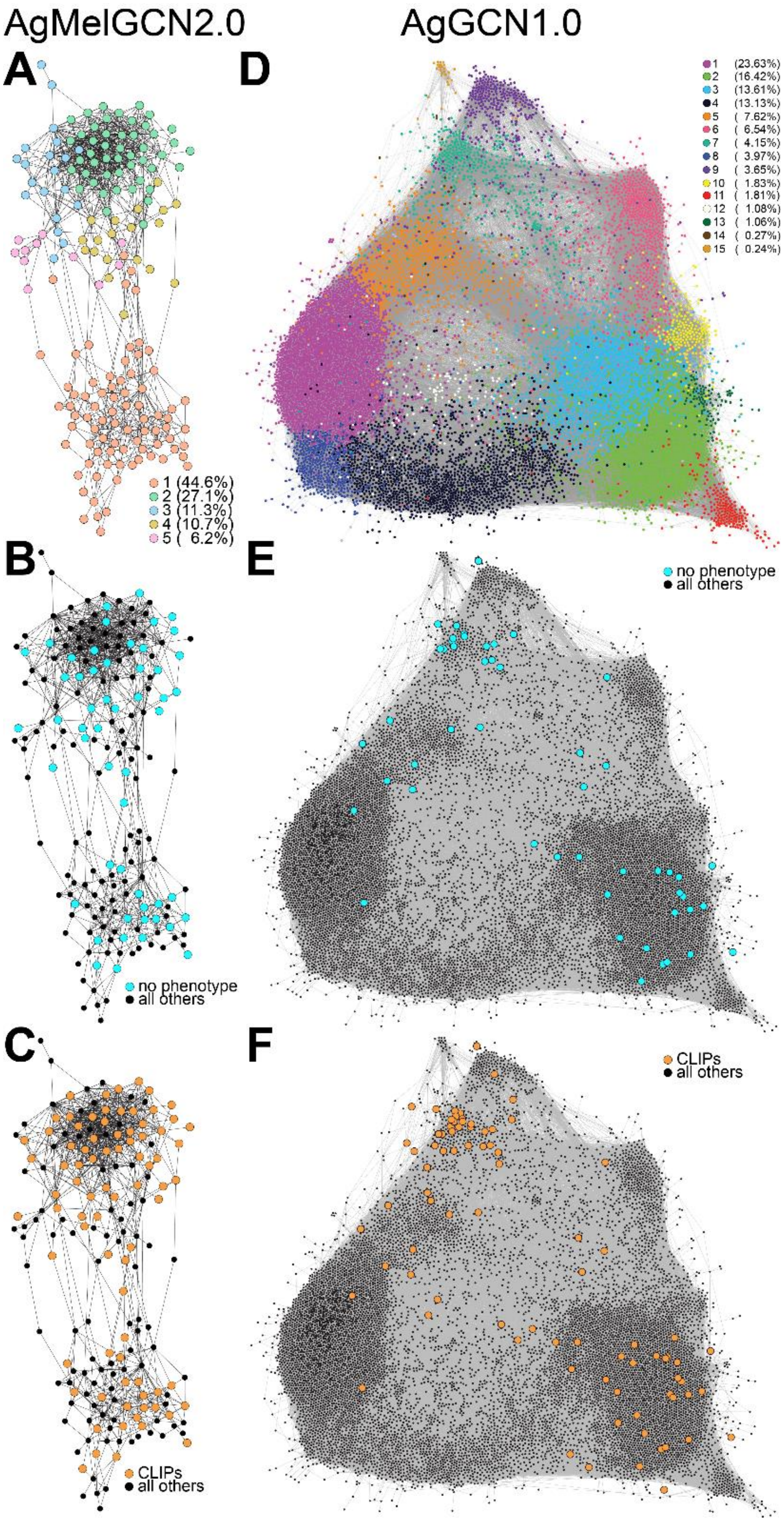
AgMelGCN2.0
AgGCN1.0
A
D
1 (44.6%)
2 (27.1%)
3 (11.3%)
4 (10.7%)
5 ( 6.2%)
1 (23.63%)
2 (16.42%)
3 (13.61%)
4 (13.13%)
5 ( 7.62%)
6 ( 6.54%)
7 ( 4.15%)
8 ( 3.97%)
9 ( 3.65%)
10 ( 1.83%)
11 ( 1.81%)
12 ( 1.08%)
13 ( 1.06%)
14 ( 0.27%)
15 ( 0.24%)
B
E
no phenotype
all others
no phenotype
all others
C
F
CLIPs
all others
CLIPs
all others

**Fig. S6. Distribution of non-phenotypic CLIPs in AgMelGCN2.0 and AgGCN1.0 networks.** Network communities in AgMelGCN2.0 (**A**) and AgGCN1.0 (**D**). Communities are sequentially numbered based on the node count and colored as indicated in the accompanying sub-panel. %, percentage of nodes within each community as compared to the entire network. Middle and bottom panels show the distribution of non-phenotypic CLIPs (blue, **B, E**) and all CLIPs (orange, **C, F**) in AgMelGCN2.0 (left) and AgGCN1.0 (right) networks, respectively. Network images generated in Gephi 0.10 using the ForceAtlas 2.0 algorithm with default settings.

**Table S1. dsRNA primer sequences.** *Added sequence for T7 promoter is underlined.

| Primer Name | Primer Sequences (5'-3')* | Primer design |
|---|---|---|
| dsGFP_F | TAATACGACTCACTATAGGGCGATGC | (*37*) |
| dsGFP_R | TAATACGACTCACTATAGGGCGGACT | |
| dsCACT_F | TAATACGACTCACTATAGGGTAACACTGCGCTTCATTTGG | (*51*) |
| dsCACT_R | TAATACGACTCACTATAGGGGCCCTTTTCAATGCTGATGT | |
| dsREL1_F | TAATACGACTCACTATAGGGGGGCAGTGGTCGTTGTGT | (*51*) |
| dsREL1_R | TAATACGACTCACTATAGGGGCACTGAATGCCCAAATTGT | |
| dsSRPN2_F | TAATACGACTCACTATAGGGCGAGGGCGCGGTCATTACG | (*5*) |
| dsSRPN2_R | TAATACGACTCACTATAGGGCAGCATTGTTCCGAGGGTTTCATC | |
| CLIPA1_eRNAi_f | TAATACGACTCACTATAGGGGGCTGGAATTGTAGCACAGA | This study |
| CLIPA1_eRNAi_r | TAATACGACTCACTATAGGGCGCACAACACTGCATCAGA | |
| dsCLIPA2_F | TAATACGACTCACTATAGGGATCCTAACAACGGCACACTG | (*47*) |
| dsCLIPA2_R | TAATACGACTCACTATAGGGTCCTGATCGCCATGATTGGT | |
| CLIPA3_eRNAi_f | TAATACGACTCACTATAGGGAAAATATCGGACTACACCTCCG | This study |
| CLIPA3_eRNAi_r | TAATACGACTCACTATAGGGGCAAGATGGCAATGTCGTT | |
| CLIPA4_eRNAi_f | TAATACGACTCACTATAGGGATGACCCCTTTCTCCAAAGC | This study |
| CLIPA4_eRNAi_r | TAATACGACTCACTATAGGGCTGGAAGCCGCTGTTGTC | |
| CLIPA5_eRNAi_f | TAATACGACTCACTATAGGGCTGAACCTGAATGCCCTCAC | This study |
| CLIPA5_eRNAi_r | TAATACGACTCACTATAGGGCGGTCACGCTAAAACCGAG | |
| CLIPA6_eRNAi_f | TAATACGACTCACTATAGGGGATCTCAGCCTGGACGATCT | This study |
| CLIPA6_eRNAi_r | TAATACGACTCACTATAGGGGCTGCTGTTCGAGCAGAGATA | |
| dsCLIPA7_F | TAATACGACTCACTATAGGGGTGCTGGCAGTCCTGGAACT | (*30*) |
| dsCLIPA7_R | TAATACGACTCACTATAGGGCCACCACCTTGGTTATATCC | |
| dsCLIPA8_F | TAATACGACTCACTATAGGGAACAACGAACCCGTAGAATA | (*26*) |
| dsCLIPA8_R | TAATACGACTCACTATAGGGGGTTAGCGCCTCGATACCCC | |
| CLIPA9_eRNAi_f | TAATACGACTCACTATAGGGGGACGGTGGCCATTTTTAAC | This study |
| CLIPA9_eRNAi_r | TAATACGACTCACTATAGGGGCGAATGTGCTCGGTGTAGA | |

| | | |
|---|---|---|
| CLIPA10_eRNAi_f | TAATACGACTCACTATAGGGGTGCTGACGAGTGGGGTT | This study |
| CLIPA10_eRNAi_r | TAATACGACTCACTATAGGGCTGTCGAGGATGAACTTCAGG | |
| CLIPA12_eRNAi_f | TAATACGACTCACTATAGGGCTGGTGGCACCGAATGTAG | This study |
| CLIPA12_eRNAi_r | TAATACGACTCACTATAGGGGCCACATCGAAGTGATGGT | |
| CLIPA13_eRNAi_f | TAATACGACTCACTATAGGGGAAGCGTTCAACAAGGGCT | This study |
| CLIPA13_eRNAi_r | TAATACGACTCACTATAGGGGCTTTGCCGAACAGATCCT | |
| dsCLIPA14_F | TAATACGACTCACTATAGGGCGGCATCATCGACATCCGTG | (*23*) |
| dsCLIPA14_R | TAATACGACTCACTATAGGGGGTTGCTGTCGGCGACACGC | |
| CLIPA15_eRNAi_f | TAATACGACTCACTATAGGGGCCTTACTACTGCTGCTCTTCG | This study |
| CLIPA15_eRNAi_r | TAATACGACTCACTATAGGGCAGCACCTCGTAGCAGATCA | |
| CLIPA19_eRNAi_f | TAATACGACTCACTATAGGGCTGTGTAAATCCTGCCGGTG | This study |
| CLIPA19_eRNAi_r | TAATACGACTCACTATAGGGAACACGTGCCGTTTATCTGG | |
| CLIPA26_eRNAi_f | TAATACGACTCACTATAGGGGGGTTCTTGCTGCTGCTACT | This study |
| CLIPA26_eRNAi_r | TAATACGACTCACTATAGGGGAGCTGGAACGATCGAAACT | |
| CLIPA27_eRNAi_f | TAATACGACTCACTATAGGGGACCGTCTTCAGTACCGGG | This study |
| CLIPA27_eRNAi_r | TAATACGACTCACTATAGGGGCAAGCACGTTCGGATCATA | |
| dsCLIPA28_F | TAATACGACTCACTATAGGGGCACCACCAAGGAACCGTTC | (*22*) |
| dsCLIPA28_R | TAATACGACTCACTATAGGGCCGCCGCAACCGATGCCCCA | |
| dsSPCLIP1_F | TAATACGACTCACTATAGGGGTCACCGAACACGTCCAACC | (*24*) |
| dsSPCLIP1_R | TAATACGACTCACTATAGGGATCGAAGCTGATCGGATCGG | |
| CLIPB1_eRNAi_f | TAATACGACTCACTATAGGGCAGGCGGGACGATTTCTACT | This study |
| CLIPB1_eRNAi_r | TAATACGACTCACTATAGGGTGTTCTCCTTCTTCAGCAGC | |
| CLIPB2_eRNAi_f | TAATACGACTCACTATAGGGAAACGCAGTTCCTCGCAA | This study |
| CLIPB2_eRNAi_r | TAATACGACTCACTATAGGGTCCATGGAAACTCTTCCAGC | |
| CLIPB3b_eRNAi_f | TAATACGACTCACTATAGGGAGTGCTCCGTGTATCGGGT | This study |
| CLIPB3b_eRNAi_r | TAATACGACTCACTATAGGGGGCTTTGCGTTAGTAGCAGTG | |
| dsCLIPB4_F | TAATACGACTCACTATAGGGTCAGGATTGCGTGAATCCGG | (*32*) |
| dsCLIPB4_R | TAATACGACTCACTATAGGGAGTTTTACGCTCGTAGAGAC | |
| CLIPB5_eRNAi_f | TAATACGACTCACTATAGGGCACTGAGGTTGCACCGTCTAC | This study |
| CLIPB5_eRNAi_r | TAATACGACTCACTATAGGGGGCATCTAGCAGCGGTTTT | |

| | | |
|---|---|---|
| CLIPB6_eRNAi_f | TAATACGACTCACTATAGGGGGTGGTGGTAGTGGTGGTACA | This study |
| CLIPB6_eRNAi_r | TAATACGACTCACTATAGGGCGGACATTGGGTAACCGTAA | |
| CLIPB7_eRNAi_f | TAATACGACTCACTATAGGGATTGACGGGGAGCAGATTT | This study |
| CLIPB7_eRNAi_r | TAATACGACTCACTATAGGGTTCCAGATTCTCCACGATCC | |
| dsCLIPB8_F | TAATACGACTCACTATAGGGTGTGCGACATCCCGAACGAG | (*31*) |
| dsCLIPB8_R | TAATACGACTCACTATAGGGCATCCGGTCCGAGATTGTCC | |
| dsCLIPB9_F | TAATACGACTCACTATAGGGAATGCACGACACCGACGAGG | (*31*) |
| dsCLIPB9_R | TAATACGACTCACTATAGGGTTTGCCCTCCTTGCGCTCAA | |
| dsCLIPB10_F | TAATACGACTCACTATAGGGCCGCAAAACTTTGGCATCGA | (*13*) |
| dsCLIPB10_R | TAATACGACTCACTATAGGGCTGGTAGGCGGGAAGTCATC | |
| CLIPB11_eRNAi_f | TAATACGACTCACTATAGGGCGAGTTCGATCTGAGCACAC | This study |
| CLIPB11_eRNAi_r | TAATACGACTCACTATAGGGCAGACAGATTGGCAGCACAT | |
| CLIPB12_eRNAi_f | TAATACGACTCACTATAGGGGGGTCTAACTCAAGGGCCAC | This study |
| CLIPB12_eRNAi_r | TAATACGACTCACTATAGGGCACAGCCTTTGAGGTCAAGC | |
| dsCLIPB13_F | TAATACGACTCACTATAGGGTACTACCGTCGCTCCGAGTA | (*22*) |
| dsCLIPB13_R | TAATACGACTCACTATAGGGTCGATGTCCGGACACAATGT | |
| dsCLIPB14_F | TAATACGACTCACTATAGGGATGTCGTACCAGCTGAAGCA | (*25*) |
| dsCLIPB14 _R | TAATACGACTCACTATAGGGCTACGCCAAAATTCCACGGA | |
| CLIPB15_eRNAi_f | TAATACGACTCACTATAGGGTGTGCTGTCCAAAGTTTAGC | This study |
| CLIPB15_eRNAi_r | TAATACGACTCACTATAGGGACGTATCGCTCGGAAATGAG | |
| CLIPB16_eRNAi_f | TAATACGACTCACTATAGGGCATCCGGACTACATCGAGGG | This study |
| CLIPB16_eRNAi_r | TAATACGACTCACTATAGGGCCCAGCCAATGATCTGTACC | |
| dsCLIPB17_F | TAATACGACTCACTATAGGGAGCGTGGGGAATTCCCGTGG | (*22*) |
| dsCLIPB17_R | TAATACGACTCACTATAGGGGGATCGTCCATCAGCAGCGA | |
| CLIPB18_eRNAi_f | TAATACGACTCACTATAGGGTGGGCGAGTTCGATATAAGC | This study |
| CLIPB18_eRNAi_r | TAATACGACTCACTATAGGGAGACAGATCGGCGACACATT | |
| CLIPB19_eRNAi_f | TAATACGACTCACTATAGGGTTGTGATAGTCGAAGCGGTG | This study |
| CLIPB19_eRNAi_r | TAATACGACTCACTATAGGGCGTTCGTGCATGCTACATCT | |
| CLIPB20_eRNAi_f | TAATACGACTCACTATAGGGCGTGTCCAGTGAATTCGGAG | This study |
| CLIPB20_eRNAi_r | TAATACGACTCACTATAGGGCCAATCGATCCACTGTCCAT | |

| | | |
|---|---|---|
| CLIPB36_eRNAi_f | TAATACGACTCACTATAGGGCTGCGTACCGGTGAAACAGT | This study |
| CLIPB36_eRNAi_r | TAATACGACTCACTATAGGGCACCGGGACTAGCGAACTCT | |
| CLIPB41_eRNAi_f | TAATACGACTCACTATAGGGCACACCTTGGGGCAGTTATT | This study |
| CLIPB41_eRNAi_r | TAATACGACTCACTATAGGGGTAGCTCGGCTTGGGATTG | |
| CLIPB44_eRNAi_f | TAATACGACTCACTATAGGGCTGGGGCCAAACACAGAAC | This study |
| CLIPB44_eRNAi_r | TAATACGACTCACTATAGGGCAGTTGTCGGACAGATTGGA | |
| CLIPB46_eRNAi_f | TAATACGACTCACTATAGGGCAATGTACCACATTTCGGGA | This study |
| CLIPB46_eRNAi_r | TAATACGACTCACTATAGGGCCCACGGAAACTGATCCA | |
| CLIPB47_eRNAi_f | TAATACGACTCACTATAGGGGCCATGATCGTGTACGTGTC | This study |
| CLIPB47_eRNAi_r | TAATACGACTCACTATAGGGCGTATCATACTCTCCGATTCGT | |
| CLIPC1_eRNAi_f | TAATACGACTCACTATAGGGCGGTATGCGTCATCGGTAG | This study |
| CLIPC1_eRNAi_r | TAATACGACTCACTATAGGGGGATGTCATCGATTACGGG | |
| CLIPC2_eRNAi_f | TAATACGACTCACTATAGGGCTAACCATTGATAACGCACGG | This study |
| CLIPC2_eRNAi_r | TAATACGACTCACTATAGGGAGCACACAACTGCTTGTTGG | |
| CLIPC3_eRNAi_f | TAATACGACTCACTATAGGGATGGGAAGCACATTCAAACG | This study |
| CLIPC3_eRNAi_r | TAATACGACTCACTATAGGGCGCGTTTGCACTTCGAGTAT | |
| CLIPC4_eRNAi_f | TAATACGACTCACTATAGGGGAGTTCAGCGGCAACGAT | This study |
| CLIPC4_eRNAi_r | TAATACGACTCACTATAGGGGGCCAGATATATGGTCGATGA | |
| CLIPC5_eRNAi_f | TAATACGACTCACTATAGGGGCACCATCGATCTGCTGTC | This study |
| CLIPC5_eRNAi_r | TAATACGACTCACTATAGGGACAGATCGGCTGCAGGAC | |
| CLIPC6_eRNAi_f | TAATACGACTCACTATAGGGCCCTCTGCCAGACGCTTTAC | This study |
| CLIPC6_eRNAi_r | TAATACGACTCACTATAGGGGTTCGTCTTGCAGCAGATCA | |
| CLIPC7_eRNAi_f | TAATACGACTCACTATAGGGTGTATCGGTGGGGTTGGTTA | This study |
| CLIPC7_eRNAi_r | TAATACGACTCACTATAGGGACACCGTCCGGAGATACTTC | |
| CLIPC9_eRNAi_f | TAATACGACTCACTATAGGGATGCAATGCGATCAACAAAA | This study |
| CLIPC9_eRNAi_r | TAATACGACTCACTATAGGGGCTCTCATGCAAATCAGTCTT | |
| CLIPC10_eRNAi_f | TAATACGACTCACTATAGGGCTCATCTCGTCCCGGTTTCT | This study |
| CLIPC10_eRNAi_r | TAATACGACTCACTATAGGGGCGATATCGTTCTGGTACGTG | |
| CLIPC12_eRNAi_f | TAATACGACTCACTATAGGGGCGCTCAATGTAATCGAACA | This study |
| CLIPC12_eRNAi_r | TAATACGACTCACTATAGGGACATTTGGGACCAAAGCTGA | |

| | | |
|---|---|---|
| CLIPC14_eRNAi_f | TAATACGACTCACTATAGGGCTACCACAACAACCACAGGG | This study |
| CLIPC14_eRNAi_r | TAATACGACTCACTATAGGGTGTGGTGCTGGATGTTTTCA | |
| CLIPD1_eRNAi_f | TAATACGACTCACTATAGGGGAGTGGCCCTGGATGGTAG | This study |
| CLIPD1_eRNAi_r | TAATACGACTCACTATAGGGCGTTTCGTTGAACTGCTTGA | |
| CLIPD2_eRNAi_f | TAATACGACTCACTATAGGGACGAATCGTTGGAGGTCACA | This study |
| CLIPD2_eRNAi_r | TAATACGACTCACTATAGGGGTGCCACGTCGTAGGAGC | |
| CLIPD3_eRNAi_f | TAATACGACTCACTATAGGGCAAGGTCAAGCGCAAGGAT | This study |
| CLIPD3_eRNAi_r | TAATACGACTCACTATAGGGGGACAGGTGCTCAGATCTTCA | |
| CLIPD4_eRNAi_f | TAATACGACTCACTATAGGGTTTCCTCGGAATTGGAAGTG | This study |
| CLIPD4_eRNAi_r | TAATACGACTCACTATAGGGCGTGCCGTAGTTGGAAAAGT | |
| CLIPD6_eRNAi_f | TAATACGACTCACTATAGGGCCGATCTTGCTATACCCCG | This study |
| CLIPD6_eRNAi_r | TAATACGACTCACTATAGGGACTGGGCTGCCGTTAAAGTT | |
| CLIPD7_eRNAi_f | TAATACGACTCACTATAGGGGGACACCTGCGTCTACCAGA | This study |
| CLIPD7_eRNAi_r | TAATACGACTCACTATAGGGCCATCGGTACCATCGGTC | |
| CLIPD8_eRNAi_f | TAATACGACTCACTATAGGGGCGTCGCACGTCATTTTT | This study |
| CLIPD8_eRNAi_r | TAATACGACTCACTATAGGGGCACGTGCGAAAAGTCATAC | |
| CLIPD9_eRNAi_f | TAATACGACTCACTATAGGGGACGTTCGGGATCAGGAAGA | This study |
| CLIPD9_eRNAi_r | TAATACGACTCACTATAGGGGACGGGCAGTATATGTTGTCG | |
| CLIPD11_eRNAi_f | TAATACGACTCACTATAGGGGGCAAGAATAAAAGGATTCCC | This study |
| CLIPD11_eRNAi_r | TAATACGACTCACTATAGGGCGCAGATGACAATCGTTGAG | |
| CLIPD12_eRNAi_f | TAATACGACTCACTATAGGGGGCCTGTTCACCAAGAACAA | This study |
| CLIPD12_eRNAi_r | TAATACGACTCACTATAGGGCGATGACACGTTTCACGTTCT | |
| CLIPD13_eRNAi_f | TAATACGACTCACTATAGGGCGACTACGTGATCAACTCGG | This study |
| CLIPD13_eRNAi_r | TAATACGACTCACTATAGGGGGGCATAAAGTGCACCGTC | |
| CLIPD14_eRNAi_f | TAATACGACTCACTATAGGGAGCTGGCGCTGGACTACAT | This study |
| CLIPD14_eRNAi_r | TAATACGACTCACTATAGGGGATGAAGGAATTCACCGGAC | |
| CLIPD20_eRNAi_f | TAATACGACTCACTATAGGGATGTCGTTGGACGAAATCTG | This study |
| CLIPD20_eRNAi_r | TAATACGACTCACTATAGGGGGTCGAAGCAGGAAAGGTGT | |
| CLIPD22_eRNAi_f | TAATACGACTCACTATAGGGTCCAGATTGTGGCCTCTCAT | This study |
| CLIPD22_eRNAi_r | TAATACGACTCACTATAGGGGGTAACGAATGCTGTTCTGC | |

| | | |
|---|---|---|
| CLIPE1_eRNAi_f | TAATACGACTCACTATAGGGGCGGCCAACTGTGTCTATG | This study |
| CLIPE1_eRNAi_r | TAATACGACTCACTATAGGGGATGTGCGTGTCGAACTTCA | |
| CLIPE4_eRNAi_f | TAATACGACTCACTATAGGGCTATTCGTCGGGTTCGTGTC | This study |
| CLIPE4_eRNAi_r | TAATACGACTCACTATAGGGGTTGCATCTCGATCATCGCT | |
| CLIPE5_eRNAi_f | TAATACGACTCACTATAGGGACAATGACATCATTCCCGAT | This study |
| CLIPE5_eRNAi_r | TAATACGACTCACTATAGGGGGCATTCGTTGTTGGAGTG | |
| CLIPE6_eRNAi_f | TAATACGACTCACTATAGGGTGCACTGAAGTTCCAGCAGA | This study |
| CLIPE6_eRNAi_r | TAATACGACTCACTATAGGGGGGACGAACCGTTTACCC | |
| CLIPE7_eRNAi_f | TAATACGACTCACTATAGGGGTGCAACCGATCTGCCTT | This study |
| CLIPE7_eRNAi_r | TAATACGACTCACTATAGGGCGAATGTCTGCAGAAACCAA | |
| CLIPE10_eRNAi_f | TAATACGACTCACTATAGGGGAGTTCGGCAACGCTTAGAA | This study |
| CLIPE10_eRNAi_r | TAATACGACTCACTATAGGGCGACGGAATGACTCGAAAC | |
| CLIPE12_eRNAi_f | TAATACGACTCACTATAGGGCGGCACTGCTATGGATCACT | This study |
| CLIPE12_eRNAi_r | TAATACGACTCACTATAGGGACTATTTCCAGCAGCGCAAC | |
| CLIPE13_eRNAi_f | TAATACGACTCACTATAGGGCTGATCGTGATCGCTAGTGC | This study |
| CLIPE13_eRNAi_r | TAATACGACTCACTATAGGGTCCTCCAAAAATAAAGAATCCG | |
| CLIPE15_eRNAi_f | TAATACGACTCACTATAGGGTCTGCGGTACTACAAATTCGG | This study |
| CLIPE15_eRNAi_r | TAATACGACTCACTATAGGGCGCCGTTAGGACATAGTTCTC | |
| CLIPE17_eRNAi_f | TAATACGACTCACTATAGGGGCCAATGTGCTTCCCATAAA | This study |
| CLIPE17_eRNAi_r | TAATACGACTCACTATAGGGACTTCTTCCCTCCGATTGGT | |
| CLIPE18_eRNAi_f | TAATACGACTCACTATAGGGCGTTTGTGGTGCGGAACT | This study |
| CLIPE18_eRNAi_r | TAATACGACTCACTATAGGGGGGAACTCGGTCACATTCAC | |
| CLIPE19_eRNAi_f | TAATACGACTCACTATAGGGCGTTCTTACAGCAGCACACTG | This study |
| CLIPE19_eRNAi_r | TAATACGACTCACTATAGGGACACGCCGGTATAACTCCAA | |
| CLIPE21_eRNAi_f | TAATACGACTCACTATAGGGCGACGCATGTGCTGGTATT | This study |
| CLIPE21_eRNAi_r | TAATACGACTCACTATAGGGGGTGAAACCAGGCAGTTGAT | |
| CLIPE22_eRNAi_f | TAATACGACTCACTATAGGGGATGGCCGCATGAAGTTAGT | This study |
| CLIPE22_eRNAi_r | TAATACGACTCACTATAGGGTACGGTGAAATGGAGGAATTG | |
| CLIPE25_eRNAi_f | TAATACGACTCACTATAGGGTGCAATTTACGATACACCCTTC | This study |
| CLIPE25_eRNAi_r | TAATACGACTCACTATAGGGATATCAGCGACCCTCCACAG | |

| | | |
|---|---|---|
| CLIPE30_eRNAi_f | TAATACGACTCACTATAGGGTCGCTAGATGGAATCATCGC | This study |
| CLIPE30_eRNAi_r | TAATACGACTCACTATAGGGTCCACCACTCAAATCCATTG | This study |
| MODSP1_ eRNAi_f | TAATACGACTCACTATAGGGGCGTCGGCTACCAGAAACCG | This study |
| MODSP1_ eRNAi_r | TAATACGACTCACTATAGGGGTGTCGTCGTGCAGGTTTTT | This study |
| TEP1 _f | TAATACGACTCACTATAGGGTTTGTGGGCCTTAAAGCGCTG | (*24*) |
| TEP1 _r | TAATACGACTCACTATAGGGACCACGTAACCGCTCGGTAAG | (*24*) |
| SPZ1 _f | TAATACGACTCACTATAGGGCTTCCGAAAGGACTTTGGCA | This study |
| SPZ1 _r | TAATACGACTCACTATAGGGCGGTGGACTGCTGCTCCTGT | This study |
| SPZ3 _f | TAATACGACTCACTATAGGGGGTGACTCCGATTGTTGGGA | This study |
| SPZ3_r | TAATACGACTCACTATAGGGGGGCAGAACAGTTGATGGTACT | This study |
| T7 | TAATACGACTCACTATAGGG | (*31*) |

**Table S2. RT-qPCR primers, primer efficiency and kd efficiency.** *RT-qPCR was used to measure the relative expression levels with ribosomal protein S7 as the internal reference and ds*GFP*-treated samples as the calibrator condition. R calculated using the Pfaffl **(*52*)** mathematical model for relative quantification.

| Target name | KD efficiency | Primer name | Primer Sequences (5'-3') | Primer design | Slope ± SE | $R^2$ | *P* | Primer efficiency | | Relative expression ratio (R)* ± SE |
|---|---|---|---|---|---|---|---|---|---|---|
| RpS7 | not applicable | qS7_F | GTGCGCGAGTTGGAGAAGA | (*5*) | -3.107 ± 0.08 | 0.987 | <0.0001 | 2.1 | 109.80% | |
| | | qS7_R | ATCGGTTTGGGCAGAATGC | | | | | | | |
| CACT | 49.1% (*34*) | qCACT_F | AATCTGGGCCTGATGGACA | (*51*) | | | | | | |
| | | qCACT_R | ACTGCCAGGTGCAGTTGAGT | | | | | | | |
| REL1 | 57.2% (*34*) | qREL1_F | TCAACAGATGCCAAAAGAGGAAAT | (*51*) | | | | | | |
| | | qREL1_R | CTGGTTGGAGGGATTGTG | | | | | | | |
| SRPN2 | 50% (*5*) | qSRPN2_F | TGCCGTGTCCAACACCAA | (*5*) | | | | | | |
| | | qSRPN2_R | CGCGTATGGTCGATGTTATCG | | | | | | | |
| CLIPA2 | 96.3% | qCLIPA2_F | GATACTACCTGCACGGGTTGGT | (*29*) | -3.141 ± 0.08 | 0.988 | <0.0001 | 2.08 | 108.10% | 0.037 ± 0.001 |
| | | qCLIPA2_R | CAGTATAAGGTATCTGCTTCTGATGGC | | | | | | | |
| CLIPA3 | 92.92% | qCLIPA3_f | CGTTGGTGTGCGCGCTAAAC | This study | -3.411 ± 0.12 | 0.978 | <0.0001 | 1.96 | 96.40% | 0.071 ± 0.011 |
| | | qCLIPA3_r | CGTTTGCACCGCAACCGATG | | | | | | | |
| CLIPA4 | 94.49% | qCLIPA4_f | TGATTCTGTTCCGGTCGTGGATC | This study | -3.432 ± 0.08 | 0.988 | <0.0001 | 1.96 | 95.60% | 0.055 ± 0.007 |
| | | qCLIPA4_r | TCGCATCGTACACGGTCGTGT | | | | | | | |
| CLIPA5 | 83.60% | qCLIPA5_f | TACATGCATCAGAACCGTCG | This study | -3.144 ± 0.08 | 0.987 | <0.0001 | 2.08 | 108.00% | 0.164 ± 0.023 |
| | | qCLIPA5_r | CATCGTTCGCCAATGATTCG | | | | | | | |
| CLIPA7 | 86.95% | qCLIPA7_F | GTACGGTGAGTTCCCGTGGAT | (*29*) | -3.197 ± 0.11 | 0.977 | <0.0001 | 2.05 | 105.50% | 0.131 ± 0.018 |
| | | qCLIPA7_R | ACGTTGATCACTTGGTCCAGC | | | | | | | |

| | | | | | | | | | | |
|---|---|---|---|---|---|---|---|---|---|---|
| CLIPA8 | 82.11% | qCLIPA8_f | TACGTTCTGGCGG GAATTGT | This study | -3.337 ± 0.12 | 0.975 | <0.0001 | 1.99 | 99.40% | 0.179 ± 0.007 |
| | | qCLIPA8_r | CACATTCACGTAA GCACCGG | | | | | | | |
| CLIPA13 | 78.1% | qCLIPA13_f | ACTCACTACTATC AGGCGGG | This study | -3.142 ± 0.07 | 0.990 | <0.0001 | 2.08 | 108.10% | 0.219 ± 0.054 |
| | | qCLIPA13_r | CACGGAAGACTGA CACACTG | | | | | | | |
| CLIPA14 | 81.5% | qCLIPA14_f | TCGACGACCTCTA CCTCTCG | This study | -3.366 ± 0.13 | 0.975 | <0.0001 | 1.98 | 98.20% | 0.185 ± 0.063 |
| | | qCLIPA14_r | CGGTTCCGTGACA GTGTTTT | | | | | | | |
| CLIPA19 | 89.27% | qCLIPA19_f | GCAGTGCTGATGC TGGAGGT | This study | -3.452 ± 0.08 | 0.989 | <0.0001 | 1.95 | 94.80% | 0.107 ± 0.051 |
| | | qCLIPA19_r | TCGCCCGATCCGC ACTTTTG | | | | | | | |
| CLIPA28 | 97.35% | qCLIPA28_F | ATAAAGCATGCCC AAACCAC | (*27*) | -3.171 ± 0.08 | 0.989 | <0.0001 | 2.07 | 106.70% | 0.026 ± 0.009 |
| | | qCLIPA28_R | CTGACAGCACACC AGCAGAT | | | | | | | |
| SPCLIP1 | 88.29% | qSPCLIP1_F | CTTGCTGAACGAC GACGATA | (*27*) | -3.147 ± 0.08 | 0.987 | <0.0001 | 2.08 | 107.90% | 0.117 ± 0.003 |
| | | qSPCLIP1_R | TCCTGTTCCTCCT CGTCACT | | | | | | | |
| CLIPB4 | 82.55% | qCLIPB4_f | AGGAGGATGACTT CTACG | This study | -3.158 ± 0.08 | 0.988 | <0.0001 | 2.07 | 107.30% | 0.175 ± 0.017 |
| | | qCLIPB4_r | GTTATGGTGGCTC TTGTC | | | | | | | |
| CLIPB8 | 90% (*31*) | qCLIPB8_F | CATGAGCTACGAT ATGAAGC | (*31*) | | | | | | |
| | | qCLIPB8_R | TTGATCCACGGCA GATAG | | | | | | | |
| CLIPB9 | 65% (*37*) | qCLIPB9_F | GAAGCAATATCAA GGAGTA | (*37*) | | | | | | |
| | | qCLIPB9_R | AGTATCGTAGAAT ATAGCATTA | | | | | | | |
| CLIPB10 | 95% (*13*) | qCLIPB10_F | TTGATGGTAAAGC GGTTT | (*13*) | | | | | | |
| | | qCLIPB10_R | AGATACGACGACA CTCTC | | | | | | | |
| CLIPB12 | 74.48% | qCLIPB12_f | GTTGGCTACAGAG GACGGTT | This study | -3.118 ± 0.11 | 0.979 | <0.0001 | 2.09 | 109.30% | 0.255 ± 0.062 |
| | | qCLIPB12_r | AGCTGTTAGTGAT GCACCGT | | | | | | | |

| | | | | | | | | | | | | | | |
|---|---|---|---|---|---|---|---|---|---|---|---|---|---|---|
| CLIPB13 | 93.12% | qCLIPB13_f | GTGGATGTGGCTGGGGCAAA | This study | -3.192 | ± | 0.1 | 0.984 | <0.0001 | 2.06 | 105.70% | 0.069 | ± | 0.018 |
| | | qCLIPB13_r | TGTAAATGCCGGGCACGCTT | | | | | | | | | | | |
| CLIPB14 | 97.43% | qCLIPB14_f | GGCGTACACGTGAACGATAT | This study | -3.210 | ± | 0.05 | 0.995 | <0.0001 | 2.05 | 104.90% | 0.026 | ± | 0.006 |
| | | qCLIPB14_r | AAACGGGCAAACATATCGGT | | | | | | | | | | | |
| CLIPB17 | 74.39% | qCLIPB17_f | TCCTGGACAATATGGCGTTG | This study | -3.341 | ± | 0.12 | 0.976 | <0.0001 | 1.99 | 99.20% | 0.256 | ± | 0.008 |
| | | qCLIPB17_r | TTTGAGACCGGAAACCACTG | | | | | | | | | | | |
| CLIPB20 | 65.79% | qCLIPB20_f | ACACACTTAGCTGCTTCGGT | This study | -3.246 | ± | 0.1 | 0.983 | <0.0001 | 2.03 | 103.30% | 0.342 | ± | 0.063 |
| | | qCLIPB20_r | CCAGCGAGATACGTTCCAGT | | | | | | | | | | | |
| CLIPC2 | 95.02% | qCLIPC2_F | CTCGTTCGGTATTGGCTGTG | (*27*) | -3.171 | ± | 0.09 | 0.985 | <0.0001 | 2.07 | 106.70% | 0.050 | ± | 0.005 |
| | | qCLIPC2_R | TGCCCTCGATCCAGTCTATG | | | | | | | | | | | |
| CLIPC3 | 65.96% | qCLIPC3_f | ACACTACCCTCCATTGTGCG | This study | -3.519 | ± | 0.12 | 0.977 | <0.0001 | 1.92 | 92.40% | 0.340 | ± | 0.016 |
| | | qCLIPC3_r | CGATGAAGCGCAAGATGTCG | | | | | | | | | | | |
| CLIPC4 | 90.10% | qCLIPC4_F | ATTGGACCGTCGATCAACCT | (*27*) | -3.354 | ± | 0.06 | 0.995 | <0.0001 | 1.99 | 98.70% | 0.099 | ± | 0.014 |
| | | qCLIPC4_R | GTGCCCTTCATCAGCTTGTC | | | | | | | | | | | |
| CLIPC9 | 74.48% | qCLIPC9_f | ATTGAAGGCATCCCCGAACA | This study | -3.210 | ± | 0.09 | 0.984 | <0.0001 | 2.04 | 104.90% | 0.255 | ± | 0.033 |
| | | qCLIPC9_r | CCCTGCGCACAGTACATTTC | | | | | | | | | | | |
| CLIPC12 | 75.56% | qCLIPC12_f | GATGCGGGTGTCAGAGTACA | This study | -3.181 | ± | 0.07 | 0.991 | <0.0001 | 2.06 | 106.20% | 0.244 | ± | 0.015 |
| | | qCLIPC12_r | CTCCCACCATCGATTCGCTT | | | | | | | | | | | |

| | | | | | | | | | | |
|---|---|---|---|---|---|---|---|---|---|---|
| CLIPC14 | 72.35% | qCLIPC14_f | GTATGTCGACGCG CGTGTCA | This study | -3.167 ± 0.11 | 0.979 | <0.0001 | 2.07 | 106.90% | 0.277 ± 0.025 |
| | | qCLIPC14_r | ACCACGCACACTC CAGTCCT | | | | | | | |
| CLIPD3 | 96.70% | qCLIPD3_f | GTTTTCTGAGCCTT CCTGCG | This study | -3.142 ± 0.06 | 0.986 | <0.0001 | 2.08 | 108.10% | 0.033 ± 0.008 |
| | | qCLIPD3_r | GAACTTTCCGCCTC GTTCAC | | | | | | | |
| MODSP1 | 62.14% | qMODSP1_f | GTGCGACAACAAC AAGTACG | This study | -3.131 ± 0.08 | 0.988 | <0.0001 | 2.09 | 108.60% | 0.379 ± 0.055 |
| | | qMODSP1_r | CGCGTCCAGTATCA TTGGAA | | | | | | | |
| TEP1 | 65.92 | qTEP1_F | AAAGCTGTTGCGTC AGGG | (*24*) | -3.317 ± 0.08 | 0.990 | <0.0001 | 2 | 100.20% | 0.341 ± 0.032 |
| | | qTEP1_R | TTCTCCCACACACC AAACGAA | | | | | | | |
| SPZ1 | 80.14 | qSPZ1_f | CCCACTCACGGAA ACAATGA | This study | -3.125 ± 0.09 | 0.985 | <0.0001 | 2.09 | 108.90% | 0.199 ± 0.008 |
| | | qSPZ1_r | TTATTTTCGTTGCC CGTTGC | | | | | | | |
| SPZ3 | 79.40 | qSPZ3_f | GGCTGTGAGCAGA AGTACAA | This study | -3.108 ± 0.08 | 0.988 | <0.0001 | 2.1 | 109.80% | 0.206 ± 0.021 |
| | | qSPZ3_r | ATGCCCTTACAGTC GTTGTC | | | | | | | |

**Table S3: MelASA RNAi screen: Total melanotic spot area of genes tested as positive regulators.** DsRNA injected mosquitoes were challenged with 50.6 nL of resuspended lyophilized *Micrococcus luteus* at $OD_{600}$ = 5. Batches of 7-10 candidate genes were tested in independent groups using their own positive (ds*CLIPB4*) and negative controls. Three independent biological replicates were performed using different mosquito generations.

| CLIP family | sub family | Gene name | AGAP | Assay type | Tested as | ds*GFP* control (individual replicates) | | | | | Positive control-ds*CLIPB4* (individual replicates) | | | | | ds*GOI* (individual replicates) | | | | | Phenotype |
|---|---|---|---|---|---|---|---|---|---|---|---|---|---|---|---|---|---|---|---|---|---|
| | | | | | | 1 | 2 | 3 | Mean | 95% CI of mean | 1 | 2 | 3 | Mean | 95% CI of mean | 1 | 2 | 3 | Mean | 95% CI of mean | |
| | | TEP1 | AGAP010815 | MelASA | positive | 0.184 | 0.202 | 0.198 | 0.195 | 0.171 - 0.218 | 0.139 | 0.128 | 0.133 | 0.133 | 0.120 - 0.147 | 0.071 | 0.082 | 0.064 | 0.072 | 0.050 - 0.095 | yes |
| | | MODSP1 | AGAP001798 | MelASA | positive | 0.194 | 0.204 | 0.197 | 0.198 | 0.186 - 0.211 | 0.117 | 0.122 | 0.126 | 0.122 | 0.111 - 0.133 | 0.145 | 0.153 | 0.139 | 0.146 | 0.128 - 0.163 | yes |
| cSPH | CLIPA | CLIPA1 | AGAP011791 | MelASA | positive | 0.202 | 0.207 | 0.189 | 0.199 | 0.176 - 0.222 | 0.124 | 0.106 | 0.131 | 0.120 | 0.088 - 0.151 | 0.202 | 0.182 | 0.201 | 0.195 | 0.167 - 0.223 | no |
| cSPH | CLIPA | CLIPA2 | AGAP011790 | MelASA | positive | 0.194 | 0.203 | 0.194 | 0.197 | 0.188 - 0.210 | 0.137 | 0.136 | 0.104 | 0.126 | 0.079 - 0.172 | 0.223 | 0.178 | 0.189 | 0.197 | 0.138 - 0.255 | no |
| cSPH | CLIPA | CLIPA3 | AGAP012591 | MelASA | positive | 0.194 | 0.203 | 0.194 | 0.197 | 0.188 - 0.210 | 0.137 | 0.136 | 0.104 | 0.126 | 0.079 - 0.172 | 0.197 | 0.199 | 0.182 | 0.193 | 0.170 - 0.216 | no |
| cSPH | CLIPA | CLIPA4 | AGAP011780 | MelASA | positive | 0.194 | 0.203 | 0.194 | 0.197 | 0.188 - 0.210 | 0.137 | 0.136 | 0.104 | 0.126 | 0.079 - 0.172 | 0.206 | 0.202 | 0.174 | 0.194 | 0.151 - 0.237 | no |
| cSPH | CLIPA | CLIPA5 | AGAP011787 | MelASA | positive | 0.194 | 0.203 | 0.194 | 0.197 | 0.188 - 0.210 | 0.137 | 0.136 | 0.104 | 0.126 | 0.079 - 0.172 | 0.197 | 0.194 | 0.201 | 0.197 | 0.189 - 0.206 | no |
| cSPH | CLIPA | CLIPA6 | AGAP011789 | MelASA | positive | 0.194 | 0.203 | 0.194 | 0.197 | 0.188 - 0.210 | 0.137 | 0.136 | 0.104 | 0.126 | 0.079 - 0.172 | 0.214 | 0.183 | 0.218 | 0.205 | 0.157 - 0.253 | no |
| cSPH | CLIPA | CLIPA7 | AGAP011792 | MelASA | positive | 0.194 | 0.203 | 0.194 | 0.197 | 0.188 - 0.210 | 0.137 | 0.136 | 0.104 | 0.126 | 0.079 - 0.172 | 0.186 | 0.201 | 0.183 | 0.190 | 0.166 - 0.214 | no |
| cSPH | CLIPA | CLIPA8 | AGAP010731 | MelASA | positive | 0.194 | 0.203 | 0.194 | 0.197 | 0.188 - 0.210 | 0.137 | 0.136 | 0.104 | 0.126 | 0.079 - 0.172 | 0.057 | 0.079 | 0.084 | 0.073 | 0.038 - 0.109 | yes |
| cSPH | CLIPA | CLIPA9 | AGAP010968 | MelASA | positive | 0.194 | 0.203 | 0.194 | 0.197 | 0.188 - 0.210 | 0.137 | 0.136 | 0.104 | 0.126 | 0.079 - 0.172 | 0.192 | 0.209 | 0.213 | 0.205 | 0.177 - 0.232 | no |
| cSPH | CLIPA | CLIPA10 | AGAP006954 | MelASA | positive | 0.194 | 0.203 | 0.194 | 0.197 | 0.188 - 0.210 | 0.137 | 0.136 | 0.104 | 0.126 | 0.079 - 0.172 | 0.181 | 0.198 | 0.197 | 0.192 | 0.168 - 0.216 | no |
| cSPH | CLIPA | CLIPA12 | AGAP011781 | MelASA | positive | 0.194 | 0.203 | 0.194 | 0.197 | 0.188 - 0.210 | 0.137 | 0.136 | 0.104 | 0.126 | 0.079 - 0.172 | 0.186 | 0.189 | 0.213 | 0.196 | 0.159 - 0.233 | no |
| cSPH | CLIPA | CLIPA13 | AGAP011783 | MelASA | positive | 0.193 | 0.199 | 0.191 | 0.194 | 0.184 - 0.205 | 0.125 | 0.132 | 0.140 | 0.132 | 0.114 - 0.151 | 0.175 | 0.197 | 0.185 | 0.186 | 0.158 - 0.213 | no |
| cSPH | CLIPA | CLIPA14 | AGAP011788 | MelASA | positive | 0.193 | 0.199 | 0.191 | 0.194 | 0.184 - 0.205 | 0.125 | 0.132 | 0.140 | 0.132 | 0.114 - 0.151 | 0.195 | 0.209 | 0.208 | 0.204 | 0.185 - 0.223 | no |
| cSPH | CLIPA | CLIPA15 | AGAP002815 | MelASA | positive | 0.193 | 0.199 | 0.191 | 0.194 | 0.184 - 0.205 | 0.125 | 0.132 | 0.140 | 0.132 | 0.114 - 0.151 | 0.234 | 0.184 | 0.204 | 0.207 | 0.145 - 0.270 | no |
| cSPH | CLIPA | CLIPA19 | AGAP003245 | MelASA | positive | 0.193 | 0.199 | 0.191 | 0.194 | 0.184 - 0.205 | 0.125 | 0.132 | 0.140 | 0.132 | 0.114 - 0.151 | 0.194 | 0.202 | 0.189 | 0.195 | 0.179 - 0.211 | no |
| cSPH | CLIPA | CLIPA26 | AGAP001964 | MelASA | positive | 0.193 | 0.199 | 0.191 | 0.194 | 0.184 - 0.205 | 0.125 | 0.132 | 0.140 | 0.132 | 0.114 - 0.151 | 0.207 | 0.189 | 0.197 | 0.198 | 0.175 - 0.220 | no |
| cSPH | CLIPA | CLIPA27 | AGAP000290 | MelASA | positive | 0.193 | 0.199 | 0.191 | 0.194 | 0.184 - 0.205 | 0.125 | 0.132 | 0.140 | 0.132 | 0.114 - 0.151 | 0.193 | 0.197 | 0.193 | 0.194 | 0.187 - 0.201 | no |
| cSPH | CLIPA | CLIPA28 | AGAP010730 | MelASA | positive | 0.193 | 0.199 | 0.191 | 0.194 | 0.184 - 0.205 | 0.125 | 0.132 | 0.140 | 0.132 | 0.114 - 0.151 | 0.119 | 0.117 | 0.125 | 0.120 | 0.110 - 0.131 | yes |
| cSPH | CLIPA | SPCLIP1 | AGAP028725 | MelASA | positive | 0.193 | 0.199 | 0.191 | 0.194 | 0.184 - 0.205 | 0.125 | 0.132 | 0.140 | 0.132 | 0.114 - 0.151 | 0.059 | 0.057 | 0.048 | 0.055 | 0.040 - 0.069 | yes |
| cSP | CLIPB | CLIPB1 | AGAP003251 | MelASA | positive | 0.205 | 0.217 | 0.223 | 0.215 | 0.192 - 0.238 | 0.122 | 0.119 | 0.135 | 0.125 | 0.104 - 0.147 | 0.187 | 0.213 | 0.204 | 0.201 | 0.169 - 0.234 | no |
| cSP | CLIPB | CLIPB2 | AGAP003246 | MelASA | positive | 0.205 | 0.217 | 0.223 | 0.215 | 0.192 - 0.238 | 0.122 | 0.119 | 0.135 | 0.125 | 0.104 - 0.147 | 0.201 | 0.200 | 0.211 | 0.204 | 0.189 - 0.219 | no |
| cSP | CLIPB | CLIPB3b | AGAP003249 | MelASA | positive | 0.205 | 0.217 | 0.223 | 0.215 | 0.192 - 0.238 | 0.122 | 0.119 | 0.135 | 0.125 | 0.104 - 0.147 | 0.191 | 0.202 | 0.200 | 0.198 | 0.183 - 0.212 | no |
| cSP | CLIPB | CLIPB5 | AGAP004148 | MelASA | positive | 0.205 | 0.217 | 0.223 | 0.215 | 0.192 - 0.238 | 0.122 | 0.119 | 0.135 | 0.125 | 0.104 - 0.147 | 0.195 | 0.208 | 0.192 | 0.198 | 0.177 - 0.220 | no |
| cSP | CLIPB | CLIPB6 | AGAP003252 | MelASA | positive | 0.205 | 0.217 | 0.223 | 0.215 | 0.192 - 0.238 | 0.122 | 0.119 | 0.135 | 0.125 | 0.104 - 0.147 | 0.194 | 0.197 | 0.186 | 0.192 | 0.178 - 0.207 | no |
| cSPH | CLIPB | CLIPB7 | AGAP002270 | MelASA | positive | 0.205 | 0.217 | 0.223 | 0.215 | 0.192 - 0.238 | 0.122 | 0.119 | 0.135 | 0.125 | 0.104 - 0.147 | 0.208 | 0.195 | 0.182 | 0.195 | 0.163 - 0.227 | no |
| cSP | CLIPB | CLIPB8 | AGAP003057 | MelASA | positive | 0.194 | 0.204 | 0.197 | 0.198 | 0.186 - 0.211 | 0.117 | 0.122 | 0.126 | 0.122 | 0.111 - 0.133 | 0.135 | 0.158 | 0.132 | 0.142 | 0.167 - 0.220 | yes |
| cSP | CLIPB | CLIPB9 | AGAP029769 | MelASA | positive | 0.194 | 0.204 | 0.197 | 0.198 | 0.186 - 0.211 | 0.117 | 0.122 | 0.126 | 0.122 | 0.111 - 0.133 | 0.126 | 0.118 | 0.125 | 0.123 | 0.106 - 0.177 | yes |
| cSP | CLIPB | CLIPB10 | AGAP029770 | MelASA | positive | 0.194 | 0.204 | 0.197 | 0.198 | 0.186 - 0.211 | 0.117 | 0.122 | 0.126 | 0.122 | 0.111 - 0.133 | 0.101 | 0.110 | 0.105 | 0.105 | 0.112 - 0.134 | yes |
| cSP | CLIPB | CLIPB11 | AGAP009214 | MelASA | positive | 0.205 | 0.217 | 0.223 | 0.215 | 0.192 - 0.238 | 0.122 | 0.119 | 0.135 | 0.125 | 0.104 - 0.147 | 0.205 | 0.191 | 0.184 | 0.193 | 0.094 - 0.117 | no |
| cSPH | CLIPB | CLIPB12 | AGAP009217 | MelASA | positive | 0.217 | 0.208 | 0.195 | 0.207 | 0.179 - 0.234 | 0.111 | 0.124 | 0.119 | 0.118 | 0.102 - 0.134 | 0.219 | 0.220 | 0.217 | 0.219 | 0.215 - 0.223 | no |
| cSP | CLIPB | CLIPB13 | AGAP004855 | MelASA | positive | 0.202 | 0.227 | 0.218 | 0.216 | 0.184 - 0.247 | 0.110 | 0.136 | 0.125 | 0.124 | 0.091 - 0.156 | 0.128 | 0.117 | 0.123 | 0.123 | 0.109 - 0.136 | yes |
| cSP | CLIPB | CLIPB14 | AGAP010833 | MelASA | positive | 0.202 | 0.227 | 0.218 | 0.216 | 0.184 - 0.247 | 0.110 | 0.136 | 0.125 | 0.124 | 0.091 - 0.156 | 0.198 | 0.185 | 0.206 | 0.196 | 0.170 - 0.223 | no |
| cSP | CLIPB | CLIPB15 | AGAP009844 | MelASA | positive | 0.202 | 0.227 | 0.218 | 0.216 | 0.184 - 0.247 | 0.110 | 0.136 | 0.125 | 0.124 | 0.091 - 0.156 | 0.193 | 0.185 | 0.191 | 0.190 | 0.179 - 0.200 | no |
| cSPH | CLIPB | CLIPB16 | AGAP009263 | MelASA | positive | 0.202 | 0.227 | 0.218 | 0.216 | 0.184 - 0.247 | 0.110 | 0.136 | 0.125 | 0.124 | 0.091 - 0.156 | 0.184 | 0.172 | 0.213 | 0.190 | 0.137 - 0.242 | no |
| cSP | CLIPB | CLIPB17 | AGAP001648 | MelASA | positive | 0.191 | 0.182 | 0.191 | 0.188 | 0.175 - 0.201 | 0.124 | 0.126 | 0.114 | 0.121 | 0.105 - 0.137 | 0.128 | 0.124 | 0.136 | 0.129 | 0.114 - 0.145 | yes |
| cSP | CLIPB | CLIPB18 | AGAP009215 | MelASA | positive | 0.191 | 0.182 | 0.191 | 0.188 | 0.175 - 0.201 | 0.124 | 0.126 | 0.114 | 0.121 | 0.105 - 0.137 | 0.183 | 0.180 | 0.193 | 0.185 | 0.168 - 0.202 | no |
| cSP | CLIPB | CLIPB19 | AGAP003247 | MelASA | positive | 0.202 | 0.227 | 0.218 | 0.216 | 0.184 - 0.247 | 0.110 | 0.136 | 0.125 | 0.124 | 0.091 - 0.156 | 0.191 | 0.188 | 0.179 | 0.186 | 0.171 - 0.202 | no |
| cSP | CLIPB | CLIPB20 | AGAP012037 | MelASA | positive | 0.205 | 0.217 | 0.223 | 0.215 | 0.192 - 0.238 | 0.122 | 0.119 | 0.135 | 0.125 | 0.104 - 0.147 | 0.148 | 0.159 | 0.136 | 0.148 | 0.119 - 0.176 | yes |

| | | | | | | | | | | | | | | | | | | | | | |
|---|---|---|---|---|---|---|---|---|---|---|---|---|---|---|---|---|---|---|---|---|---|
| cSPH | CLIPB | CLIPB36 | AGAP013184 | MelASA | positive | 0.202 | 0.227 | 0.218 | 0.216 | 0.184 - 0.247 | 0.110 | 0.136 | 0.125 | 0.124 | 0.091 - 0.156 | 0.222 | 0.228 | 0.204 | 0.218 | 0.187 - 0.249 | no |
| cSP | CLIPB | CLIPB41 | AGAP004149 | MelASA | positive | 0.202 | 0.227 | 0.218 | 0.216 | 0.184 - 0.247 | 0.110 | 0.136 | 0.125 | 0.124 | 0.091 - 0.156 | 0.200 | 0.195 | 0.187 | 0.194 | 0.178 - 0.210 | no |
| cSP | CLIPB | CLIPB44 | AGAP009220 | MelASA | positive | 0.202 | 0.227 | 0.218 | 0.216 | 0.184 - 0.247 | 0.110 | 0.136 | 0.125 | 0.124 | 0.091 - 0.156 | 0.188 | 0.192 | 0.216 | 0.199 | 0.161 - 0.236 | no |
| cSP | CLIPB | CLIPB46 | AGAP009849 | MelASA | positive | 0.202 | 0.227 | 0.218 | 0.216 | 0.184 - 0.247 | 0.110 | 0.136 | 0.125 | 0.124 | 0.091 - 0.156 | 0.194 | 0.208 | 0.201 | 0.201 | 0.184 - 0.218 | no |
| cSP | CLIPB | CLIPB47 | AGAP003686 | MelASA | positive | 0.202 | 0.227 | 0.218 | 0.216 | 0.184 - 0.247 | 0.110 | 0.136 | 0.125 | 0.124 | 0.091 - 0.156 | 0.188 | 0.210 | 0.205 | 0.201 | 0.172 - 0.230 | no |
| cSP | CLIPC | CLIPC1 | AGAP008835 | MelASA | positive | 0.194 | 0.204 | 0.197 | 0.198 | 0.186 - 0.211 | 0.117 | 0.122 | 0.126 | 0.122 | 0.111 - 0.133 | 0.216 | 0.209 | 0.215 | 0.213 | 0.204 - 0.223 | no |
| cSP | CLIPC | CLIPC2 | AGAP004317 | MelASA | positive | 0.217 | 0.208 | 0.195 | 0.207 | 0.179 - 0.234 | 0.111 | 0.124 | 0.119 | 0.118 | 0.102 - 0.134 | 0.204 | 0.193 | 0.217 | 0.205 | 0.175 - 0.235 | no |
| cSP | CLIPC | CLIPC3 | AGAP004318 | MelASA | positive | 0.205 | 0.217 | 0.223 | 0.215 | 0.192 - 0.238 | 0.122 | 0.119 | 0.135 | 0.125 | 0.104 - 0.147 | 0.125 | 0.133 | 0.142 | 0.133 | 0.112 - 0.155 | yes |
| cSP | CLIPC | CLIPC4 | AGAP000573 | MelASA | positive | 0.217 | 0.208 | 0.195 | 0.207 | 0.179 - 0.234 | 0.111 | 0.124 | 0.119 | 0.118 | 0.102 - 0.134 | 0.182 | 0.197 | 0.218 | 0.199 | 0.154 - 0.244 | no |
| cSP | CLIPC | CLIPC5 | AGAP000571 | MelASA | positive | 0.217 | 0.208 | 0.195 | 0.207 | 0.179 - 0.234 | 0.111 | 0.124 | 0.119 | 0.118 | 0.102 - 0.134 | 0.204 | 0.189 | 0.181 | 0.191 | 0.162 - 0.220 | no |
| cSP | CLIPC | CLIPC6 | AGAP000315 | MelASA | positive | 0.217 | 0.208 | 0.195 | 0.207 | 0.179 - 0.234 | 0.111 | 0.124 | 0.119 | 0.118 | 0.102 - 0.134 | 0.215 | 0.185 | 0.192 | 0.197 | 0.158 - 0.236 | no |
| cSP | CLIPC | CLIPC7 | AGAP003689 | MelASA | positive | 0.217 | 0.208 | 0.195 | 0.207 | 0.179 - 0.234 | 0.111 | 0.124 | 0.119 | 0.118 | 0.102 - 0.134 | 0.206 | 0.189 | 0.203 | 0.199 | 0.177 - 0.222 | no |
| cSP | CLIPC | CLIPC9 | AGAP004719 | MelASA | positive | 0.217 | 0.208 | 0.195 | 0.207 | 0.179 - 0.234 | 0.111 | 0.124 | 0.119 | 0.118 | 0.102 - 0.134 | 0.152 | 0.159 | 0.140 | 0.150 | 0.127 - 0.174 | yes |
| cSP | CLIPC | CLIPC10 | AGAP000572 | MelASA | positive | 0.217 | 0.208 | 0.195 | 0.207 | 0.179 - 0.234 | 0.111 | 0.124 | 0.119 | 0.118 | 0.102 - 0.134 | 0.202 | 0.192 | 0.184 | 0.193 | 0.170 - 0.215 | no |
| cSP | CLIPC | CLIPC12 | AGAP012034 | MelASA | positive | 0.217 | 0.208 | 0.195 | 0.207 | 0.179 - 0.234 | 0.111 | 0.124 | 0.119 | 0.118 | 0.102 - 0.134 | 0.139 | 0.123 | 0.141 | 0.134 | 0.110 - 0.159 | yes |
| cSP | CLIPC | CLIPC14 | AGAP028167 | MelASA | positive | 0.217 | 0.208 | 0.195 | 0.207 | 0.179 - 0.234 | 0.111 | 0.124 | 0.119 | 0.118 | 0.102 - 0.134 | 0.123 | 0.134 | 0.148 | 0.135 | 0.104 - 0.166 | yes |
| cSP | CLIPD | CLIPD1 | AGAP002422 | MelASA | positive | 0.191 | 0.182 | 0.191 | 0.188 | 0.175 - 0.201 | 0.124 | 0.126 | 0.114 | 0.121 | 0.105 - 0.137 | 0.249 | 0.189 | 0.219 | 0.219 | 0.145 - 0.294 | no |
| cSP | CLIPD | CLIPD2 | AGAP008183 | MelASA | positive | 0.191 | 0.182 | 0.191 | 0.188 | 0.175 - 0.201 | 0.124 | 0.126 | 0.114 | 0.121 | 0.105 - 0.137 | 0.178 | 0.205 | 0.188 | 0.190 | 0.156 - 0.224 | no |
| cSP | CLIPD | CLIPD3 | AGAP001433 | MelASA | positive | 0.205 | 0.217 | 0.223 | 0.215 | 0.192 - 0.238 | 0.122 | 0.119 | 0.135 | 0.125 | 0.104 - 0.147 | 0.192 | 0.189 | 0.201 | 0.194 | 0.179 - 0.210 | no |
| cSP | CLIPD | CLIPD4 | AGAP002811 | MelASA | positive | 0.191 | 0.182 | 0.191 | 0.188 | 0.175 - 0.201 | 0.124 | 0.126 | 0.114 | 0.121 | 0.105 - 0.137 | 0.185 | 0.178 | 0.201 | 0.188 | 0.159 - 0.217 | no |
| cSP | CLIPD | CLIPD6 | AGAP002813 | MelASA | positive | 0.191 | 0.182 | 0.191 | 0.188 | 0.175 - 0.201 | 0.124 | 0.126 | 0.114 | 0.121 | 0.105 - 0.137 | 0.216 | 0.213 | 0.199 | 0.209 | 0.187 - 0.232 | no |
| cSP | CLIPD | CLIPD7 | AGAP008998 | MelASA | positive | 0.191 | 0.182 | 0.191 | 0.188 | 0.175 - 0.201 | 0.124 | 0.126 | 0.114 | 0.121 | 0.105 - 0.137 | 0.209 | 0.207 | 0.195 | 0.204 | 0.185 - 0.223 | no |
| cSP | CLIPD | CLIPD8 | AGAP002784 | MelASA | positive | 0.191 | 0.182 | 0.191 | 0.188 | 0.175 - 0.201 | 0.124 | 0.126 | 0.114 | 0.121 | 0.105 - 0.137 | 0.193 | 0.188 | 0.204 | 0.195 | 0.175 - 0.215 | no |
| cSP | CLIPD | CLIPD9 | AGAP008997 | MelASA | positive | 0.191 | 0.182 | 0.191 | 0.188 | 0.175 - 0.201 | 0.124 | 0.126 | 0.114 | 0.121 | 0.105 - 0.137 | 0.201 | 0.209 | 0.173 | 0.194 | 0.147 - 0.241 | no |
| cSP | CLIPD | CLIPD11 | AGAP029106 | MelASA | positive | 0.191 | 0.182 | 0.191 | 0.188 | 0.175 - 0.201 | 0.117 | 0.122 | 0.126 | 0.122 | 0.105 - 0.137 | 0.203 | 0.211 | 0.205 | 0.206 | 0.196 - 0.217 | no |
| cSP | CLIPD | CLIPD12 | AGAP008995 | MelASA | positive | 0.194 | 0.204 | 0.197 | 0.198 | 0.186 - 0.211 | 0.117 | 0.122 | 0.126 | 0.122 | 0.111 - 0.133 | 0.214 | 0.195 | 0.199 | 0.203 | 0.178 - 0.228 | no |
| cSP | CLIPD | CLIPD13 | AGAP009000 | MelASA | positive | 0.194 | 0.204 | 0.197 | 0.198 | 0.186 - 0.211 | 0.117 | 0.122 | 0.126 | 0.122 | 0.111 - 0.133 | 0.205 | 0.216 | 0.203 | 0.208 | 0.191 - 0.225 | no |
| cSP | CLIPD | CLIPD14 | AGAP009006 | MelASA | positive | 0.194 | 0.204 | 0.197 | 0.198 | 0.186 - 0.211 | 0.117 | 0.122 | 0.126 | 0.122 | 0.111 - 0.133 | 0.207 | 0.194 | 0.209 | 0.203 | 0.183 - 0.224 | no |
| cSP | CLIPD | CLIPD20 | AGAP013089 | MelASA | positive | 0.194 | 0.204 | 0.197 | 0.198 | 0.186 - 0.211 | 0.117 | 0.122 | 0.126 | 0.122 | 0.111 - 0.133 | 0.206 | 0.210 | 0.199 | 0.205 | 0.191 - 0.219 | no |
| cSP | CLIPD | CLIPD22 | AGAP008996 | MelASA | positive | 0.194 | 0.204 | 0.197 | 0.198 | 0.186 - 0.211 | 0.117 | 0.122 | 0.126 | 0.122 | 0.111 - 0.133 | 0.193 | 0.203 | 0.215 | 0.204 | 0.176 - 0.231 | no |
| cSPH | CLIPE | CLIPE1 | AGAP008091 | MelASA | positive | 0.191 | 0.192 | 0.181 | 0.188 | 0.173 - 0.203 | 0.134 | 0.126 | 0.104 | 0.121 | 0.083 - 0.160 | 0.198 | 0.184 | 0.196 | 0.193 | 0.174 - 0.212 | no |
| cSPH | CLIPE | CLIPE4 | AGAP010530 | MelASA | positive | 0.191 | 0.192 | 0.181 | 0.188 | 0.173 - 0.203 | 0.134 | 0.126 | 0.104 | 0.121 | 0.083 - 0.160 | 0.183 | 0.180 | 0.173 | 0.179 | 0.166 - 0.191 | no |
| cSP | CLIPE | CLIPE5 | AGAP028728 | MelASA | positive | 0.191 | 0.192 | 0.181 | 0.188 | 0.173 - 0.203 | 0.134 | 0.126 | 0.104 | 0.121 | 0.083 - 0.160 | 0.199 | 0.179 | 0.189 | 0.189 | 0.164 - 0.214 | no |
| cSPH | CLIPE | CLIPE6 | AGAP011785 | MelASA | positive | 0.191 | 0.192 | 0.181 | 0.188 | 0.173 - 0.203 | 0.134 | 0.126 | 0.104 | 0.121 | 0.083 - 0.160 | 0.178 | 0.209 | 0.198 | 0.195 | 0.156 - 0.234 | no |
| cSPH | CLIPE | CLIPE7 | AGAP011786 | MelASA | positive | 0.191 | 0.192 | 0.181 | 0.188 | 0.173 - 0.203 | 0.134 | 0.126 | 0.104 | 0.121 | 0.083 - 0.160 | 0.185 | 0.198 | 0.201 | 0.195 | 0.174 - 0.216 | no |
| cSP | CLIPE | CLIPE10 | AGAP010545 | MelASA | positive | 0.191 | 0.192 | 0.181 | 0.188 | 0.173 - 0.203 | 0.134 | 0.126 | 0.104 | 0.121 | 0.083 - 0.160 | 0.216 | 0.203 | 0.189 | 0.203 | 0.169 - 0.236 | no |
| cSPH | CLIPE | CLIPE12 | AGAP003691 | MelASA | positive | 0.191 | 0.192 | 0.181 | 0.188 | 0.173 - 0.203 | 0.134 | 0.126 | 0.104 | 0.121 | 0.083 - 0.160 | 0.209 | 0.197 | 0.205 | 0.204 | 0.189 - 0.219 | no |
| cSP | CLIPE | CLIPE13 | AGAP003748 | MelASA | positive | 0.191 | 0.192 | 0.181 | 0.188 | 0.173 - 0.203 | 0.134 | 0.126 | 0.104 | 0.121 | 0.083 - 0.160 | 0.193 | 0.178 | 0.184 | 0.185 | 0.166 - 0.204 | no |
| cSP | CLIPE | CLIPE15 | AGAP009249 | MelASA | positive | 0.191 | 0.192 | 0.181 | 0.188 | 0.173 - 0.203 | 0.134 | 0.126 | 0.104 | 0.121 | 0.083 - 0.160 | 0.201 | 0.199 | 0.193 | 0.198 | 0.187 - 0.208 | no |
| cSPH | CLIPE | CLIPE17 | AGAP009252 | MelASA | positive | 0.191 | 0.192 | 0.181 | 0.188 | 0.173 - 0.203 | 0.134 | 0.126 | 0.104 | 0.121 | 0.083 - 0.160 | 0.203 | 0.171 | 0.215 | 0.196 | 0.140 - 0.253 | no |
| cSP | CLIPE | CLIPE18 | AGAP009273 | MelASA | positive | 0.202 | 0.207 | 0.189 | 0.199 | 0.176 - 0.222 | 0.124 | 0.106 | 0.131 | 0.120 | 0.088 - 0.151 | 0.178 | 0.195 | 0.201 | 0.191 | 0.162 - 0.221 | no |
| cSP | CLIPE | CLIPE19 | AGAP011040 | MelASA | positive | 0.202 | 0.207 | 0.189 | 0.199 | 0.176 - 0.222 | 0.124 | 0.106 | 0.131 | 0.120 | 0.088 - 0.151 | 0.184 | 0.175 | 0.223 | 0.194 | 0.131 - 0.257 | no |
| cSP | CLIPE | CLIPE21 | AGAP011719 | MelASA | positive | 0.202 | 0.207 | 0.189 | 0.199 | 0.176 - 0.222 | 0.124 | 0.106 | 0.131 | 0.120 | 0.088 - 0.151 | 0.173 | 0.192 | 0.189 | 0.185 | 0.159 - 0.210 | no |
| cSP | CLIPE | CLIPE22 | AGAP012020 | MelASA | positive | 0.202 | 0.207 | 0.189 | 0.199 | 0.176 - 0.222 | 0.124 | 0.106 | 0.131 | 0.120 | 0.088 - 0.151 | 0.195 | 0.178 | 0.194 | 0.189 | 0.165 - 0.213 | no |
| cSP | CLIPE | CLIPE25 | AGAP012502 | MelASA | positive | 0.202 | 0.207 | 0.189 | 0.199 | 0.176 - 0.222 | 0.124 | 0.106 | 0.131 | 0.120 | 0.088 - 0.151 | 0.201 | 0.198 | 0.207 | 0.202 | 0.191 - 0.213 | no |
| cSP | CLIPE | CLIPE30 | AGAP010628 | MelASA | positive | 0.202 | 0.207 | 0.189 | 0.199 | 0.176 - 0.222 | 0.124 | 0.106 | 0.131 | 0.120 | 0.088 - 0.151 | 0.192 | 0.225 | 0.192 | 0.203 | 0.156 - 0.250 | no |

**Table S4: MelASA RNAi screen: Total melanotic spot area of genes tested as negative regulators.** DsRNA injected mosquitoes were challenged with 50.6 nL of resuspended lyophilized *Micrococcus luteus* at $OD_{600} = 0.1$. Batches of 7-10 candidate genes were tested in independent groups using their own positive (ds*SRPN2)* and negative controls. Three independent biological replicates were performed using different mosquito generations.

| CLIP family | sub family | Gene name | AGAP | Assay type | Tested as | ds*GFP* control (individual replicates) | | | | | Positive control-ds*SRPN2* (individual replicates) | | | | | ds*GOI* (individual replicates) | | | | | Phenotype |
|---|---|---|---|---|---|---|---|---|---|---|---|---|---|---|---|---|---|---|---|---|---|
| | | | | | | 1 | 2 | 3 | Mean | 95% CI of mean | 1 | 2 | 3 | Mean | 95% CI of mean | 1 | 2 | 3 | Mean | 95% CI of mean | |
| cSPH | CLIPA | CLIPA1 | AGAP011791 | MelASA | negative | 0.033 | 0.035 | 0.045 | 0.038 | 0.176 - 0.222 | 0.116 | 0.121 | 0.129 | 0.122 | 0.106 - 0.138 | 0.031 | 0.039 | 0.048 | 0.039 | 0.018 - 0.060 | no |
| cSPH | CLIPA | CLIPA2 | AGAP011790 | MelASA | negative | 0.037 | 0.043 | 0.039 | 0.040 | 0.022 - 0.054 | 0.138 | 0.127 | 0.149 | 0.138 | 0.106 - 0.138 | 0.099 | 0.105 | 0.117 | 0.107 | 0.084 - 0.130 | yes |
| cSPH | CLIPA | CLIPA3 | AGAP012591 | MelASA | negative | 0.037 | 0.043 | 0.039 | 0.040 | 0.022 - 0.054 | 0.138 | 0.127 | 0.149 | 0.138 | 0.106 - 0.138 | 0.043 | 0.038 | 0.038 | 0.040 | 0.033 - 0.047 | no |
| cSPH | CLIPA | CLIPA4 | AGAP011780 | MelASA | negative | 0.037 | 0.043 | 0.039 | 0.040 | 0.022 - 0.054 | 0.138 | 0.127 | 0.149 | 0.138 | 0.106 - 0.138 | 0.039 | 0.043 | 0.039 | 0.040 | 0.035 - 0.046 | no |
| cSPH | CLIPA | CLIPA5 | AGAP011787 | MelASA | negative | 0.037 | 0.043 | 0.039 | 0.040 | 0.022 - 0.054 | 0.138 | 0.127 | 0.149 | 0.138 | 0.106 - 0.138 | 0.079 | 0.092 | 0.106 | 0.092 | 0.059 - 0.126 | yes |
| cSPH | CLIPA | CLIPA6 | AGAP011789 | MelASA | negative | 0.037 | 0.043 | 0.039 | 0.040 | 0.022 - 0.054 | 0.138 | 0.127 | 0.149 | 0.138 | 0.106 - 0.138 | 0.039 | 0.036 | 0.028 | 0.034 | 0.020 - 0.048 | no |
| cSPH | CLIPA | CLIPA7 | AGAP011792 | MelASA | negative | 0.037 | 0.043 | 0.039 | 0.040 | 0.022 - 0.054 | 0.138 | 0.127 | 0.149 | 0.138 | 0.106 - 0.138 | 0.064 | 0.078 | 0.091 | 0.078 | 0.044 - 0.111 | yes |
| cSPH | CLIPA | CLIPA8 | AGAP010731 | MelASA | negative | 0.037 | 0.043 | 0.039 | 0.040 | 0.022 - 0.054 | 0.138 | 0.127 | 0.149 | 0.138 | 0.106 - 0.138 | 0.018 | 0.014 | 0.011 | 0.014 | 0.006 - 0.023 | no |
| cSPH | CLIPA | CLIPA9 | AGAP010968 | MelASA | negative | 0.037 | 0.043 | 0.039 | 0.040 | 0.022 - 0.054 | 0.138 | 0.127 | 0.149 | 0.138 | 0.106 - 0.138 | 0.034 | 0.039 | 0.043 | 0.039 | 0.027 - 0.050 | no |
| cSPH | CLIPA | CLIPA10 | AGAP006954 | MelASA | negative | 0.037 | 0.043 | 0.039 | 0.040 | 0.022 - 0.054 | 0.138 | 0.127 | 0.149 | 0.138 | 0.106 - 0.138 | 0.041 | 0.035 | 0.037 | 0.038 | 0.030 - 0.045 | no |
| cSPH | CLIPA | CLIPA12 | AGAP011781 | MelASA | negative | 0.037 | 0.043 | 0.039 | 0.040 | 0.022 - 0.054 | 0.138 | 0.127 | 0.149 | 0.138 | 0.106 - 0.138 | 0.032 | 0.047 | 0.05 | 0.043 | 0.019 - 0.067 | no |
| cSPH | CLIPA | CLIPA13 | AGAP011783 | MelASA | negative | 0.032 | 0.047 | 0.041 | 0.040 | 0.021 - 0.059 | 0.127 | 0.114 | 0.131 | 0.124 | 0.102 - 0.146 | 0.044 | 0.068 | 0.061 | 0.058 | 0.027 - 0.088 | yes |
| cSPH | CLIPA | CLIPA14 | AGAP011788 | MelASA | negative | 0.032 | 0.047 | 0.041 | 0.040 | 0.021 - 0.059 | 0.127 | 0.114 | 0.131 | 0.124 | 0.102 - 0.146 | 0.102 | 0.099 | 0.097 | 0.099 | 0.093 - 0.106 | yes |
| cSPH | CLIPA | CLIPA15 | AGAP002815 | MelASA | negative | 0.032 | 0.047 | 0.041 | 0.040 | 0.021 - 0.059 | 0.127 | 0.114 | 0.131 | 0.124 | 0.102 - 0.146 | 0.029 | 0.046 | 0.041 | 0.039 | 0.017 - 0.060 | no |
| cSPH | CLIPA | CLIPA19 | AGAP003245 | MelASA | negative | 0.032 | 0.047 | 0.041 | 0.040 | 0.021 - 0.059 | 0.127 | 0.114 | 0.131 | 0.124 | 0.102 - 0.146 | 0.032 | 0.074 | 0.038 | 0.048 | -0.008 - 0.104 | no |
| cSPH | CLIPA | CLIPA26 | AGAP001964 | MelASA | negative | 0.032 | 0.047 | 0.041 | 0.040 | 0.021 - 0.059 | 0.127 | 0.114 | 0.131 | 0.124 | 0.102 - 0.146 | 0.031 | 0.056 | 0.039 | 0.042 | 0.010 - 0.074 | no |
| cSPH | CLIPA | CLIPA27 | AGAP000290 | MelASA | negative | 0.032 | 0.047 | 0.041 | 0.040 | 0.021 - 0.059 | 0.127 | 0.114 | 0.131 | 0.124 | 0.102 - 0.146 | 0.028 | 0.042 | 0.039 | 0.036 | 0.018 - 0.055 | no |
| cSPH | CLIPA | CLIPA28 | AGAP010730 | MelASA | negative | 0.032 | 0.047 | 0.041 | 0.040 | 0.021 - 0.059 | 0.127 | 0.114 | 0.131 | 0.124 | 0.102 - 0.146 | 0.031 | 0.043 | 0.034 | 0.036 | 0.020 - 0.052 | no |
| cSPH | CLIPA | SPCLIP1 | AGAP028725 | MelASA | negative | 0.032 | 0.047 | 0.041 | 0.040 | 0.021 - 0.059 | 0.127 | 0.114 | 0.131 | 0.124 | 0.102 - 0.146 | 0.017 | 0.025 | 0.019 | 0.020 | 0.010 - 0.031 | no |
| cSPH | CLIPB | CLIPB7 | AGAP002270 | MelASA | negative | 0.043 | 0.055 | 0.038 | 0.045 | 0.024 - 0.067 | 0.147 | 0.143 | 0.131 | 0.140 | 0.120 - 0.161 | 0.038 | 0.045 | 0.033 | 0.039 | 0.024 - 0.054 | no |
| cSPH | CLIPB | CLIPB12 | AGAP009217 | MelASA | negative | 0.043 | 0.055 | 0.038 | 0.045 | 0.024 - 0.067 | 0.147 | 0.143 | 0.131 | 0.140 | 0.120 - 0.161 | 0.067 | 0.078 | 0.086 | 0.077 | 0.053 - 0.101 | yes |
| cSPH | CLIPB | CLIPB16 | AGAP009263 | MelASA | negative | 0.043 | 0.055 | 0.038 | 0.045 | 0.024 - 0.067 | 0.147 | 0.143 | 0.131 | 0.140 | 0.120 - 0.161 | 0.036 | 0.044 | 0.043 | 0.041 | 0.030 - 0.052 | no |
| cSPH | CLIPB | CLIPB36 | AGAP013184 | MelASA | negative | 0.043 | 0.055 | 0.038 | 0.045 | 0.024 - 0.067 | 0.147 | 0.143 | 0.131 | 0.140 | 0.120 - 0.161 | 0.045 | 0.048 | 0.039 | 0.044 | 0.033 - 0.055 | no |
| cSPH | CLIPC | CLIPC7 | AGAP003689 | MelASA | negative | 0.033 | 0.035 | 0.045 | 0.038 | 0.022 - 0.054 | 0.116 | 0.121 | 0.129 | 0.122 | 0.106 - 0.138 | 0.033 | 0.04 | 0.041 | 0.038 | 0.027 - 0.049 | no |
| cSPH | CLIPD | CLIPD14 | AGAP009006 | MelASA | negative | 0.033 | 0.035 | 0.045 | 0.038 | 0.022 - 0.054 | 0.116 | 0.121 | 0.129 | 0.122 | 0.106 - 0.138 | 0.039 | 0.041 | 0.044 | 0.041 | 0.035 - 0.048 | no |
| cSPH | CLIPE | CLIPE1 | AGAP008091 | MelASA | negative | 0.033 | 0.035 | 0.045 | 0.038 | 0.022 - 0.054 | 0.116 | 0.121 | 0.129 | 0.122 | 0.106 - 0.138 | 0.031 | 0.038 | 0.042 | 0.037 | 0.023 - 0.051 | no |
| cSPH | CLIPE | CLIPE4 | AGAP010530 | MelASA | negative | 0.033 | 0.035 | 0.045 | 0.038 | 0.022 - 0.054 | 0.116 | 0.121 | 0.129 | 0.122 | 0.106 - 0.138 | 0.034 | 0.045 | 0.051 | 0.043 | 0.022 - 0.065 | no |
| cSPH | CLIPE | CLIPE6 | AGAP011785 | MelASA | negative | 0.033 | 0.035 | 0.045 | 0.038 | 0.022 - 0.054 | 0.116 | 0.121 | 0.129 | 0.122 | 0.106 - 0.138 | 0.042 | 0.037 | 0.048 | 0.042 | 0.029 - 0.056 | no |
| cSPH | CLIPE | CLIPE7 | AGAP011786 | MelASA | negative | 0.033 | 0.035 | 0.045 | 0.038 | 0.022 - 0.054 | 0.116 | 0.121 | 0.129 | 0.122 | 0.106 - 0.138 | 0.035 | 0.044 | 0.053 | 0.044 | 0.022 - 0.066 | no |
| cSPH | CLIPE | CLIPE12 | AGAP003691 | MelASA | negative | 0.033 | 0.035 | 0.045 | 0.038 | 0.022 - 0.054 | 0.116 | 0.121 | 0.129 | 0.122 | 0.106 - 0.138 | 0.038 | 0.041 | 0.044 | 0.041 | 0.034 - 0.048 | no |
| cSPH | CLIPE | CLIPE17 | AGAP009252 | MelASA | negative | 0.033 | 0.035 | 0.045 | 0.038 | 0.022 - 0.054 | 0.116 | 0.121 | 0.129 | 0.122 | 0.106 - 0.138 | 0.04 | 0.047 | 0.052 | 0.046 | 0.031 - 0.061 | no |

**Table S5: ZOI screen: Zone of inhibition of genes tested as positive regulators.** DsRNA injected mosquitoes were challenged with 50.6 nL of resuspended lyophilized *Micrococcus luteus* at OD600 = 5. Batches of 7-10 candidate genes were tested in independent groups using their own positive (ds*REL1*) and negative controls. Three independent biological replicates were performed using different mosquito generations.

| CLIP family | sub family | Gene name | AGAP | Assay type | Tested as | ds*GFP* controls (individual replicates) | | | | | Positive controls-ds*REL1* (individual replicates) | | | | | ds*GOI* (individual replicates) | | | | | Phenotype |
|---|---|---|---|---|---|---|---|---|---|---|---|---|---|---|---|---|---|---|---|---|---|
| | | | | | | 1 | 2 | 3 | Mean | 95% CI of mean | 1 | 2 | 3 | Mean | 95% CI of mean | 1 | 2 | 3 | Mean | 95% CI of mean | |
| | | TEP1 | AGAP010815 | ZOI | positive | 35.031 | 33.602 | 31.849 | 33.512 | 29.530 - 37.450 | 17.184 | 18.267 | 15.213 | 16.888 | 13.040 - 20.730 | 34.816 | 32.157 | 31.884 | 32.595 | 28.930 - 36.980 | no |
| | | MODSP1 | AGAP001798 | ZOI | positive | 33.799 | 35.920 | 30.816 | 33.512 | 27.140 - 39.880 | 19.012 | 16.212 | 17.164 | 17.463 | 13.930 - 21.000 | 32.649 | 34.284 | 30.853 | 32.595 | 28.330 - 36.860 | no |
| cSPH | CLIPA | CLIPA1 | AGAP011791 | ZOI | positive | 31.951 | 34.689 | 35.026 | 33.889 | 29.700 - 38.080 | 15.192 | 12.623 | 16.820 | 14.878 | 9.622 - 20.130 | 36.67 | 29.96 | 31.3 | 32.642 | 23.820 - 41.460 | no |
| cSPH | CLIPA | CLIPA2 | AGAP011790 | ZOI | positive | 31.951 | 34.689 | 35.026 | 33.889 | 30.700 - 39.080 | 15.192 | 12.623 | 16.820 | 14.878 | 9.622 - 20.130 | 28.38 | 28.75 | 34.62 | 30.582 | 21.890 - 39.270 | no |
| cSPH | CLIPA | CLIPA3 | AGAP012591 | ZOI | positive | 31.951 | 34.689 | 35.026 | 33.889 | 31.700 - 40.080 | 15.192 | 12.623 | 16.820 | 14.878 | 9.622 - 20.130 | 14.59 | 19.63 | 15.16 | 16.459 | 9.610 - 23.310 | yes |
| cSPH | CLIPA | CLIPA4 | AGAP011780 | ZOI | positive | 31.951 | 34.689 | 35.026 | 33.889 | 32.700 - 41.080 | 15.192 | 12.623 | 16.820 | 14.878 | 9.622 - 20.130 | 35.53 | 35.21 | 34.06 | 34.933 | 33.010 - 36.850 | no |
| cSPH | CLIPA | CLIPA5 | AGAP011787 | ZOI | positive | 31.951 | 34.689 | 35.026 | 33.889 | 33.700 - 42.080 | 15.192 | 12.623 | 16.820 | 14.878 | 9.622 - 20.130 | 34.29 | 30.99 | 30.67 | 31.984 | 27.000 - 36.970 | no |
| cSPH | CLIPA | CLIPA6 | AGAP011789 | ZOI | positive | 31.951 | 34.689 | 35.026 | 33.889 | 34.700 - 43.080 | 15.192 | 12.623 | 16.820 | 14.878 | 9.622 - 20.130 | 28.44 | 30.33 | 33.15 | 30.636 | 24.750 - 36.520 | no |
| cSPH | CLIPA | CLIPA7 | AGAP011792 | ZOI | positive | 31.951 | 34.689 | 35.026 | 33.889 | 35.700 - 44.080 | 15.192 | 12.623 | 16.820 | 14.878 | 9.622 - 20.130 | 34.28 | 31.22 | 34.9 | 33.466 | 28.580 - 38.350 | no |
| cSPH | CLIPA | CLIPA8 | AGAP010731 | ZOI | positive | 31.951 | 34.689 | 35.026 | 33.889 | 36.700 - 45.080 | 15.192 | 12.623 | 16.820 | 14.878 | 9.622 - 20.130 | 34.63 | 28.92 | 29.21 | 30.919 | 22.930 - 38.910 | no |
| cSPH | CLIPA | CLIPA9 | AGAP010968 | ZOI | positive | 31.951 | 34.689 | 35.026 | 33.889 | 37.700 - 46.080 | 15.192 | 12.623 | 16.820 | 14.878 | 9.622 - 20.130 | 33.94 | 34.81 | 34.88 | 34.542 | 33.250 - 35.840 | no |
| cSPH | CLIPA | CLIPA10 | AGAP006954 | ZOI | positive | 31.951 | 34.689 | 35.026 | 33.889 | 38.700 - 47.080 | 15.192 | 12.623 | 16.820 | 14.878 | 9.622 - 20.130 | 32.57 | 31.38 | 29.47 | 31.140 | 27.250 - 35.030 | no |
| cSPH | CLIPA | CLIPA12 | AGAP011781 | ZOI | positive | 31.951 | 34.689 | 35.026 | 33.889 | 39.700 - 48.080 | 15.192 | 12.623 | 16.820 | 14.878 | 9.622 - 20.130 | 28 | 34.76 | 32.1 | 31.622 | 23.160 - 40.080 | no |
| cSPH | CLIPA | CLIPA13 | AGAP011783 | ZOI | positive | 32.048 | 33.297 | 29.284 | 31.543 | 26.440 - 36.640 | 14.234 | 14.134 | 18.197 | 15.522 | 9.765 - 21.280 | 35.29 | 29.29 | 32.22 | 32.265 | 24.800 - 39.730 | no |
| cSPH | CLIPA | CLIPA14 | AGAP011788 | ZOI | positive | 32.048 | 33.297 | 29.284 | 31.543 | 26.440 - 36.640 | 14.234 | 14.134 | 18.197 | 15.522 | 9.765 - 21.280 | 31.29 | 29.73 | 33.26 | 31.427 | 27.040 - 35.820 | no |
| cSPH | CLIPA | CLIPA15 | AGAP002815 | ZOI | positive | 32.048 | 33.297 | 29.284 | 31.543 | 26.440 - 36.640 | 14.234 | 14.134 | 18.197 | 15.522 | 9.765 - 21.280 | 29.39 | 31.27 | 34.2 | 31.618 | 25.600 - 37.630 | no |
| cSPH | CLIPA | CLIPA19 | AGAP003245 | ZOI | positive | 32.048 | 33.297 | 29.284 | 31.543 | 26.440 - 36.640 | 14.234 | 14.134 | 18.197 | 15.522 | 9.765 - 21.280 | 31.37 | 32.27 | 31.25 | 31.631 | 30.240 - 33.020 | no |
| cSPH | CLIPA | CLIPA26 | AGAP001964 | ZOI | positive | 32.048 | 33.297 | 29.284 | 31.543 | 26.440 - 36.640 | 14.234 | 14.134 | 18.197 | 15.522 | 9.765 - 21.280 | 30.19 | 30.39 | 31.71 | 30.765 | 28.720 - 32.810 | no |
| cSPH | CLIPA | CLIPA27 | AGAP000290 | ZOI | positive | 32.048 | 33.297 | 29.284 | 31.543 | 26.440 - 36.640 | 14.234 | 14.134 | 18.197 | 15.522 | 9.765 - 21.280 | 32.19 | 31.27 | 32.25 | 31.905 | 30.540 - 33.270 | no |
| cSPH | CLIPA | CLIPA28 | AGAP010730 | ZOI | positive | 32.048 | 33.297 | 29.284 | 31.543 | 26.440 - 36.640 | 14.234 | 14.134 | 18.197 | 15.522 | 9.765 - 21.280 | 28.93 | 32.39 | 29.39 | 30.240 | 25.570 - 34.910 | no |
| cSPH | CLIPA | SPCLIP1 | AGAP028725 | ZOI | positive | 32.048 | 33.297 | 29.284 | 31.543 | 26.440 - 36.640 | 14.234 | 14.134 | 18.197 | 15.522 | 9.765 - 21.280 | 29.91 | 34.76 | 32.22 | 32.298 | 26.270 - 38.330 | no |
| cSP | CLIPB | CLIPB1 | AGAP003251 | ZOI | positive | 36.106 | 36.816 | 32.499 | 35.140 | 29.390 - 40.890 | 18.140 | 18.661 | 17.835 | 18.212 | 17.170 - 19.250 | 30.69 | 33.57 | 35.19 | 33.148 | 27.480 - 38.820 | no |
| cSP | CLIPB | CLIPB2 | AGAP003246 | ZOI | positive | 36.106 | 36.816 | 32.499 | 35.140 | 29.390 - 40.890 | 18.140 | 18.661 | 17.835 | 18.212 | 17.170 - 19.250 | 31.53 | 38.1 | 34.53 | 34.722 | 26.560 - 42.890 | no |
| cSP | CLIPB | CLIPB3b | AGAP003249 | ZOI | positive | 36.106 | 36.816 | 32.499 | 35.140 | 29.390 - 40.890 | 18.140 | 18.661 | 17.835 | 18.212 | 17.170 - 19.250 | 37.67 | 27.95 | 35.1 | 33.573 | 21.050 - 46.090 | no |
| cSP | CLIPB | CLIPB5 | AGAP004148 | ZOI | positive | 36.106 | 36.816 | 32.499 | 35.140 | 29.390 - 40.890 | 18.140 | 18.661 | 17.835 | 18.212 | 17.170 - 19.250 | 32.19 | 38.25 | 33.15 | 34.531 | 26.440 - 42.620 | no |
| cSP | CLIPB | CLIPB6 | AGAP003252 | ZOI | positive | 36.106 | 36.816 | 32.499 | 35.140 | 29.390 - 40.890 | 18.140 | 18.661 | 17.835 | 18.212 | 17.170 - 19.250 | 32.43 | 30.6 | 32.18 | 31.735 | 29.270 - 34.200 | no |
| cSPH | CLIPB | CLIPB7 | AGAP002270 | ZOI | positive | 36.106 | 36.816 | 32.499 | 35.140 | 29.390 - 40.890 | 18.140 | 18.661 | 17.835 | 18.212 | 17.170 - 19.250 | 32.64 | 30.29 | 35.64 | 32.859 | 26.200 - 39.520 | no |
| cSP | CLIPB | CLIPB8 | AGAP003057 | ZOI | positive | 33.799 | 35.92 | 30.816 | 33.512 | 27.140 - 39.880 | 19.012 | 16.212 | 17.164 | 17.463 | 13.930 - 21.000 | 34.95 | 32.85 | 30.35 | 32.716 | 27.000 - 38.430 | no |
| cSP | CLIPB | CLIPB9 | AGAP029769 | ZOI | positive | 33.799 | 35.92 | 30.816 | 33.512 | 28.140 - 40.880 | 19.012 | 16.212 | 17.164 | 17.463 | 13.930 - 21.000 | 31.8 | 30.38 | 32.46 | 31.550 | 28.910 - 34.190 | no |
| cSP | CLIPB | CLIPB10 | AGAP029770 | ZOI | positive | 33.799 | 35.92 | 30.816 | 33.512 | 29.140 - 41.880 | 19.012 | 16.212 | 17.164 | 17.463 | 13.930 - 21.000 | 14.16 | 16.02 | 15.38 | 15.183 | 12.840 - 17.530 | yes |
| cSP | CLIPB | CLIPB11 | AGAP009214 | ZOI | positive | 36.106 | 36.816 | 32.499 | 35.140 | 29.390 - 40.890 | 18.140 | 18.661 | 17.835 | 18.212 | 17.170 - 19.250 | 37.85 | 38.24 | 31.75 | 35.942 | 26.900 - 44.980 | no |
| cSPH | CLIPB | CLIPB12 | AGAP009217 | ZOI | positive | 34.073 | 33.706 | 33.895 | 33.891 | 33.440 - 34.350 | 14.611 | 18.889 | 17.587 | 17.029 | 11.580 - 22.480 | 32.94 | 34.66 | 35.83 | 34.477 | 30.860 - 38.090 | no |
| cSP | CLIPB | CLIPB13 | AGAP004855 | ZOI | positive | 31.901 | 32.36 | 35.805 | 33.355 | 28.050 - 38.660 | 17.426 | 13.105 | 15.204 | 15.245 | 9.877 - 20.610 | 28.89 | 30.58 | 31.43 | 30.298 | 27.090 - 33.510 | no |
| cSP | CLIPB | CLIPB14 | AGAP010833 | ZOI | positive | 31.901 | 32.36 | 35.805 | 33.355 | 28.050 - 38.660 | 17.426 | 13.105 | 15.204 | 15.245 | 9.877 - 20.610 | 10.93 | 14.37 | 13.53 | 12.946 | 8.491 - 17.400 | yes |
| cSP | CLIPB | CLIPB15 | AGAP009844 | ZOI | positive | 31.901 | 32.36 | 35.805 | 33.355 | 28.050 - 38.660 | 17.426 | 13.105 | 15.204 | 15.245 | 9.877 - 20.610 | 32.5 | 37.72 | 36.99 | 35.735 | 28.710 - 42.760 | no |
| cSPH | CLIPB | CLIPB16 | AGAP009263 | ZOI | positive | 31.901 | 32.36 | 35.805 | 33.355 | 28.050 - 38.660 | 17.426 | 13.105 | 15.204 | 15.245 | 9.877 - 20.610 | 30.45 | 36.5 | 30.94 | 32.631 | 24.280 - 40.980 | no |
| cSP | CLIPB | CLIPB17 | AGAP001648 | ZOI | positive | 34.84 | 34.732 | 33.571 | 34.381 | 32.630 - 36.130 | 16.473 | 19.127 | 15.678 | 17.093 | 12.610 - 21.580 | 30.39 | 34.78 | 31.2 | 32.122 | 26.330 - 37.920 | no |
| cSP | CLIPB | CLIPB18 | AGAP009215 | ZOI | positive | 34.84 | 34.732 | 33.571 | 34.381 | 32.630 - 36.130 | 16.473 | 19.127 | 15.678 | 17.093 | 12.610 - 21.580 | 31.37 | 34.15 | 30.55 | 32.024 | 27.340 - 36.710 | no |
| cSP | CLIPB | CLIPB19 | AGAP003247 | ZOI | positive | 31.901 | 32.36 | 35.805 | 33.355 | 28.050 - 38.660 | 17.426 | 13.105 | 15.204 | 15.245 | 9.877 - 20.610 | 36.41 | 31.25 | 32.78 | 33.480 | 26.890 - 40.070 | no |
| cSP | CLIPB | CLIPB20 | AGAP012037 | ZOI | positive | 36.106 | 36.816 | 32.499 | 35.140 | 29.390 - 40.890 | 18.140 | 18.661 | 17.835 | 18.212 | 17.170 - 19.250 | 31.41 | 37.45 | 32.76 | 33.871 | 26.000 - 41.740 | no |
| cSPH | CLIPB | CLIPB36 | AGAP013184 | ZOI | positive | 31.901 | 32.36 | 35.805 | 33.355 | 28.050 - 38.660 | 17.426 | 13.105 | 15.204 | 15.245 | 9.877 - 20.610 | 35.66 | 30.22 | 32.53 | 32.803 | 26.030 - 39.580 | no |
| cSP | CLIPB | CLIPB41 | AGAP004149 | ZOI | positive | 31.901 | 32.36 | 35.805 | 33.355 | 28.050 - 38.660 | 17.426 | 13.105 | 15.204 | 15.245 | 9.877 - 20.610 | 32.91 | 27.79 | 31.15 | 30.615 | 24.150 - 37.080 | no |
| cSP | CLIPB | CLIPB44 | AGAP009220 | ZOI | positive | 31.901 | 32.36 | 35.805 | 33.355 | 28.050 - 38.660 | 17.426 | 13.105 | 15.204 | 15.245 | 9.877 - 20.610 | 29.26 | 30.86 | 28.86 | 29.661 | 27.030 - 32.290 | no |
| cSP | CLIPB | CLIPB46 | AGAP009849 | ZOI | positive | 31.901 | 32.36 | 35.805 | 33.355 | 28.050 - 38.660 | 17.426 | 13.105 | 15.204 | 15.245 | 9.877 - 20.610 | 31.2 | 35.54 | 29.17 | 31.970 | 23.880 - 40.060 | no |

| | | | | | | | | | | | | | | | | | | | | | |
|---|---|---|---|---|---|---|---|---|---|---|---|---|---|---|---|---|---|---|---|---|---|
| cSP | CLIPB | CLIPB47 | AGAP003686 | ZOI | positive | 31.901 | 32.36 | 35.805 | 33.355 | 28.050 - 38.660 | 17.426 | 13.105 | 15.204 | 15.245 | 9.877 - 20.610 | 35.8 | 33.09 | 33.02 | 33.970 | 30.030 - 37.910 | no |
| cSP | CLIPC | CLIPC1 | AGAP008835 | ZOI | positive | 33.799 | 35.92 | 30.816 | 33.512 | 29.140 - 41.880 | 19.012 | 16.212 | 17.164 | 17.463 | 13.930 - 21.000 | 32.82 | 31.22 | 35.46 | 33.164 | 27.840 - 38.490 | no |
| cSP | CLIPC | CLIPC2 | AGAP004317 | ZOI | positive | 34.073 | 33.706 | 33.895 | 33.891 | 33.440 - 34.350 | 14.611 | 18.889 | 17.587 | 17.029 | 11.580 - 22.480 | 9.903 | 12.02 | 13.18 | 11.701 | 7.571 - 15.830 | yes |
| cSP | CLIPC | CLIPC3 | AGAP004318 | ZOI | positive | 36.106 | 36.816 | 32.499 | 35.140 | 29.390 - 40.890 | 18.140 | 18.661 | 17.835 | 18.212 | 17.170 - 19.250 | 30.35 | 38.27 | 37.13 | 35.248 | 24.620 - 45.880 | no |
| cSP | CLIPC | CLIPC4 | AGAP000573 | ZOI | positive | 34.073 | 33.706 | 33.895 | 33.891 | 33.440 - 34.350 | 14.611 | 18.889 | 17.587 | 17.029 | 11.580 - 22.480 | 10.22 | 11.9 | 12.53 | 11.550 | 8.586 - 14.510 | yes |
| cSP | CLIPC | CLIPC5 | AGAP000571 | ZOI | positive | 34.073 | 33.706 | 33.895 | 33.891 | 33.440 - 34.350 | 14.611 | 18.889 | 17.587 | 17.029 | 11.580 - 22.480 | 32.28 | 32.23 | 32.44 | 32.315 | 32.040 - 32.590 | no |
| cSP | CLIPC | CLIPC6 | AGAP000315 | ZOI | positive | 34.073 | 33.706 | 33.895 | 33.891 | 33.440 - 34.350 | 14.611 | 18.889 | 17.587 | 17.029 | 11.580 - 22.480 | 31.67 | 37.26 | 35.79 | 34.907 | 27.700 - 42.120 | no |
| cSP | CLIPC | CLIPC7 | AGAP003689 | ZOI | positive | 34.073 | 33.706 | 33.895 | 33.891 | 33.440 - 34.350 | 14.611 | 18.889 | 17.587 | 17.029 | 11.580 - 22.480 | 35.13 | 37.14 | 34.59 | 35.620 | 32.290 - 38.950 | no |
| cSP | CLIPC | CLIPC9 | AGAP004719 | ZOI | positive | 34.073 | 33.706 | 33.895 | 33.891 | 33.440 - 34.350 | 14.611 | 18.889 | 17.587 | 17.029 | 11.580 - 22.480 | 29.63 | 37.24 | 34.39 | 33.754 | 24.190 - 43.320 | no |
| cSP | CLIPC | CLIPC10 | AGAP000572 | ZOI | positive | 34.073 | 33.706 | 33.895 | 33.891 | 33.440 - 34.350 | 14.611 | 18.889 | 17.587 | 17.029 | 11.580 - 22.480 | 35.35 | 27.92 | 35.08 | 32.780 | 22.320 - 43.240 | no |
| cSP | CLIPC | CLIPC12 | AGAP012034 | ZOI | positive | 34.073 | 33.706 | 33.895 | 33.891 | 33.440 - 34.350 | 14.611 | 18.889 | 17.587 | 17.029 | 11.580 - 22.480 | 34.24 | 28.46 | 34.99 | 32.562 | 23.690 - 41.430 | no |
| cSP | CLIPC | CLIPC14 | AGAP028167 | ZOI | positive | 34.073 | 33.706 | 33.895 | 33.891 | 33.440 - 34.350 | 14.611 | 18.889 | 17.587 | 17.029 | 11.580 - 22.480 | 33.19 | 31.7 | 33.71 | 32.864 | 30.280 - 35.450 | no |
| cSP | CLIPD | CLIPD1 | AGAP002422 | ZOI | positive | 34.84 | 34.732 | 33.571 | 34.381 | 32.630 - 36.130 | 16.473 | 19.127 | 15.678 | 17.093 | 12.610 - 21.580 | 31.44 | 35.89 | 32.61 | 33.314 | 27.580 - 39.050 | no |
| cSP | CLIPD | CLIPD2 | AGAP008183 | ZOI | positive | 34.84 | 34.732 | 33.571 | 34.381 | 32.630 - 36.130 | 16.473 | 19.127 | 15.678 | 17.093 | 12.610 - 21.580 | 32.04 | 30.67 | 31.5 | 31.399 | 29.690 - 33.110 | no |
| cSP | CLIPD | CLIPD3 | AGAP001433 | ZOI | positive | 36.106 | 36.816 | 32.499 | 35.140 | 29.390 - 40.890 | 18.140 | 18.661 | 17.835 | 18.212 | 17.170 - 19.250 | 20.18 | 21.86 | 19.89 | 20.642 | 18.000 - 23.280 | yes |
| cSP | CLIPD | CLIPD4 | AGAP002811 | ZOI | positive | 34.84 | 34.732 | 33.571 | 34.381 | 32.630 - 36.130 | 16.473 | 19.127 | 15.678 | 17.093 | 12.610 - 21.580 | 35.99 | 35.27 | 33.35 | 34.869 | 31.480 - 38.250 | no |
| cSP | CLIPD | CLIPD6 | AGAP002813 | ZOI | positive | 34.84 | 34.732 | 33.571 | 34.381 | 32.630 - 36.130 | 16.473 | 19.127 | 15.678 | 17.093 | 12.610 - 21.580 | 38.01 | 36.93 | 32.36 | 35.767 | 28.320 - 43.210 | no |
| cSP | CLIPD | CLIPD7 | AGAP008998 | ZOI | positive | 34.84 | 34.732 | 33.571 | 34.381 | 32.630 - 36.130 | 16.473 | 19.127 | 15.678 | 17.093 | 12.610 - 21.580 | 35.05 | 29.23 | 34.58 | 32.953 | 24.920 - 40.990 | no |
| cSP | CLIPD | CLIPD8 | AGAP002784 | ZOI | positive | 34.84 | 34.732 | 33.571 | 34.381 | 32.630 - 36.130 | 16.473 | 19.127 | 15.678 | 17.093 | 12.610 - 21.580 | 35.4 | 35.02 | 34.45 | 34.955 | 33.760 - 36.150 | no |
| cSP | CLIPD | CLIPD9 | AGAP008997 | ZOI | positive | 34.84 | 34.732 | 33.571 | 34.381 | 32.630 - 36.130 | 16.473 | 19.127 | 15.678 | 17.093 | 12.610 - 21.580 | 36.61 | 32.1 | 35.76 | 34.822 | 28.870 - 40.770 | no |
| cSP | CLIPD | CLIPD11 | AGAP029106 | ZOI | positive | 34.84 | 34.732 | 33.571 | 34.381 | 33.630 - 37.130 | 16.473 | 19.127 | 15.678 | 17.093 | 12.610 - 21.580 | 33.81 | 30.35 | 30.86 | 31.673 | 27.020 - 36.320 | no |
| cSP | CLIPD | CLIPD12 | AGAP008995 | ZOI | positive | 33.799 | 35.92 | 30.816 | 33.512 | 27.140 - 39.880 | 19.012 | 16.212 | 17.164 | 17.463 | 13.930 - 21.000 | 32.41 | 33.35 | 31.51 | 32.423 | 30.140 - 34.710 | no |
| cSP | CLIPD | CLIPD13 | AGAP009000 | ZOI | positive | 33.799 | 35.92 | 30.816 | 33.512 | 27.140 - 39.880 | 19.012 | 16.212 | 17.164 | 17.463 | 13.930 - 21.000 | 31.27 | 30.82 | 32.29 | 31.459 | 29.600 - 33.320 | no |
| cSP | CLIPD | CLIPD14 | AGAP009006 | ZOI | positive | 33.799 | 35.92 | 30.816 | 33.512 | 27.140 - 39.880 | 19.012 | 16.212 | 17.164 | 17.463 | 13.930 - 21.000 | 31.91 | 33.16 | 34.51 | 33.193 | 29.950 - 36.430 | no |
| cSP | CLIPD | CLIPD20 | AGAP013089 | ZOI | positive | 33.799 | 35.92 | 30.816 | 33.512 | 27.140 - 39.880 | 19.012 | 16.212 | 17.164 | 17.463 | 13.930 - 21.000 | 31.48 | 29.13 | 33.1 | 31.234 | 26.270 - 36.190 | no |
| cSP | CLIPD | CLIPD22 | AGAP008996 | ZOI | positive | 33.799 | 35.92 | 30.816 | 33.512 | 27.140 - 39.880 | 19.012 | 16.212 | 17.164 | 17.463 | 13.930 - 21.000 | 31.01 | 30.16 | 32.62 | 31.263 | 28.150 - 34.370 | no |
| cSPH | CLIPE | CLIPE1 | AGAP008091 | ZOI | positive | 33.814 | 32.732 | 33.571 | 33.372 | 31.960 - 34.780 | 16.274 | 18.154 | 18.218 | 17.549 | 14.810 - 20.290 | 31.25 | 34.27 | 34.22 | 33.243 | 28.950 - 37.530 | no |
| cSPH | CLIPE | CLIPE4 | AGAP010530 | ZOI | positive | 33.814 | 32.732 | 33.571 | 33.372 | 31.960 - 34.780 | 16.274 | 18.154 | 18.218 | 17.549 | 14.810 - 20.290 | 28.4 | 34.18 | 31.28 | 31.286 | 24.100 - 38.470 | no |
| cSP | CLIPE | CLIPE5 | AGAP028728 | ZOI | positive | 33.814 | 32.732 | 33.571 | 33.372 | 31.960 - 34.780 | 16.274 | 18.154 | 18.218 | 17.549 | 14.810 - 20.290 | 30.49 | 28.46 | 34.55 | 31.168 | 23.470 - 38.860 | no |
| cSPH | CLIPE | CLIPE6 | AGAP011785 | ZOI | positive | 33.814 | 32.732 | 33.571 | 33.372 | 31.960 - 34.780 | 16.274 | 18.154 | 18.218 | 17.549 | 14.810 - 20.290 | 33.15 | 29.21 | 35.32 | 32.561 | 24.870 - 40.250 | no |
| cSPH | CLIPE | CLIPE7 | AGAP011786 | ZOI | positive | 33.814 | 32.732 | 33.571 | 33.372 | 31.960 - 34.780 | 16.274 | 18.154 | 18.218 | 17.549 | 14.810 - 20.290 | 36.01 | 34.52 | 33.64 | 34.725 | 31.760 - 37.690 | no |
| cSP | CLIPE | CLIPE10 | AGAP010545 | ZOI | positive | 33.814 | 32.732 | 33.571 | 33.372 | 31.960 - 34.780 | 16.274 | 18.154 | 18.218 | 17.549 | 14.810 - 20.290 | 30.34 | 29.01 | 29.65 | 29.671 | 28.020 - 31.320 | no |
| cSPH | CLIPE | CLIPE12 | AGAP003691 | ZOI | positive | 33.814 | 32.732 | 33.571 | 33.372 | 31.960 - 34.780 | 16.274 | 18.154 | 18.218 | 17.549 | 14.810 - 20.290 | 29.44 | 31.35 | 32.35 | 31.047 | 27.370 - 34.720 | no |
| cSP | CLIPE | CLIPE13 | AGAP003748 | ZOI | positive | 31.048 | 33.953 | 29.241 | 31.414 | 25.510 - 37.320 | 17.947 | 19.059 | 18.135 | 18.380 | 16.900 - 19.860 | 29.28 | 30.45 | 35.22 | 31.653 | 23.840 - 39.470 | no |
| cSP | CLIPE | CLIPE15 | AGAP009249 | ZOI | positive | 31.048 | 33.953 | 29.241 | 31.414 | 25.510 - 37.320 | 17.947 | 19.059 | 18.135 | 18.380 | 16.900 - 19.860 | 29.5 | 30.28 | 31.54 | 30.441 | 27.870 - 33.010 | no |
| cSPH | CLIPE | CLIPE17 | AGAP009252 | ZOI | positive | 31.048 | 33.953 | 29.241 | 31.414 | 25.510 - 37.320 | 17.947 | 19.059 | 18.135 | 18.380 | 16.900 - 19.860 | 30.4 | 28.94 | 33.57 | 30.970 | 25.100 - 36.840 | no |
| cSP | CLIPE | CLIPE18 | AGAP009273 | ZOI | positive | 31.048 | 33.953 | 29.241 | 31.414 | 25.510 - 37.320 | 17.947 | 19.059 | 18.135 | 18.380 | 16.900 - 19.860 | 35.31 | 30.39 | 32.35 | 32.682 | 26.530 - 38.830 | no |
| cSP | CLIPE | CLIPE19 | AGAP011040 | ZOI | positive | 31.048 | 33.953 | 29.241 | 31.414 | 25.510 - 37.320 | 17.947 | 19.059 | 18.135 | 18.380 | 16.900 - 19.860 | 32.86 | 32.4 | 34.14 | 33.133 | 30.900 - 35.370 | no |
| cSP | CLIPE | CLIPE21 | AGAP011719 | ZOI | positive | 31.048 | 33.953 | 29.241 | 31.414 | 25.510 - 37.320 | 17.947 | 19.059 | 18.135 | 18.380 | 16.900 - 19.860 | 32.06 | 31.09 | 29.14 | 30.759 | 27.070 - 34.450 | no |
| cSP | CLIPE | CLIPE22 | AGAP012020 | ZOI | positive | 31.048 | 33.953 | 29.241 | 31.414 | 25.510 - 37.320 | 17.947 | 19.059 | 18.135 | 18.380 | 16.900 - 19.860 | 35.06 | 35.37 | 35.14 | 35.188 | 34.780 - 35.590 | no |
| cSP | CLIPE | CLIPE25 | AGAP012502 | ZOI | positive | 31.048 | 33.953 | 29.241 | 31.414 | 25.510 - 37.320 | 17.947 | 19.059 | 18.135 | 18.380 | 16.900 - 19.860 | 31.96 | 29.79 | 34.93 | 32.227 | 25.820 - 38.630 | no |
| cSP | CLIPE | CLIPE30 | AGAP010628 | ZOI | positive | 31.048 | 33.953 | 29.241 | 31.414 | 25.510 - 37.320 | 17.947 | 19.059 | 18.135 | 18.380 | 16.900 - 19.860 | 29.92 | 34.49 | 30.2 | 31.533 | 25.170 - 37.900 | no |

**Table S6: ZOI screen: Zone of inhibition of genes tested as negative regulators.** DsRNA injected mosquitoes were challenged with 50.6 nL of resuspended lyophilized *Micrococcus luteus* at $OD_{600}$ = 0.1. Batches of 7- 10 candidate genes were tested in independent groups using their own positive (ds*CACT*) and negative controls. Three independent biological replicates were performed using different mosquito generations.

| CLIP family | sub family | Gene name | AGAP | Assay type | Tested as | ds*GFP* controls (individual replicates) | | | | | Positive controls-ds*CACT* (individual replicates) | | | | | ds*GOI* (individual replicates) | | | | | Phenotype |
|---|---|---|---|---|---|---|---|---|---|---|---|---|---|---|---|---|---|---|---|---|---|
| | | | | | | 1 | 2 | 3 | Mean | 95% CI of mean | 1 | 2 | 3 | Mean | 95% CI of mean | 1 | 2 | 3 | Mean | 95% CI of mean | |
| cSPH | CLIPA | CLIPA1 | AGAP011791 | ZOI | negative | 11.203 | 13.255 | 10.304 | 11.587 | 7.830 - 15.340 | 31.666 | 33.036 | 34.407 | 33.036 | 29.630 - 36.440 | 12.948 | 13.463 | 14.079 | 13.497 | 12.090 - 14.900 | no |
| cSPH | CLIPA | CLIPA2 | AGAP011790 | ZOI | negative | 8.665 | 11.315 | 11.921 | 10.634 | 6.332 - 14.940 | 26.561 | 27.584 | 31.596 | 28.580 | 21.970 - 35.190 | 8.032 | 11.682 | 10.362 | 10.025 | 5.434 - 14.620 | no |
| cSPH | CLIPA | CLIPA3 | AGAP012591 | ZOI | negative | 8.665 | 11.315 | 11.921 | 10.634 | 6.332 - 14.940 | 26.561 | 27.584 | 31.596 | 28.580 | 21.970 - 35.190 | 9.187 | 10.617 | 10.146 | 9.983 | 8.173 - 11.790 | no |
| cSPH | CLIPA | CLIPA4 | AGAP011780 | ZOI | negative | 8.665 | 11.315 | 11.921 | 10.634 | 6.332 - 14.940 | 26.561 | 27.584 | 31.596 | 28.580 | 21.970 - 35.190 | 15.046 | 19.054 | 19.054 | 17.718 | 11.970 - 23.470 | yes |
| cSPH | CLIPA | CLIPA5 | AGAP011787 | ZOI | negative | 8.665 | 11.315 | 11.921 | 10.634 | 6.332 - 14.940 | 26.561 | 27.584 | 31.596 | 28.580 | 21.970 - 35.190 | 8.784 | 9.953 | 13.803 | 10.847 | 4.323 - 17.370 | no |
| cSPH | CLIPA | CLIPA6 | AGAP011789 | ZOI | negative | 8.665 | 11.315 | 11.921 | 10.634 | 6.332 - 14.940 | 26.561 | 27.584 | 31.596 | 28.580 | 21.970 - 35.190 | 10.887 | 11.639 | 12.210 | 11.579 | 9.930 - 13.230 | no |
| cSPH | CLIPA | CLIPA7 | AGAP011792 | ZOI | negative | 8.665 | 11.315 | 11.921 | 10.634 | 6.332 - 14.940 | 26.561 | 27.584 | 31.596 | 28.580 | 21.970 - 35.190 | 10.698 | 9.849 | 12.550 | 11.032 | 7.601 - 14.460 | no |
| cSPH | CLIPA | CLIPA8 | AGAP010731 | ZOI | negative | 8.665 | 11.315 | 11.921 | 10.634 | 6.332 - 14.940 | 26.561 | 27.584 | 31.596 | 28.580 | 21.970 - 35.190 | 10.189 | 11.350 | 12.202 | 11.247 | 8.737 - 13.760 | no |
| cSPH | CLIPA | CLIPA9 | AGAP010968 | ZOI | negative | 8.665 | 11.315 | 11.921 | 10.634 | 6.332 - 14.940 | 26.561 | 27.584 | 31.596 | 28.580 | 21.970 - 35.190 | 10.119 | 11.836 | 11.304 | 11.086 | 8.903 - 13.270 | no |
| cSPH | CLIPA | CLIPA10 | AGAP006954 | ZOI | negative | 8.665 | 11.315 | 11.921 | 10.634 | 6.332 - 14.940 | 26.561 | 27.584 | 31.596 | 28.580 | 21.970 - 35.190 | 10.733 | 8.865 | 11.936 | 10.511 | 6.667 - 14.360 | no |
| cSPH | CLIPA | CLIPA12 | AGAP011781 | ZOI | negative | 8.665 | 11.315 | 11.921 | 10.634 | 6.332 - 14.940 | 26.561 | 27.584 | 31.596 | 28.580 | 21.970 - 35.190 | 11.126 | 10.409 | 13.260 | 11.598 | 7.914 - 15.280 | no |
| cSPH | CLIPA | CLIPA13 | AGAP011783 | ZOI | negative | 9.068 | 10.085 | 11.167 | 10.107 | 7.499 - 12.710 | 27.274 | 29.486 | 30.015 | 28.925 | 25.310 - 32.540 | 7.254 | 9.213 | 10.285 | 8.917 | 5.099 - 12.740 | no |
| cSPH | CLIPA | CLIPA14 | AGAP011788 | ZOI | negative | 9.068 | 10.085 | 11.167 | 10.107 | 7.499 - 12.710 | 27.274 | 29.486 | 30.015 | 28.925 | 25.310 - 32.540 | 8.315 | 8.514 | 9.209 | 8.679 | 7.513 - 9.845 | no |
| cSPH | CLIPA | CLIPA15 | AGAP002815 | ZOI | negative | 9.068 | 10.085 | 11.167 | 10.107 | 7.499 - 12.710 | 27.274 | 29.486 | 30.015 | 28.925 | 25.310 - 32.540 | 8.197 | 11.017 | 11.049 | 10.088 | 6.020 - 14.160 | no |
| cSPH | CLIPA | CLIPA19 | AGAP003245 | ZOI | negative | 9.068 | 10.085 | 11.167 | 10.107 | 7.499 - 12.710 | 27.274 | 29.486 | 30.015 | 28.925 | 25.310 - 32.540 | 14.088 | 15.284 | 13.955 | 14.442 | 12.620 - 16.260 | yes |
| cSPH | CLIPA | CLIPA26 | AGAP001964 | ZOI | negative | 9.068 | 10.085 | 11.167 | 10.107 | 7.499 - 12.710 | 27.274 | 29.486 | 30.015 | 28.925 | 25.310 - 32.540 | 10.319 | 9.001 | 11.208 | 10.176 | 7.418 - 12.930 | no |
| cSPH | CLIPA | CLIPA27 | AGAP000290 | ZOI | negative | 9.068 | 10.085 | 11.167 | 10.107 | 7.499 - 12.710 | 27.274 | 29.486 | 30.015 | 28.925 | 25.310 - 32.540 | 9.279 | 9.106 | 11.138 | 9.841 | 7.042 - 12.640 | no |
| cSPH | CLIPA | CLIPA28 | AGAP010730 | ZOI | negative | 9.068 | 10.085 | 11.167 | 10.107 | 7.499 - 12.710 | 27.274 | 29.486 | 30.015 | 28.925 | 25.310 - 32.540 | 8.384 | 11.297 | 10.084 | 9.922 | 6.287 - 13.560 | no |
| cSPH | CLIPA | SPCLIP1 | AGAP028725 | ZOI | negative | 9.068 | 10.085 | 11.167 | 10.107 | 7.499 - 12.710 | 27.274 | 29.486 | 30.015 | 28.925 | 25.310 - 32.540 | 10.315 | 11.471 | 11.297 | 11.028 | 9.479 - 12.580 | no |
| cSPH | CLIPB | CLIPB7 | AGAP002270 | ZOI | negative | 7.415 | 11.431 | 7.002 | 8.616 | 2.538 - 14.690 | 27.985 | 29.505 | 28.021 | 28.504 | 26.350 - 30.660 | 8.663 | 9.741 | 9.182 | 9.862 | 7.856 - 10.530 | no |
| cSPH | CLIPB | CLIPB12 | AGAP009217 | ZOI | negative | 7.415 | 11.431 | 7.002 | 8.616 | 3.538 - 15.690 | 27.985 | 29.505 | 28.021 | 28.504 | 26.350 - 30.660 | 8.534 | 8.252 | 7.577 | 8.121 | 6.899 - 9.343 | no |
| cSPH | CLIPB | CLIPB16 | AGAP009263 | ZOI | negative | 7.415 | 11.431 | 7.002 | 8.616 | 4.538 - 16.690 | 27.985 | 29.505 | 28.021 | 28.504 | 26.350 - 30.660 | 7.214 | 11.473 | 6.543 | 8.410 | 1.768 - 15.050 | no |
| cSPH | CLIPB | CLIPB36 | AGAP013184 | ZOI | negative | 7.415 | 11.431 | 7.002 | 8.616 | 5.538 - 17.690 | 27.985 | 29.505 | 28.021 | 28.504 | 26.350 - 30.660 | 8.580 | 9.197 | 6.786 | 8.188 | 5.076 - 11.300 | no |
| cSPH | CLIPC | CLIPC7 | AGAP003689 | ZOI | negative | 11.203 | 13.255 | 10.304 | 11.587 | 7.830 - 15.340 | 31.666 | 33.036 | 34.407 | 33.036 | 29.630 - 36.440 | 9.957 | 11.547 | 11.246 | 10.917 | 8.819 - 13.010 | no |
| cSPH | CLIPD | CLIPD14 | AGAP009006 | ZOI | negative | 11.203 | 13.255 | 10.304 | 11.587 | 7.830 - 15.340 | 31.666 | 33.036 | 34.407 | 33.036 | 29.630 - 36.440 | 14.012 | 12.854 | 9.726 | 12.197 | 6.690 - 17.710 | no |
| cSPH | CLIPE | CLIPE1 | AGAP008091 | ZOI | negative | 11.203 | 13.255 | 10.304 | 11.587 | 7.830 - 15.340 | 31.666 | 33.036 | 34.407 | 33.036 | 29.630 - 36.440 | 12.984 | 11.174 | 12.695 | 12.284 | 9.869 - 14.700 | no |
| cSPH | CLIPE | CLIPE4 | AGAP010530 | ZOI | negative | 11.203 | 13.255 | 10.304 | 11.587 | 7.830 - 15.340 | 31.666 | 33.036 | 34.407 | 33.036 | 29.630 - 36.440 | 11.465 | 12.127 | 9.627 | 11.073 | 7.855 - 14.290 | no |
| cSPH | CLIPE | CLIPE6 | AGAP011785 | ZOI | negative | 11.203 | 13.255 | 10.304 | 11.587 | 7.830 - 15.340 | 31.666 | 33.036 | 34.407 | 33.036 | 29.630 - 36.440 | 12.217 | 11.370 | 11.373 | 11.653 | 10.440 - 12.870 | no |
| cSPH | CLIPE | CLIPE7 | AGAP011786 | ZOI | negative | 11.203 | 13.255 | 10.304 | 11.587 | 7.830 - 15.340 | 31.666 | 33.036 | 34.407 | 33.036 | 29.630 - 36.440 | 9.018 | 12.811 | 11.769 | 11.199 | 6.331 - 16.070 | no |
| cSPH | CLIPE | CLIPE12 | AGAP003691 | ZOI | negative | 11.203 | 13.255 | 10.304 | 11.587 | 7.830 - 15.340 | 31.666 | 33.036 | 34.407 | 33.036 | 29.630 - 36.440 | 13.654 | 11.552 | 11.071 | 12.092 | 8.680 - 15.500 | no |
| cSPH | CLIPE | CLIPE17 | AGAP009252 | ZOI | negative | 11.203 | 13.255 | 10.304 | 11.587 | 7.830 - 15.340 | 31.666 | 33.036 | 34.407 | 33.036 | 29.630 - 36.440 | 10.283 | 10.928 | 11.854 | 11.022 | 9.060 - 12.980 | no |

**Table S7: Total melanotic spot area of positive regulators of melanization.** DsRNA injected mosquitoes were challenged with 50.6 nL of resuspended lyophilized *Micrococcus luteus* at OD600 = 5. *CLIPB4* kd was used as a positive control for positive regulation of antimicrobial activity. Five independent biological replicates were performed using different mosquito generations. One-way ANOVA followed by Bonferroni's post-tests were performed on total melanotic spot area data to calculate statistical significance ($P < 0.05$).

| CLIP family | sub family | Gene name | AGAP | Assay type | Tested as | ds*GFP* controls (individual replicates) | | | | | | | ds*GOI* (individual replicates) | | | | | | | *P* |
|---|---|---|---|---|---|---|---|---|---|---|---|---|---|---|---|---|---|---|---|---|
| | | | | | | 1 | 2 | 3 | 4 | 5 | Mean | 95% CI of mean | 1 | 2 | 3 | 4 | 5 | Mean | 95% CI of mean | |
| | | TEP1 | AGAP010815 | MelASA | positive | 0.184 | 0.202 | 0.198 | 0.187 | 0.182 | 0.189 | 0.180 - 0.198 | 0.071 | 0.082 | 0.064 | 0.069 | 0.078 | 0.073 | 0.064 - 0.082 | <0.0001 |
| | | MODSP1 | AGAP001798 | MelASA | positive | 0.194 | 0.204 | 0.197 | 0.198 | 0.187 | 0.196 | 0.188 - 0.204 | 0.145 | 0.153 | 0.139 | 0.149 | 0.141 | 0.145 | 0.138 - 0.153 | <0.0001 |
| cSPH | CLIPA | CLIPA8 | AGAP010731 | MelASA | positive | 0.194 | 0.203 | 0.194 | 0.184 | 0.202 | 0.195 | 0.186 - 0.205 | 0.057 | 0.079 | 0.084 | 0.062 | 0.073 | 0.071 | 0.057 - 0.085 | <0.0001 |
| cSPH | CLIPA | CLIPA28 | AGAP010730 | MelASA | positive | 0.193 | 0.199 | 0.191 | 0.184 | 0.202 | 0.194 | 0.185 - 0.203 | 0.119 | 0.117 | 0.125 | 0.122 | 0.123 | 0.121 | 0.117 - 0.125 | <0.0001 |
| cSPH | CLIPA | SPCLIP1 | AGAP028725 | MelASA | positive | 0.193 | 0.199 | 0.191 | 0.184 | 0.202 | 0.194 | 0.185 - 0.203 | 0.059 | 0.057 | 0.048 | 0.053 | 0.059 | 0.055 | 0.049 - 0.061 | <0.0001 |
| cSP | CLIPB | CLIPB8 | AGAP003057 | MelASA | positive | 0.194 | 0.204 | 0.197 | 0.184 | 0.202 | 0.196 | 0.186 - 0.206 | 0.135 | 0.158 | 0.132 | 0.147 | 0.154 | 0.145 | 0.131 - 0.159 | <0.0001 |
| cSP | CLIPB | CLIPB9 | AGAP029769 | MelASA | positive | 0.194 | 0.204 | 0.197 | 0.184 | 0.202 | 0.196 | 0.186 - 0.206 | 0.126 | 0.118 | 0.125 | 0.121 | 0.128 | 0.124 | 0.119 - 0.129 | <0.0001 |
| cSP | CLIPB | CLIPB10 | AGAP029770 | MelASA | positive | 0.194 | 0.204 | 0.197 | 0.184 | 0.202 | 0.196 | 0.186 - 0.206 | 0.101 | 0.110 | 0.105 | 0.104 | 0.109 | 0.106 | 0.101 - 0.110 | <0.0001 |
| cSP | CLIPB | CLIPB13 | AGAP004855 | MelASA | positive | 0.202 | 0.227 | 0.218 | 0.198 | 0.187 | 0.206 | 0.187 - 0.226 | 0.128 | 0.117 | 0.123 | 0.126 | 0.119 | 0.123 | 0.117 - 0.128 | <0.0001 |
| cSP | CLIPB | CLIPB17 | AGAP001648 | MelASA | positive | 0.191 | 0.182 | 0.191 | 0.198 | 0.187 | 0.190 | 0.183 - 0.197 | 0.128 | 0.124 | 0.136 | 0.132 | 0.122 | 0.128 | 0.121 - 0.136 | <0.0001 |
| cSP | CLIPB | CLIPB20 | AGAP012037 | MelASA | positive | 0.205 | 0.217 | 0.223 | 0.198 | 0.187 | 0.206 | 0.188 - 0.224 | 0.148 | 0.159 | 0.136 | 0.143 | 0.137 | 0.145 | 0.133 - 0.156 | <0.0001 |
| cSP | CLIPC | CLIPC3 | AGAP004318 | MelASA | positive | 0.205 | 0.217 | 0.223 | 0.198 | 0.187 | 0.206 | 0.188 - 0.224 | 0.125 | 0.133 | 0.142 | 0.139 | 0.135 | 0.135 | 0.127 - 0.143 | <0.0001 |
| cSP | CLIPC | CLIPC9 | AGAP004719 | MelASA | positive | 0.217 | 0.208 | 0.195 | 0.198 | 0.187 | 0.201 | 0.187 - 0.216 | 0.152 | 0.159 | 0.140 | 0.142 | 0.149 | 0.148 | 0.139 - 0.158 | <0.0001 |
| cSP | CLIPC | CLIPC12 | AGAP012034 | MelASA | positive | 0.217 | 0.208 | 0.195 | 0.198 | 0.187 | 0.201 | 0.187 - 0.216 | 0.139 | 0.123 | 0.141 | 0.134 | 0.126 | 0.133 | 0.123 - 0.142 | <0.0001 |
| cSP | CLIPC | CLIPC14 | AGAP028167 | MelASA | positive | 0.217 | 0.208 | 0.195 | 0.198 | 0.187 | 0.201 | 0.187 - 0.216 | 0.123 | 0.134 | 0.148 | 0.125 | 0.133 | 0.133 | 0.120 - 0.145 | <0.0001 |

| CLIP family | sub family | Gene name | AGAP | Assay type | Tested as | Positive controls-ds*CLIPB4* | | | | | | |
|---|---|---|---|---|---|---|---|---|---|---|---|---|
| | | | | | | 1 | 2 | 3 | 4 | 5 | Mean | 95% CI of mean |
| | | TEP1 | AGAP010815 | MelASA | positive | 0.139 | 0.128 | 0.133 | 0.126 | 0.121 | 0.138 | 0.115 - 0.162 |
| | | MODSP1 | AGAP001798 | MelASA | positive | 0.117 | 0.122 | 0.126 | 0.133 | 0.126 | 0.125 | 0.118 - 0.132 |
| cSPH | CLIPA | CLIPA8 | AGAP010731 | MelASA | positive | 0.137 | 0.136 | 0.104 | 0.139 | 0.128 | 0.129 | 0.111 - 0.147 |
| cSPH | CLIPA | CLIPA28 | AGAP010730 | MelASA | positive | 0.125 | 0.132 | 0.140 | 0.139 | 0.128 | 0.133 | 0.125 - 0.141 |
| cSPH | CLIPA | SPCLIP1 | AGAP028725 | MelASA | positive | 0.125 | 0.132 | 0.140 | 0.139 | 0.128 | 0.133 | 0.125 - 0.141 |
| cSP | CLIPB | CLIPB8 | AGAP003057 | MelASA | positive | 0.117 | 0.122 | 0.126 | 0.139 | 0.128 | 0.126 | 0.116 - 0.137 |
| cSP | CLIPB | CLIPB9 | AGAP029769 | MelASA | positive | 0.117 | 0.122 | 0.126 | 0.139 | 0.128 | 0.126 | 0.116 - 0.137 |
| cSP | CLIPB | CLIPB10 | AGAP029770 | MelASA | positive | 0.117 | 0.122 | 0.126 | 0.139 | 0.128 | 0.126 | 0.116 - 0.137 |
| cSP | CLIPB | CLIPB13 | AGAP004855 | MelASA | positive | 0.110 | 0.136 | 0.125 | 0.133 | 0.126 | 0.126 | 0.114 - 0.139 |
| cSP | CLIPB | CLIPB17 | AGAP001648 | MelASA | positive | 0.124 | 0.126 | 0.114 | 0.133 | 0.126 | 0.125 | 0.116 - 0.133 |
| cSP | CLIPB | CLIPB20 | AGAP012037 | MelASA | positive | 0.122 | 0.119 | 0.135 | 0.133 | 0.126 | 0.127 | 0.118 - 0.136 |
| cSP | CLIPC | CLIPC3 | AGAP004318 | MelASA | positive | 0.122 | 0.119 | 0.135 | 0.133 | 0.126 | 0.127 | 0.118 - 0.136 |
| cSP | CLIPC | CLIPC9 | AGAP004719 | MelASA | positive | 0.111 | 0.124 | 0.119 | 0.133 | 0.126 | 0.123 | 0.112 - 0.133 |
| cSP | CLIPC | CLIPC12 | AGAP012034 | MelASA | positive | 0.111 | 0.124 | 0.119 | 0.133 | 0.126 | 0.123 | 0.112 - 0.133 |
| cSP | CLIPC | CLIPC14 | AGAP028167 | MelASA | positive | 0.111 | 0.124 | 0.119 | 0.133 | 0.126 | 0.123 | 0.112 - 0.133 |

**Table S8: Total melanotic spot area of negative regulators of melanization.** DsRNA injected mosquitoes were challenged with 50.6 nL of resuspended lyophilized *Micrococcus luteus* at $OD_{600}$ =0.1. *SRPN2* kd was used as a positive control for positive regulation of antimicrobial activity. Five independent biological replicates were performed using different mosquito generations. One-way ANOVA followed by Bonferroni's post-tests were performed on total melanotic spot area data to calculate statistical significance (*P*< 0.05).

| CLIP family | sub family | Gene name | AGAP | Assay type | Tested as | ds*GFP* controls (individual replicates) | | | | | | | ds*GOI* (individual replicates) | | | | | | | *P* |
|---|---|---|---|---|---|---|---|---|---|---|---|---|---|---|---|---|---|---|---|---|
| | | | | | | 1 | 2 | 3 | 4 | 5 | Mean | 95% CI of mean | 1 | 2 | 3 | 4 | 5 | Mean | 95% CI of mean | |
| cSPH | CLIPA | CLIPA2 | AGAP011790 | MelASA | negative | 0.037 | 0.043 | 0.039 | 0.042 | 0.034 | 0.063 | 0.001 - 0.124 | 0.099 | 0.105 | 0.117 | 0.108 | 0.113 | 0.108 | 0.100 - 0.117 | <0.0001 |
| cSPH | CLIPA | CLIPA5 | AGAP011787 | MelASA | negative | 0.037 | 0.043 | 0.039 | 0.042 | 0.034 | 0.039 | 0.034 - 0.044 | 0.079 | 0.092 | 0.106 | 0.086 | 0.101 | 0.093 | 0.079 - 0.106 | <0.0001 |
| cSPH | CLIPA | CLIPA7 | AGAP011792 | MelASA | negative | 0.037 | 0.043 | 0.039 | 0.042 | 0.034 | 0.039 | 0.034 - 0.044 | 0.064 | 0.078 | 0.091 | 0.083 | 0.069 | 0.077 | 0.064 - 0.090 | <0.0001 |
| cSPH | CLIPA | CLIPA13 | AGAP011783 | MelASA | negative | 0.032 | 0.047 | 0.041 | 0.042 | 0.034 | 0.039 | 0.032 - 0.047 | 0.044 | 0.068 | 0.061 | 0.067 | 0.071 | 0.066 | 0.061 - 0.071 | <0.0001 |
| cSPH | CLIPA | CLIPA14 | AGAP011788 | MelASA | negative | 0.032 | 0.047 | 0.041 | 0.042 | 0.034 | 0.039 | 0.032 - 0.047 | 0.102 | 0.099 | 0.097 | 0.106 | 0.095 | 0.100 | 0.094 - 0.105 | <0.0001 |
| cSPH | CLIPB | CLIPB12 | AGAP009217 | MelASA | negative | 0.043 | 0.055 | 0.038 | 0.042 | 0.034 | 0.042 | 0.033 - 0.052 | 0.067 | 0.078 | 0.086 | 0.072 | 0.064 | 0.073 | 0.062 - 0.084 | <0.0001 |

| CLIP family | sub family | Gene name | AGAP | Assay type | Tested as | (individual replicates) | | | | | | |
|---|---|---|---|---|---|---|---|---|---|---|---|---|
| | | | | | | 1 | 2 | 3 | 4 | 5 | Mean | 95% CI of mean |
| cSPH | CLIPA | CLIPA2 | AGAP011790 | MelASA | negative | 0.138 | 0.127 | 0.149 | 0.135 | 0.119 | 0.134 | 0.120 - 0.148 |
| cSPH | CLIPA | CLIPA5 | AGAP011787 | MelASA | negative | 0.138 | 0.127 | 0.149 | 0.135 | 0.119 | 0.134 | 0.120 - 0.148 |
| cSPH | CLIPA | CLIPA7 | AGAP011792 | MelASA | negative | 0.138 | 0.127 | 0.149 | 0.135 | 0.119 | 0.134 | 0.120 - 0.148 |
| cSPH | CLIPA | CLIPA13 | AGAP011783 | MelASA | negative | 0.127 | 0.114 | 0.131 | 0.135 | 0.119 | 0.125 | 0.115 - 0.136 |
| cSPH | CLIPA | CLIPA14 | AGAP011788 | MelASA | negative | 0.127 | 0.114 | 0.131 | 0.135 | 0.119 | 0.125 | 0.115 - 0.136 |
| cSPH | CLIPB | CLIPB12 | AGAP009217 | MelASA | negative | 0.147 | 0.143 | 0.131 | 0.135 | 0.119 | 0.135 | 0.121 - 0.149 |

**Table S9: Zone of Inhibition of positive regulators of antimicrobial activity.** DsRNA injected mosquitoes were challenged with 50.6 nL of resuspended lyophilized *Micrococcus luteus* at OD600= 5. *REL1* kd was used as a positive control for positive regulation of antimicrobial activity. Five independent biological replicates were performed using different mosquito generations. One-way ANOVA followed by Bonferroni's post-tests were performed on ZOI data to calculate statistical significance ($P < 0.05$).

| CLIP family | sub family | Gene name | AGAP | Assay type | Tested as | ds*GFP* controls (individual replicates) | | | | | | | ds*GOI* (individual replicates) | | | | | | | *P* |
|---|---|---|---|---|---|---|---|---|---|---|---|---|---|---|---|---|---|---|---|---|
| | | | | | | 1 | 2 | 3 | 4 | 5 | Mean | 95% CI of mean | 1 | 2 | 3 | 4 | 5 | Mean | 95% CI of mean | |
| cSPH | CLIPA | CLIPA3 | AGAP012591 | ZOI | positive | 31.951 | 34.689 | 35.026 | 34.219 | 35.241 | 34.230 | 32.580 - 35.870 | 14.590 | 19.625 | 15.161 | 16.334 | 18.945 | 16.930 | 14.130 - 19.730 | <0.0001 |
| cSP | CLIPB | CLIPB10 | AGAP029770 | ZOI | positive | 33.799 | 35.920 | 30.816 | 34.219 | 35.241 | 34.000 | 31.560 - 36.440 | 14.158 | 16.016 | 15.375 | 14.732 | 15.227 | 15.100 | 14.230 - 15.970 | <0.0001 |
| cSP | CLIPB | CLIPB14 | AGAP010833 | ZOI | positive | 31.901 | 32.360 | 35.805 | 34.219 | 35.241 | 33.910 | 31.760 - 36.050 | 10.933 | 14.374 | 13.530 | 14.945 | 11.067 | 12.970 | 10.650 - 15.290 | <0.0001 |
| cSP | CLIPC | CLIPC2 | AGAP004317 | ZOI | positive | 34.073 | 33.706 | 33.895 | 34.219 | 35.241 | 34.230 | 33.480 - 34.970 | 9.903 | 12.017 | 13.182 | 10.168 | 10.983 | 11.250 | 9.564 - 12.940 | <0.0001 |
| cSP | CLIPC | CLIPC4 | AGAP000573 | ZOI | positive | 34.073 | 33.706 | 33.895 | 34.219 | 35.241 | 34.230 | 33.480 - 34.970 | 10.220 | 11.902 | 12.527 | 10.386 | 12.954 | 11.600 | 10.060 - 13.140 | <0.0001 |
| cSP | CLIPD | CLIPD3 | AGAP001433 | ZOI | positive | 36.106 | 36.816 | 32.499 | 34.219 | 35.241 | 34.980 | 32.880 - 37.080 | 20.177 | 21.859 | 19.891 | 19.344 | 16.253 | 19.500 | 16.960 - 22.050 | <0.0001 |

| CLIP family | sub family | Gene name | AGAP | Assay type | Tested as | Positive controls-ds*REL1* | | | | | | |
|---|---|---|---|---|---|---|---|---|---|---|---|---|
| | | | | | | 1 | 2 | 3 | 4 | 5 | Mean | 95% CI of mean |
| cSPH | CLIPA | CLIPA3 | AGAP012591 | ZOI | positive | 15.192 | 12.623 | 16.820 | 14.872 | 18.69 | 15.640 | 12.820 - 18.460 |
| cSP | CLIPB | CLIPB10 | AGAP029770 | ZOI | positive | 19.012 | 16.212 | 17.164 | 14.872 | 18.69 | 17.190 | 15.050 - 19.330 |
| cSP | CLIPB | CLIPB14 | AGAP010833 | ZOI | positive | 17.426 | 13.105 | 15.204 | 14.872 | 18.69 | 15.860 | 13.120 - 18.600 |
| cSP | CLIPC | CLIPC2 | AGAP004317 | ZOI | positive | 14.611 | 18.889 | 17.587 | 14.872 | 18.69 | 16.930 | 14.370 - 19.490 |
| cSP | CLIPC | CLIPC4 | AGAP000573 | ZOI | positive | 14.611 | 18.889 | 17.587 | 14.872 | 18.69 | 16.930 | 14.370 - 19.490 |
| cSP | CLIPD | CLIPD3 | AGAP001433 | ZOI | positive | 18.140 | 18.661 | 17.835 | 14.872 | 18.69 | 17.640 | 15.670 - 19.610 |

**Table S10: Zone of Inhibition of negative regulators of antimicrobial activity.** DsRNA injected mosquitoes were challenged with 50.6 nL of resuspended lyophilized *Micrococcus luteus* at $OD_{600}$= 0.1. *CACT* kd was used as a positive control for positive regulation of antimicrobial activity. Five independent biological replicates were performed using different mosquito generations. One-way ANOVA followed by Bonferroni's post-tests were performed on ZOI data to calculate statistical significance ($P < 0.05$).

| CLIP family | sub family | Gene name | AGAP | Assay type | Tested as | ds*GFP* controls (individual replicates) | | | | | | | ds*GOI* (individual replicates) | | | | | | | *P* |
|---|---|---|---|---|---|---|---|---|---|---|---|---|---|---|---|---|---|---|---|---|
| | | | | | | 1 | 2 | 3 | 4 | 5 | Mean | 95% CI of mean | 1 | 2 | 3 | 4 | 5 | Mean | 95% CI of mean | |
| cSPH | CLIPA | CLIPA4 | AGAP011780 | ZOI | negative | 8.665 | 11.315 | 11.921 | 10.284 | 9.546 | 10.350 | 8.717 - 11.980 | 15.046 | 19.054 | 19.054 | 19.823 | 16.719 | 17.940 | 15.460 - 20.410 | <0.0001 |
| cSPH | CLIPA | CLIPA19 | AGAP003245 | ZOI | negative | 9.068 | 10.085 | 11.167 | 10.284 | 9.546 | 10.030 | 9.044 - 11.020 | 14.088 | 15.284 | 13.955 | 14.975 | 16.374 | 14.940 | 13.710 - 16.160 | <0.0001 |

| CLIP family | sub family | Gene name | AGAP | Assay type | Tested as | Positive controls-ds*CACT* | | | | | | |
|---|---|---|---|---|---|---|---|---|---|---|---|---|
| | | | | | | 1 | 2 | 3 | 4 | 5 | Mean | 95% CI of mean |
| cSPH | CLIPA | CLIPA4 | AGAP011780 | ZOI | negative | 26.561 | 27.584 | 31.596 | 28.557 | 29.013 | 28.660 | 26.310 - 31.010 |
| cSPH | CLIPA | CLIPA19 | AGAP003245 | ZOI | negative | 27.274 | 29.486 | 30.015 | 28.557 | 29.013 | 28.870 | 27.570 - 30.160 |

**Table S11: Zone of Inhibition results after silencing Spätzle genes.** DsRNA injected mosquitoes were challenged with 50.6 nL of resuspended lyophilized *Micrococcus luteus* at $OD_{600} = 5$. *REL1* kd was used as a positive control for positive regulation of antimicrobial activity. Five independent biological replicates were performed using different mosquito generations. One-way ANOVA followed by Bonferroni's post-tests were performed on ZOI data to calculate statistical significance ($P < 0.05$).

| Gene name | AGAP | Assay type | Tested as | ds*GFP* controls (individual Replicates) | | | | | | | ds*GOI* (individual replicates) | | | | | | | *P* |
|---|---|---|---|---|---|---|---|---|---|---|---|---|---|---|---|---|---|---|
| | | | | 1 | 2 | 3 | 4 | 5 | Mean | 95% CI of mean | 1 | 2 | 3 | 4 | 5 | Mean | 95% CI of mean | |
| Spz1 | AGAP000346 | ZOI | positive | 33.382 | 36.831 | 34.995 | 33.194 | 34.537 | 34.590 | 32.772 - 36.414 | 12.125 | 12.206 | 10.787 | 9.265 | 12.917 | 11.460 | 9.661 - 13.265 | <0.0001 |
| Spz2 | AGAP029552 | ZOI | positive | | | | | | | | 32.684 | 30.628 | 34.258 | 32.164 | 33.843 | 32.720 | 30.932 - 34.516 | NS |
| Spz3 | AGAP008360 | ZOI | positive | | | | | | | | 12.554 | 11.277 | 13.070 | 14.862 | 13.103 | 12.970 | 11.374 - 14.570 | <0.0001 |
| Spz4 | AGAP007866 | ZOI | positive | | | | | | | | 34.933 | 31.878 | 35.852 | 32.086 | 32.397 | 27.430 | 21.681 - 33.184 | NS |
| Spz5 | AGAP007177 | ZOI | positive | | | | | | | | 32.711 | 35.635 | 36.642 | 34.549 | 35.491 | 35.010 | 33.175 - 36.859 | NS |
| Spz6 | AGAP005126 | ZOI | positive | | | | | | | | 34.736 | 33.722 | 33.378 | 32.508 | 33.163 | 33.500 | 32.483 - 34.520 | NS |

| Gene name | AGAP | Assay type | Tested as | Positive controls-ds*REL1* (individual replicates) | | | | | | |
|---|---|---|---|---|---|---|---|---|---|---|
| | | | | 1 | 2 | 3 | 4 | 5 | Mean | 95% CI of mean |
| Spz1 | AGAP000346 | ZOI | positive | 12.966 | 17.924 | 14.965 | 13.648 | 16.819 | 15.260 | 12.672-17.867 |
| Spz2 | AGAP029552 | ZOI | positive | | | | | | | |
| Spz3 | AGAP008360 | ZOI | positive | | | | | | | |
| Spz4 | AGAP007866 | ZOI | positive | | | | | | | |
| Spz5 | AGAP007177 | ZOI | positive | | | | | | | |
| Spz6 | AGAP005126 | ZOI | positive | | | | | | | |

**Table S12.** ***Spätzle*** **(*SPZ*) genes expression after challenge.** Expression data normalized as transcripts per million (TPM), and log2 TPM in adult naïve female mosquitoes (UC), *Micrococcus luteus* (Ml) and *Enterococcus faecalis* (Ef) challenged mosquitoes 12 h after bacterial injection. Fold change (FC) and log2 FC in gene expression after challenge relative to UC was corrected using the Bonferroni method as described in (*38*).

| Gene name | AGAP | ZOI phenotype | TPM | | | Log2TPM | | | Corrected FC | | Corrected Log2FC | |
|---|---|---|---|---|---|---|---|---|---|---|---|---|
| | | | UC_0h | Ml_12h | Ef_12h | UC_0h | Ml_12h | Ef_12h | Ml_12h | Ef_12h | Ml_12h | Ef_12h |
| SPZ1 | AGAP000346 | positive | 9.7 | 15.2 | 16.6 | 3.3 | 3.9 | 4.1 | 1.5 | 1.5 | 0.6 | 0.6 |
| SPZ2 | AGAP029552 | | 0.1 | 0.3 | 0.3 | -4.0 | -1.8 | -1.8 | 4.2 | 4.0 | 2.1 | 2.0 |
| SPZ3 | AGAP008360 | positive | 3.2 | 4.3 | 3.9 | 1.7 | 2.1 | 2.0 | 1.2 | 1.1 | 0.3 | 0.1 |
| SPZ4 | AGAP007866 | | 0.0 | 0.0 | 0.0 | N/A | N/A | N/A | N/A | N/A | N/A | N/A |
| SPZ5 | AGAP007177 | | 0.0 | 0.1 | 0.1 | N/A | -4.2 | -3.1 | 1.8 | 3.5 | 0.9 | 1.8 |
| SPZ6 | AGAP005126 | | 1.1 | 1.7 | 0.8 | 0.2 | 0.8 | -0.3 | 1.4 | -1.5 | 0.5 | -0.6 |

**Table S13: Bacterial load after silencing regulators of antimicrobial activity.** DsRNA injected mosquitoes were challenged with 50.6 nL of GFP-expressing, tetracycline-resistant *Enterococcus faecalis* suspension at $OD_{600}$ = 0.08. Candidate genes (8 pooled mosquito homogenates per replicate) were tested in two independent groups using their own positive (ds*REL1*) and negative (ds*CACT*) controls. Five independent biological replicates were performed using different mosquito generations. One-way ANOVA followed by Bonferroni's post-tests were performed on Colony-forming unit (CFU)/mosquito data to calculate statistical significance (P < 0.05).

| CLIP family | sub family | Gene name | AGAP | Assay type | ds*GFP* controls (individual Replicates) | | | | | | | ds*GOI* (individual replicates) | | | | | | | |
|---|---|---|---|---|---|---|---|---|---|---|---|---|---|---|---|---|---|---|---|
| | | | | | 1 | 2 | 3 | 4 | 5 | Mean | 95% CI of mean | 1 | 2 | 3 | 4 | 5 | Mean | 95% CI of mean | *P* |
| | | TEP1 | AGAP010815 | Bacteria proliferation | 913000 | 819000 | 919000 | 806000 | 744000 | 840200 | 747318 - 933082 | 3120000 | 3100000 | 3000000 | 2790000 | 2830000 | 2968000 | 2779406 - 3156594 | <0.0001 |
| | | MODSP1 | AGAP001798 | Bacteria proliferation | 913000 | 819000 | 919000 | 806000 | 744000 | 840200 | 747318 - 933082 | 2710000 | 2790000 | 3050000 | 3080000 | 2820000 | 2890000 | 2685031 - 3094969 | <0.0001 |
| cSPH | CLIPA | CLIPA3 | AGAP012591 | Bacteria proliferation | 913000 | 819000 | 919000 | 806000 | 744000 | 840200 | 747318 - 933082 | 1230000 | 1080000 | 1160000 | 1180000 | 1070000 | 1144000 | 1059512 - 1228488 | <0.01 |
| cSP | CLIPB | CLIPB10 | AGAP029770 | Bacteria proliferation | 913000 | 819000 | 919000 | 806000 | 744000 | 840200 | 747318 - 933082 | 2140000 | 2570000 | 2520000 | 2420000 | 2310000 | 2392000 | 2177763 - 2606237 | <0.0001 |
| cSP | CLIPB | CLIPB14 | AGAP010833 | Bacteria proliferation | 700000 | 790000 | 780000 | 880000 | 860000 | 802000 | 713154 - 890846 | 1700000 | 1400000 | 1500000 | 1500000 | 1400000 | 1500000 | 1347928 - 1652072 | <0.0001 |
| cSP | CLIPC | CLIPC2 | AGAP004317 | Bacteria proliferation | 913000 | 819000 | 919000 | 806000 | 744000 | 840200 | 747318 - 933082 | 1080000 | 1140000 | 1100000 | 1160000 | 1090000 | 1114000 | 1071347 - 1156653 | <0.01 |
| cSP | CLIPC | CLIPC4 | AGAP000573 | Bacteria proliferation | 700000 | 790000 | 780000 | 880000 | 860000 | 802000 | 713154 - 890846 | 970000 | 880000 | 850000 | 930000 | 950000 | 916000 | 854166 - 854166 | <0.0001 |
| cSP | CLIPD | CLIPD3 | AGAP001433 | Bacteria proliferation | 913000 | 819000 | 919000 | 806000 | 744000 | 840200 | 747318 - 933082 | 1240000 | 1080000 | 1160000 | 1180000 | 1070000 | 1146000 | 1057501 - 1234499 | <0.01 |
| cSPH | CLIPA | CLIPA4 | AGAP011780 | Bacteria proliferation | 700000 | 790000 | 780000 | 880000 | 860000 | 802000 | 713154 - 890846 | 69000 | 88000 | 100000 | 120000 | 110000 | 97400 | 72791 - 122009 | <0.0001 |
| cSPH | CLIPA | CLIPA19 | AGAP003245 | Bacteria proliferation | 913000 | 819000 | 919000 | 806000 | 744000 | 840200 | 747318 - 933082 | 531000 | 513000 | 563000 | 538000 | 575000 | 544000 | 513033 - 574967 | <0.01 |

| CLIP family | sub family | Gene name | AGAP | Assay type | Positive regulator control-ds*REL1* (individual replicates) | | | | | | | Negative regulator control-ds*CACT* (individual replicates) | | | | | | |
|---|---|---|---|---|---|---|---|---|---|---|---|---|---|---|---|---|---|---|
| | | | | | 1 | 2 | 3 | 4 | 5 | Mean | 95% CI of mean | 1 | 2 | 3 | 4 | 5 | Mean | 95% CI of mean |
| | | TEP1 | AGAP010815 | Bacteria proliferation | 3140000 | 2680000 | 2690000 | 2890000 | 2920000 | 2864000 | 2628313 - 3099687 | 181000 | 250000 | 194000 | 244000 | 169000 | 207600 | 161536 - 253664 |
| | | MODSP1 | AGAP001798 | Bacteria proliferation | 3140000 | 2680000 | 2690000 | 2890000 | 2920000 | 2864000 | 2628313 - 3099687 | 181000 | 250000 | 194000 | 244000 | 169000 | 207600 | 161536 - 253664 |
| cSPH | CLIPA | CLIPA3 | AGAP012591 | Bacteria proliferation | 3140000 | 2680000 | 2690000 | 2890000 | 2920000 | 2864000 | 2628313 - 3099687 | 181000 | 250000 | 194000 | 244000 | 169000 | 207600 | 161536 - 253664 |
| cSP | CLIPB | CLIPB10 | AGAP029770 | Bacteria proliferation | 3140000 | 2680000 | 2690000 | 2890000 | 2920000 | 2864000 | 2628313 - 3099687 | 181000 | 250000 | 194000 | 244000 | 169000 | 207600 | 161536 - 253664 |
| cSP | CLIPB | CLIPB14 | AGAP010833 | Bacteria proliferation | 2800000 | 3100000 | 2700000 | 2500000 | 2700000 | 2760000 | 2487965 - 3032035 | 140000 | 200000 | 150000 | 220000 | 130000 | 168000 | 118801 - 217199 |
| cSP | CLIPC | CLIPC2 | AGAP004317 | Bacteria proliferation | 3140000 | 2680000 | 2690000 | 2890000 | 2920000 | 2864000 | 2628313 - 3099687 | 181000 | 250000 | 194000 | 244000 | 169000 | 207600 | 161536 - 253664 |
| cSP | CLIPC | CLIPC4 | AGAP000573 | Bacteria proliferation | 2800000 | 3100000 | 2700000 | 2500000 | 2700000 | 2760000 | 2487965 - 3032035 | 140000 | 200000 | 150000 | 220000 | 130000 | 168000 | 118801 - 217199 |
| cSP | CLIPD | CLIPD3 | AGAP001433 | Bacteria proliferation | 3140000 | 2680000 | 2690000 | 2890000 | 2920000 | 2864000 | 2628313 - 3099687 | 181000 | 250000 | 194000 | 244000 | 169000 | 207600 | 161536 - 253664 |
| cSPH | CLIPA | CLIPA4 | AGAP011780 | Bacteria proliferation | 2800000 | 3100000 | 2700000 | 2500000 | 2700000 | 2760000 | 2487965 - 3032035 | 140000 | 200000 | 150000 | 220000 | 130000 | 168000 | 118801 - 217199 |
| cSPH | CLIPA | CLIPA19 | AGAP003245 | Bacteria proliferation | 3140000 | 2680000 | 2690000 | 2890000 | 2920000 | 2864000 | 2628313 - 3099687 | 181000 | 250000 | 194000 | 244000 | 169000 | 207600 | 161536 - 253664 |

**Table S14: Median Survival after silencing regulators of antimicrobial activity.** DsRNA injected mosquitoes were challenged with 50.6 nL of GFP-expressing, tetracycline-resistant *Enterococcus faecalis* suspension at $OD_{600}$ = 0.08. Candidate genes (35-40 mosquitoes per replicate) were tested in two independent groups using their own positive (ds*REL1*) and negative (ds*CACT*) controls. Five independent biological replicates were performed using different mosquito generations. One-way ANOVA followed by Bonferroni's post-tests were performed on Lethal time 50 ($LT_{50}$) data to calculate statistical significance ($P < 0.05$).

| CLIP family | sub family | Gene name | AGAP | Assay type | ds*GFP* controls (individual replicates) | | | | | | | ds*GOI* (individual replicates) | | | | | | | |
|---|---|---|---|---|---|---|---|---|---|---|---|---|---|---|---|---|---|---|---|
| | | | | | 1 | 2 | 3 | 4 | 5 | Mean | 95% CI of mean | 1 | 2 | 3 | 4 | 5 | Mean | 95% CI of mean | *P* |
| | | TEP1 | AGAP010815 | Survival | 17 | 17 | 13 | 15 | 18 | 15.8 | 13.37 - 18.23 | 2 | 3 | 2.5 | 2 | 2.5 | 2.4 | 1.88 - 2.92 | <0.0001 |
| | | MODSP1 | AGAP001798 | Survival | 17 | 17 | 13 | 15 | 18 | 15.8 | 13.37 - 18.23 | 11 | 9.5 | 8.5 | 9.5 | 10 | 9.7 | 8.57 - 10.83 | <0.0001 |
| cSPH | CLIPA | CLIPA3 | AGAP012591 | Survival | 17 | 17 | 13 | 15 | 18 | 15.8 | 13.37 - 18.23 | 16 | 17 | 15 | 16 | 18 | 16.3 | 14.68 - 17.92 | NS |
| cSP | CLIPB | CLIPB10 | AGAP029770 | Survival | 17 | 17 | 13 | 15 | 18 | 15.8 | 13.37 - 18.23 | 10 | 12 | 12 | 13 | 12 | 11.6 | 10.41 - 12.79 | <0.0001 |
| cSP | CLIPB | CLIPB14 | AGAP010833 | Survival | 17 | 16 | 16 | 17 | 19 | 16.8 | 15.51 - 18.09 | 14 | 10 | 12 | 13 | 14 | 12.5 | 10.54 - 14.46 | <0.0001 |
| cSP | CLIPC | CLIPC2 | AGAP004317 | Survival | 17 | 17 | 13 | 15 | 18 | 15.8 | 13.37 - 18.23 | 17 | 16 | 17 | 18 | 17 | 16.9 | 15.98 - 17.82 | NS |
| cSP | CLIPC | CLIPC4 | AGAP000573 | Survival | 17 | 16 | 16 | 17 | 19 | 16.8 | 15.51 - 18.09 | 14 | 15 | 14 | 15 | 17 | 15 | 13.48 - 16.52 | <0.0001 |
| cSP | CLIPD | CLIPD3 | AGAP001433 | Survival | 17 | 17 | 13 | 15 | 18 | 15.8 | 13.37 - 18.23 | 18 | 17 | 16 | 18 | 18 | 17.3 | 16.26 - 18.34 | NS |
| cSPH | CLIPA | CLIPA4 | AGAP011780 | Survival | 17 | 16 | 16 | 17 | 19 | 16.8 | 15.51 - 18.09 | 25 | 26 | 26 | 26 | 27 | 25.9 | 24.98 - 26.82 | <0.0001 |
| cSPH | CLIPA | CLIPA19 | AGAP003245 | Survival | 17 | 17 | 13 | 15 | 18 | 15.8 | 13.37 - 18.23 | 20 | 21 | 18 | 19 | 21 | 19.7 | 18.2 - 21.2 | <0.0001 |

| CLIP family | sub family | Gene name | AGAP | Assay type | Positive regulator control-ds*REL1* | | | | | | | Negative regulator control-ds*CACT* | | | | | | |
|---|---|---|---|---|---|---|---|---|---|---|---|---|---|---|---|---|---|---|
| | | | | | 1 | 2 | 3 | 4 | 5 | Mean | 95% CI of mean | 1 | 2 | 3 | 4 | 5 | Mean | 95% CI of mean |
| | | TEP1 | AGAP010815 | Survival | 6 | 4 | 4.5 | 6.5 | 5 | 5.2 | 3.913 - 6.487 | 21 | 20 | 20 | 21 | 21 | 20.4 | 19.88 - 20.92 |
| | | MODSP1 | AGAP001798 | Survival | 6 | 4 | 4.5 | 6.5 | 5 | 5.2 | 3.913 - 6.487 | 21 | 20 | 20 | 21 | 21 | 20.4 | 19.88 - 20.92 |
| cSPH | CLIPA | CLIPA3 | AGAP012591 | Survival | 6 | 4 | 4.5 | 6.5 | 5 | 5.2 | 3.913 - 6.487 | 21 | 20 | 20 | 21 | 21 | 20.4 | 19.88 - 20.92 |
| cSP | CLIPB | CLIPB10 | AGAP029770 | Survival | 6 | 4 | 4.5 | 6.5 | 5 | 5.2 | 3.913 - 6.487 | 21 | 20 | 20 | 21 | 21 | 20.4 | 19.88 - 20.92 |
| cSP | CLIPB | CLIPB14 | AGAP010833 | Survival | 4 | 4 | 5 | 4 | 3 | 4 | 3.122 - 4.878 | 21 | 19 | 19 | 21 | 21 | 20 | 18.84 - 21.16 |
| cSP | CLIPC | CLIPC2 | AGAP004317 | Survival | 6 | 4 | 4.5 | 6.5 | 5 | 5.2 | 3.913 - 6.487 | 21 | 20 | 20 | 21 | 21 | 20.4 | 19.88 - 20.92 |
| cSP | CLIPC | CLIPC4 | AGAP000573 | Survival | 4 | 4 | 5 | 4 | 3 | 4 | 3.122 - 4.878 | 21 | 19 | 19 | 21 | 21 | 20 | 18.84 - 21.16 |
| cSP | CLIPD | CLIPD3 | AGAP001433 | Survival | 6 | 4 | 4.5 | 6.5 | 5 | 5.2 | 3.913 - 6.487 | 21 | 20 | 20 | 21 | 21 | 20.4 | 19.88 - 20.92 |
| cSPH | CLIPA | CLIPA4 | AGAP011780 | Survival | 4 | 4 | 5 | 4 | 3 | 4 | 3.122 - 4.878 | 21 | 19 | 19 | 21 | 21 | 20 | 18.84 - 21.16 |
| cSPH | CLIPA | CLIPA19 | AGAP003245 | Survival | 6 | 4 | 4.5 | 6.5 | 5 | 5.2 | 3.913 - 6.487 | 21 | 20 | 20 | 21 | 21 | 20.4 | 19.88 - 20.92 |

**Table S15. *CLIP* genes expression after challenge.** Expression data normalized as transcripts per million (TPM), and log2 TPM in adult naïve female mosquitoes (UC), *Micrococcus luteus* (Ml) and *Enterococcus faecalis* (Ef) challenged mosquitoes 12 h after bacterial injection. Fold change (FC) and log2 FC in gene expression after challenge relative to UC was corrected using the Bonferroni method as described in (*38*).

| CLIP family | sub family | Gene name | AGAP | Melanization phenotype | ZOI phenotype | TPM | | | Log2TPM | | | Corrected FC | | Corrected Log2FC | |
|---|---|---|---|---|---|---|---|---|---|---|---|---|---|---|---|
| | | | | | | UC_0h | Ml_12h | Ef_12h | UC_0h | Ml_12h | Ef_12h | Ml_12h | Ef_12h | Ml_12h | Ef_12h |
| cSPH | CLIPA | CLIPA1 | AGAP011791 | | | 144.5 | 1241.2 | 1180.7 | 7.2 | 10.3 | 10.2 | 8.2 | 7.3 | 3.0 | 2.9 |
| cSPH | CLIPA | CLIPA2 | AGAP011790 | negative | | 35.7 | 490.9 | 486.4 | 5.2 | 8.9 | 8.9 | 13.3 | 12.2 | 3.7 | 3.6 |
| cSPH | CLIPA | CLIPA3 | AGAP012591 | | positive | 0.1 | 0.0 | 0.1 | -3.0 | N/A | -3.1 | -4.0 | -1.1 | -2.0 | -0.2 |
| cSPH | CLIPA | CLIPA4 | AGAP011780 | | negative | 42.5 | 151.5 | 164.7 | 5.4 | 7.2 | 7.4 | 3.4 | 3.5 | 1.8 | 1.8 |
| cSPH | CLIPA | CLIPA5 | AGAP011787 | negative | | 1.2 | 18.2 | 14.8 | 0.2 | 4.2 | 3.9 | 15.0 | 11.5 | 3.9 | 3.5 |
| cSPH | CLIPA | CLIPA6 | AGAP011789 | | | 483.0 | 1698.8 | 1775.7 | 8.9 | 10.7 | 10.8 | 3.3 | 3.3 | 1.7 | 1.7 |
| cSPH | CLIPA | CLIPA7 | AGAP011792 | negative | | 314.3 | 537.6 | 783.2 | 8.3 | 9.1 | 9.6 | 1.6 | 2.2 | 0.7 | 1.2 |
| cSPH | CLIPA | CLIPA8 | AGAP010731 | positive | | 118.6 | 557.6 | 456.3 | 6.9 | 9.1 | 8.8 | 4.5 | 3.5 | 2.2 | 1.8 |
| cSPH | CLIPA | CLIPA9 | AGAP010968 | | | 852.2 | 445.0 | 468.5 | 9.7 | 8.8 | 8.9 | -2.1 | -2.0 | -1.0 | -1.0 |
| cSPH | CLIPA | CLIPA10 | AGAP006954 | | | 0.1 | 0.0 | 0.0 | -4.0 | N/A | N/A | -2.1 | -2.1 | -1.1 | -1.1 |
| cSPH | CLIPA | CLIPA12 | AGAP011781 | | | 107.7 | 475.9 | 539.9 | 6.8 | 8.9 | 9.1 | 4.3 | 4.5 | 2.1 | 2.2 |
| cSPH | CLIPA | CLIPA13 | AGAP011783 | negative | | 0.4 | 0.5 | 0.2 | -1.4 | -1.1 | -2.0 | 1.1 | -1.7 | 0.2 | -0.8 |
| cSPH | CLIPA | CLIPA14 | AGAP011788 | negative | | 245.5 | 1124.5 | 1015.0 | 7.9 | 10.1 | 10.0 | 4.4 | 3.7 | 2.1 | 1.9 |
| cSPH | CLIPA | CLIPA15 | AGAP002815 | | | 0.0 | 0.0 | 0.1 | N/A | N/A | -4.1 | N/A | 1.8 | N/A | 0.8 |
| cSPH | CLIPA | CLIPA19 | AGAP003245 | | negative | 17.8 | 20.3 | 25.2 | 4.2 | 4.3 | 4.7 | 1.1 | 1.3 | 0.1 | 0.4 |
| cSPH | CLIPA | CLIPA26 | AGAP001964 | | | 9.0 | 4.9 | 8.4 | 3.2 | 2.3 | 3.1 | -2.0 | -1.2 | -1.0 | -0.3 |
| cSPH | CLIPA | CLIPA27 | AGAP000290 | | | 15.0 | 60.4 | 53.7 | 3.9 | 5.9 | 5.7 | 3.8 | 3.2 | 1.9 | 1.7 |
| cSPH | CLIPA | CLIPA28 | AGAP010730 | positive | | 67.1 | 313.2 | 310.4 | 6.1 | 8.3 | 8.3 | 4.5 | 4.2 | 2.2 | 2.1 |
| cSPH | CLIPA | SPCLIP1 | AGAP028725 | positive | | 70.8 | 504.3 | 474.6 | 6.1 | 9.0 | 8.9 | 6.8 | 6.0 | 2.8 | 2.6 |
| cSP | CLIPB | CLIPB1 | AGAP003251 | | | 252.0 | 276.7 | 450.1 | 8.0 | 8.1 | 8.8 | 1.0 | 1.6 | 0.1 | 0.7 |
| cSP | CLIPB | CLIPB2 | AGAP003246 | | | 51.6 | 158.2 | 257.5 | 5.7 | 7.3 | 8.0 | 3.0 | 4.5 | 1.6 | 2.2 |
| cSP | CLIPB | CLIPB3b | AGAP003249 | | | 0.2 | 0.7 | 2.0 | -2.1 | -0.5 | 1.0 | 2.8 | 7.8 | 1.5 | 3.0 |
| cSP | CLIPB | CLIPB4 | AGAP003250 | positive | | 415.5 | 585.9 | 1007.2 | 8.7 | 9.2 | 10.0 | 1.3 | 2.2 | 0.4 | 1.1 |
| cSP | CLIPB | CLIPB5 | AGAP004148 | | | 35.2 | 46.8 | 58.5 | 5.1 | 5.5 | 5.9 | 1.3 | 1.5 | 0.3 | 0.6 |
| cSP | CLIPB | CLIPB6 | AGAP003252 | | | 9.5 | 11.4 | 10.4 | 3.2 | 3.5 | 3.4 | 1.1 | -1.0 | 0.2 | 0.0 |

| | | | | | | | | | | | | | | | |
|---|---|---|---|---|---|---|---|---|---|---|---|---|---|---|---|
| cSPH | CLIPB | CLIPB7 | AGAP002270 | | | 20.8 | 20.1 | 28.4 | 4.4 | 4.3 | 4.8 | -1.1 | 1.2 | -0.1 | 0.3 |
| cSP | CLIPB | CLIPB8 | AGAP003057 | positive | | 73.1 | 166.2 | 246.0 | 6.2 | 7.4 | 7.9 | 2.2 | 3.0 | 1.1 | 1.6 |
| cSP | CLIPB | CLIPB9 | AGAP029769 | positive | | 2.0 | 2.0 | 2.7 | 1.0 | 1.0 | 1.4 | -1.1 | 1.2 | -0.1 | 0.2 |
| cSP | CLIPB | CLIPB10 | AGAP029770 | positive | positive | 111.4 | 68.9 | 168.0 | 6.8 | 6.1 | 7.4 | -1.7 | 1.4 | -0.8 | 0.4 |
| cSP | CLIPB | CLIPB11 | AGAP009214 | | | 1.2 | 11.6 | 14.4 | 0.3 | 3.5 | 3.8 | 9.7 | 11.1 | 3.3 | 3.5 |
| cSPH | CLIPB | CLIPB12 | AGAP009217 | negative | | 57.9 | 307.1 | 257.7 | 5.9 | 8.3 | 8.0 | 4.8 | 3.9 | 2.3 | 2.0 |
| cSP | CLIPB | CLIPB13 | AGAP004855 | positive | | 105.4 | 154.8 | 187.8 | 6.7 | 7.3 | 7.6 | 1.4 | 1.6 | 0.5 | 0.7 |
| cSP | CLIPB | CLIPB14 | AGAP010833 | | positive | 14.4 | 57.3 | 50.1 | 3.8 | 5.8 | 5.6 | 3.8 | 3.1 | 1.9 | 1.7 |
| cSP | CLIPB | CLIPB15 | AGAP009844 | | | 58.3 | 697.9 | 457.7 | 5.9 | 9.4 | 8.8 | 11.6 | 7.1 | 3.5 | 2.8 |
| cSPH | CLIPB | CLIPB16 | AGAP009263 | | | 2.1 | 1.7 | 2.1 | 1.0 | 0.8 | 1.0 | -1.3 | -1.1 | -0.3 | -0.1 |
| cSP | CLIPB | CLIPB17 | AGAP001648 | positive | | 4.7 | 220.8 | 138.6 | 2.2 | 7.8 | 7.1 | 46.6 | 26.6 | 5.5 | 4.7 |
| cSP | CLIPB | CLIPB18 | AGAP009215 | | | 4.5 | 49.6 | 32.6 | 2.2 | 5.6 | 5.0 | 10.7 | 6.4 | 3.4 | 2.7 |
| cSP | CLIPB | CLIPB19 | AGAP003247 | | | 9.2 | 71.1 | 118.5 | 3.2 | 6.2 | 6.9 | 7.5 | 11.6 | 2.9 | 3.5 |
| cSP | CLIPB | CLIPB20 | AGAP012037 | positive | | 2.0 | 28.5 | 26.8 | 1.0 | 4.8 | 4.7 | 13.6 | 11.9 | 3.8 | 3.6 |
| cSPH | CLIPB | CLIPB36 | AGAP013184 | | | 15.8 | 84.1 | 82.5 | 4.0 | 6.4 | 6.4 | 5.2 | 4.7 | 2.4 | 2.2 |
| cSP | CLIPB | CLIPB41 | AGAP004149 | | | 0.0 | 0.0 | 0.0 | N/A | N/A | N/A | N/A | N/A | N/A | N/A |
| cSP | CLIPB | CLIPB44 | AGAP009220 | | | 1.4 | 4.7 | 3.9 | 0.5 | 2.2 | 1.9 | 3.1 | 2.4 | 1.6 | 1.2 |
| cSP | CLIPB | CLIPB46 | AGAP009849 | | | 3.5 | 69.7 | 41.0 | 1.8 | 6.1 | 5.4 | 18.8 | 10.3 | 4.2 | 3.4 |
| cSP | CLIPB | CLIPB47 | AGAP003686 | | | 9.7 | 10.3 | 7.3 | 3.3 | 3.4 | 2.9 | 1.0 | -1.5 | 0.0 | -0.6 |
| cSP | CLIPC | CLIPC1 | AGAP008835 | | | 0.0 | 0.0 | 0.0 | N/A | N/A | N/A | N/A | N/A | N/A | N/A |
| cSP | CLIPC | CLIPC2 | AGAP004317 | | positive | 26.0 | 51.2 | 60.1 | 4.7 | 5.7 | 5.9 | 1.8 | 2.0 | 0.9 | 1.0 |
| cSP | CLIPC | CLIPC3 | AGAP004318 | positive | | 80.7 | 67.1 | 85.5 | 6.3 | 6.1 | 6.4 | -1.3 | -1.1 | -0.4 | -0.1 |
| cSP | CLIPC | CLIPC4 | AGAP000573 | | positive | 78.7 | 109.9 | 103.6 | 6.3 | 6.8 | 6.7 | 1.3 | 1.2 | 0.4 | 0.2 |
| cSP | CLIPC | CLIPC5 | AGAP000571 | | | 1.9 | 1.9 | 1.6 | 0.9 | 0.9 | 0.7 | -1.0 | -1.3 | -0.1 | -0.4 |
| cSP | CLIPC | CLIPC6 | AGAP000315 | | | 65.3 | 59.7 | 69.4 | 6.0 | 5.9 | 6.1 | -1.2 | -1.1 | -0.2 | -0.1 |
| cSP | CLIPC | CLIPC7 | AGAP003689 | | | 41.0 | 1303.6 | 938.3 | 5.4 | 10.3 | 9.9 | 30.6 | 20.6 | 4.9 | 4.4 |
| cSP | CLIPC | CLIPC9 | AGAP004719 | positive | | 17.8 | 19.5 | 20.4 | 4.2 | 4.3 | 4.4 | 1.0 | 1.0 | 0.0 | 0.0 |
| cSP | CLIPC | CLIPC10 | AGAP000572 | | | 12.8 | 13.0 | 16.2 | 3.7 | 3.7 | 4.0 | -1.0 | 1.1 | -0.1 | 0.2 |
| cSP | CLIPC | CLIPC12 | AGAP012034 | positive | | 14.5 | 12.9 | 11.4 | 3.9 | 3.7 | 3.5 | -1.2 | -1.4 | -0.3 | -0.5 |
| cSP | CLIPC | CLIPC14 | AGAP028167 | positive | | 0.8 | 5.3 | 3.8 | -0.3 | 2.4 | 1.9 | 6.2 | 4.1 | 2.6 | 2.0 |
| cSP | CLIPD | CLIPD1 | AGAP002422 | | | 27.7 | 11.4 | 16.8 | 4.8 | 3.5 | 4.1 | -2.6 | -1.8 | -1.4 | -0.9 |
| cSP | CLIPD | CLIPD2 | AGAP008183 | | | 0.2 | 0.1 | 0.5 | -2.1 | -3.1 | -1.1 | -2.0 | 1.9 | -1.0 | 0.9 |
| cSP | CLIPD | CLIPD3 | AGAP001433 | | positive | 1.3 | 1.7 | 1.1 | 0.4 | 0.7 | 0.1 | 1.2 | -1.4 | 0.2 | -0.5 |

| | | | | | | | | | | | | | | | |
|---|---|---|---|---|---|---|---|---|---|---|---|---|---|---|---|
| cSP | CLIPD | CLIPD4 | AGAP002811 | | | 0.6 | 0.7 | 0.6 | -0.8 | -0.5 | -0.7 | 1.2 | -1.1 | 0.2 | -0.1 |
| cSP | CLIPD | CLIPD6 | AGAP002813 | | | 24.2 | 32.2 | 35.1 | 4.6 | 5.0 | 5.1 | 1.3 | 1.3 | 0.3 | 0.4 |
| cSP | CLIPD | CLIPD7 | AGAP008998 | | | 0.0 | 0.0 | 0.0 | N/A | N/A | N/A | N/A | N/A | N/A | N/A |
| cSP | CLIPD | CLIPD8 | AGAP002784 | | | 0.1 | 0.5 | 0.2 | -3.1 | -0.9 | -2.1 | 4.2 | 1.9 | 2.1 | 0.9 |
| cSP | CLIPD | CLIPD9 | AGAP008997 | | | 0.0 | 0.0 | 0.0 | N/A | N/A | N/A | N/A | N/A | N/A | N/A |
| cSP | CLIPD | CLIPD11 | AGAP029106 | | | 7.6 | 11.1 | 8.3 | 2.9 | 3.5 | 3.1 | 1.3 | -1.0 | 0.4 | 0.0 |
| cSP | CLIPD | CLIPD12 | AGAP008995 | | | 0.0 | 0.0 | 0.0 | N/A | N/A | N/A | N/A | N/A | N/A | N/A |
| cSP | CLIPD | CLIPD13 | AGAP009000 | | | 0.0 | 0.0 | 0.0 | N/A | N/A | N/A | N/A | N/A | N/A | N/A |
| cSP | CLIPD | CLIPD14 | AGAP009006 | | | 5.7 | 6.2 | 6.3 | 2.5 | 2.6 | 2.7 | 1.1 | 1.0 | 0.1 | 0.0 |
| cSP | CLIPD | CLIPD20 | AGAP013089 | | | 0.1 | 0.2 | 0.1 | -4.0 | -2.1 | -4.0 | 3.5 | -1.1 | 1.8 | -0.1 |
| cSP | CLIPD | CLIPD22 | AGAP008996 | | | 0.0 | 0.0 | 0.0 | N/A | N/A | N/A | N/A | N/A | N/A | N/A |
| cSPH | CLIPE | CLIPE1 | AGAP008091 | | | 2.6 | 2.7 | 2.3 | 1.4 | 1.4 | 1.2 | -1.1 | -1.3 | -0.1 | -0.4 |
| cSPH | CLIPE | CLIPE4 | AGAP010530 | | | 98.7 | 196.3 | 226.9 | 6.6 | 7.6 | 7.8 | 1.9 | 2.1 | 0.9 | 1.0 |
| cSP | CLIPE | CLIPE5 | AGAP028728 | | | 16.9 | 42.7 | 55.4 | 4.1 | 5.4 | 5.8 | 2.4 | 2.9 | 1.3 | 1.5 |
| cSPH | CLIPE | CLIPE6 | AGAP011785 | | | 6.4 | 7.2 | 7.3 | 2.7 | 2.9 | 2.9 | 1.0 | 1.0 | 0.0 | 0.0 |
| cSPH | CLIPE | CLIPE7 | AGAP011786 | | | 1.5 | 1.7 | 3.6 | 0.6 | 0.7 | 1.8 | 1.0 | 2.1 | 0.0 | 1.1 |
| cSP | CLIPE | CLIPE10 | AGAP010545 | | | 19.2 | 62.1 | 55.1 | 4.3 | 6.0 | 5.8 | 3.1 | 2.6 | 1.6 | 1.4 |
| cSPH | CLIPE | CLIPE12 | AGAP003691 | | | 7.8 | 21.9 | 68.9 | 3.0 | 4.5 | 6.1 | 2.6 | 7.9 | 1.4 | 3.0 |
| cSP | CLIPE | CLIPE13 | AGAP003748 | | | 0.3 | 0.5 | 0.8 | -2.0 | -1.1 | -0.3 | 1.6 | 2.6 | 0.7 | 1.4 |
| cSP | CLIPE | CLIPE15 | AGAP009249 | | | 0.0 | 0.0 | 0.0 | N/A | N/A | N/A | N/A | N/A | N/A | N/A |
| cSPH | CLIPE | CLIPE17 | AGAP009252 | | | 8.6 | 12.2 | 11.1 | 3.1 | 3.6 | 3.5 | 1.3 | 1.2 | 0.4 | 0.2 |
| cSP | CLIPE | CLIPE18 | AGAP009273 | | | 0.2 | 24.0 | 3.0 | -2.0 | 4.6 | 1.6 | 99.0 | 11.6 | 6.6 | 3.5 |
| cSP | CLIPE | CLIPE19 | AGAP011040 | | | 0.0 | 0.0 | 0.0 | N/A | N/A | N/A | N/A | N/A | N/A | N/A |
| cSP | CLIPE | CLIPE21 | AGAP011719 | | | 0.0 | 0.0 | 0.0 | N/A | N/A | N/A | N/A | N/A | N/A | N/A |
| cSP | CLIPE | CLIPE22 | AGAP012020 | | | 0.1 | 0.2 | 0.2 | -4.0 | -2.5 | -2.5 | 2.6 | 2.6 | 1.4 | 1.4 |
| cSP | CLIPE | CLIPE25 | AGAP012502 | | | 0.0 | 0.0 | 0.0 | N/A | N/A | N/A | N/A | N/A | N/A | N/A |
| cSP | CLIPE | CLIPE30 | AGAP010628 | | | 0.0 | 0.0 | 0.0 | N/A | N/A | N/A | N/A | N/A | N/A | N/A |

**Table S16: Edge list of AgMelGCN2.0.** Edge list is represented in a three-column matrix consisting of (i) the AGAP number of start vertex (node1), (ii) AGAP number of the end vertex (node2), and (iii) the edge weight. Please note that the network is undirected.

| Undirected Edge | | Edge weight |
|---|---|---|
| Node 1 | Node 2 | |
| AGAP007036 | AGAP007035 | 0.497 |
| AGAP007035 | AGAP011787 | 0.445 |
| AGAP007035 | AGAP029074 | 0.450 |
| AGAP007033 | AGAP011791 | 0.547 |
| AGAP007033 | AGAP011790 | 0.640 |
| AGAP007033 | AGAP000290 | 0.627 |
| AGAP007033 | AGAP010730 | 0.591 |
| AGAP007033 | AGAP011787 | 0.430 |
| AGAP007033 | AGAP011792 | 0.440 |
| AGAP007033 | AGAP010731 | 0.558 |
| AGAP007033 | AGAP010968 | 0.570 |
| AGAP007033 | AGAP029770 | 0.454 |
| AGAP007033 | AGAP009217 | 0.463 |
| AGAP007033 | AGAP004855 | 0.481 |
| AGAP007033 | AGAP009844 | 0.571 |
| AGAP007033 | AGAP001648 | 0.429 |
| AGAP007033 | AGAP003246 | 0.519 |
| AGAP007033 | AGAP012037 | 0.518 |
| AGAP007033 | AGAP009849 | 0.494 |
| AGAP007033 | AGAP012034 | 0.437 |
| AGAP007033 | AGAP003689 | 0.511 |
| AGAP007033 | AGAP004719 | 0.456 |
| AGAP007033 | AGAP010545 | 0.461 |
| AGAP007033 | AGAP010530 | 0.511 |
| AGAP007033 | AGAP028728 | 0.498 |
| AGAP007033 | AGAP005335 | 0.570 |
| AGAP007033 | AGAP005334 | 0.529 |
| AGAP007033 | AGAP006348 | 0.546 |
| AGAP007033 | AGAP007037 | 0.450 |
| AGAP007033 | AGAP007039 | 0.448 |
| AGAP007033 | AGAP004977 | 0.460 |
| AGAP007033 | AGAP005625 | 0.508 |
| AGAP007033 | AGAP006909 | 0.440 |
| AGAP007033 | AGAP001377 | 0.466 |
| AGAP007033 | AGAP006910 | 0.567 |
| AGAP007033 | AGAP008654 | 0.576 |
| AGAP007033 | AGAP008368 | 0.504 |
| AGAP007033 | AGAP008364 | 0.446 |
| AGAP007033 | AGAP010816 | 0.571 |
| AGAP007033 | AGAP010812 | 0.456 |
| AGAP011791 | AGAP011788 | 0.475 |
| AGAP011791 | AGAP011790 | 0.570 |
| AGAP011791 | AGAP000290 | 0.488 |
| AGAP011791 | AGAP010730 | 0.637 |
| AGAP011791 | AGAP011780 | 0.459 |
| AGAP011791 | AGAP011789 | 0.467 |
| AGAP011791 | AGAP011792 | 0.507 |
| AGAP011791 | AGAP010731 | 0.571 |
| AGAP011791 | AGAP010968 | 0.635 |
| AGAP011791 | AGAP003251 | 0.434 |
| AGAP011791 | AGAP029770 | 0.479 |
| AGAP011791 | AGAP009217 | 0.506 |
| AGAP011791 | AGAP004855 | 0.512 |
| AGAP011791 | AGAP009844 | 0.594 |
| AGAP011791 | AGAP003246 | 0.489 |
| AGAP011791 | AGAP012037 | 0.452 |
| AGAP011791 | AGAP003250 | 0.478 |
| AGAP011791 | AGAP003057 | 0.469 |
| AGAP011791 | AGAP012034 | 0.569 |
| AGAP011791 | AGAP004318 | 0.463 |
| AGAP011791 | AGAP003689 | 0.459 |
| AGAP011791 | AGAP004719 | 0.463 |
| AGAP011791 | AGAP002813 | 0.490 |
| AGAP011791 | AGAP010545 | 0.439 |
| AGAP011791 | AGAP010530 | 0.452 |
| AGAP011791 | AGAP028728 | 0.511 |
| AGAP011791 | AGAP005335 | 0.581 |
| AGAP011791 | AGAP028725 | 0.609 |
| AGAP011791 | AGAP001377 | 0.503 |
| AGAP011791 | AGAP008654 | 0.497 |
| AGAP011791 | AGAP010816 | 0.481 |
| AGAP006954 | AGAP002815 | 0.517 |
| AGAP006954 | AGAP008995 | 0.619 |
| AGAP006954 | AGAP009000 | 0.552 |
| AGAP006954 | AGAP008183 | 0.447 |
| AGAP006954 | AGAP008996 | 0.554 |
| AGAP006954 | AGAP001433 | 0.535 |
| AGAP006954 | AGAP008998 | 0.583 |
| AGAP006954 | AGAP002784 | 0.450 |
| AGAP006954 | AGAP012021 | 0.466 |
| AGAP006954 | AGAP012502 | 0.574 |
| AGAP006954 | AGAP029559 | 0.594 |
| AGAP006954 | AGAP007408 | 0.518 |
| AGAP006954 | AGAP002825 | 0.513 |
| AGAP006954 | AGAP004980 | 0.539 |
| AGAP011781 | AGAP011788 | 0.481 |
| AGAP011781 | AGAP011780 | 0.504 |
| AGAP011781 | AGAP003251 | 0.451 |
| AGAP011781 | AGAP003250 | 0.462 |
| AGAP011781 | AGAP029100 | 0.456 |
| AGAP011783 | AGAP011780 | 0.463 |
| AGAP011783 | AGAP028007 | 0.544 |
| AGAP011783 | AGAP027997 | 0.468 |
| AGAP011788 | AGAP010730 | 0.449 |
| AGAP011788 | AGAP011780 | 0.510 |
| AGAP011788 | AGAP011789 | 0.639 |
| AGAP011788 | AGAP011792 | 0.536 |
| AGAP011788 | AGAP003251 | 0.435 |
| AGAP011788 | AGAP004855 | 0.532 |
| AGAP011788 | AGAP003250 | 0.497 |
| AGAP011788 | AGAP003057 | 0.443 |
| AGAP011788 | AGAP005334 | 0.439 |
| AGAP011788 | AGAP006348 | 0.469 |
| AGAP011788 | AGAP005625 | 0.497 |
| AGAP011788 | AGAP028725 | 0.552 |
| AGAP011788 | AGAP001377 | 0.521 |
| AGAP011788 | AGAP006911 | 0.519 |
| AGAP002815 | AGAP008995 | 0.603 |
| AGAP002815 | AGAP009000 | 0.500 |
| AGAP002815 | AGAP008996 | 0.549 |
| AGAP002815 | AGAP008998 | 0.430 |
| AGAP002815 | AGAP011782 | 0.458 |
| AGAP002815 | AGAP012021 | 0.541 |
| AGAP002815 | AGAP012502 | 0.537 |
| AGAP002815 | AGAP000123 | 0.453 |
| AGAP002815 | AGAP002825 | 0.523 |
| AGAP002815 | AGAP004980 | 0.493 |
| AGAP003245 | AGAP001964 | 0.447 |
| AGAP003245 | AGAP000290 | 0.442 |
| AGAP003245 | AGAP009263 | 0.475 |
| AGAP003245 | AGAP003246 | 0.472 |
| AGAP003245 | AGAP004719 | 0.469 |
| AGAP003245 | AGAP010545 | 0.483 |
| AGAP003245 | AGAP010530 | 0.499 |
| AGAP003245 | AGAP007045 | 0.441 |
| AGAP003245 | AGAP006258 | 0.445 |
| AGAP003245 | AGAP008654 | 0.479 |
| AGAP003245 | AGAP008368 | 0.454 |
| AGAP003245 | AGAP010832 | 0.492 |
| AGAP003245 | AGAP010816 | 0.450 |
| AGAP011790 | AGAP000290 | 0.441 |
| AGAP011790 | AGAP010730 | 0.655 |
| AGAP011790 | AGAP010731 | 0.598 |
| AGAP011790 | AGAP010968 | 0.519 |
| AGAP011790 | AGAP003250 | 0.475 |
| AGAP011790 | AGAP012034 | 0.466 |
| AGAP011790 | AGAP004719 | 0.477 |
| AGAP011790 | AGAP010545 | 0.524 |
| AGAP011790 | AGAP028728 | 0.448 |
| AGAP011790 | AGAP005335 | 0.513 |
| AGAP011790 | AGAP005334 | 0.439 |
| AGAP011790 | AGAP006348 | 0.537 |
| AGAP011790 | AGAP007039 | 0.450 |
| AGAP011790 | AGAP027997 | 0.492 |
| AGAP011790 | AGAP004977 | 0.433 |
| AGAP011790 | AGAP028725 | 0.472 |
| AGAP011790 | AGAP006910 | 0.526 |
| AGAP011790 | AGAP008654 | 0.461 |
| AGAP011790 | AGAP008368 | 0.477 |
| AGAP011790 | AGAP008364 | 0.444 |
| AGAP001964 | AGAP000290 | 0.467 |
| AGAP001964 | AGAP011787 | 0.476 |
| AGAP001964 | AGAP012504 | 0.470 |
| AGAP001964 | AGAP027997 | 0.476 |
| AGAP001964 | AGAP008654 | 0.473 |
| AGAP001964 | AGAP008368 | 0.504 |
| AGAP000290 | AGAP010730 | 0.535 |
| AGAP000290 | AGAP011787 | 0.449 |
| AGAP000290 | AGAP011792 | 0.565 |
| AGAP000290 | AGAP010731 | 0.605 |
| AGAP000290 | AGAP010968 | 0.542 |
| AGAP000290 | AGAP009844 | 0.503 |
| AGAP000290 | AGAP001648 | 0.503 |
| AGAP000290 | AGAP003246 | 0.567 |
| AGAP000290 | AGAP009849 | 0.489 |
| AGAP000290 | AGAP003057 | 0.516 |
| AGAP000290 | AGAP012034 | 0.437 |
| AGAP000290 | AGAP004719 | 0.525 |
| AGAP000290 | AGAP010545 | 0.515 |
| AGAP000290 | AGAP028728 | 0.434 |
| AGAP000290 | AGAP005335 | 0.482 |
| AGAP000290 | AGAP005334 | 0.516 |
| AGAP000290 | AGAP006348 | 0.468 |
| AGAP000290 | AGAP007045 | 0.471 |
| AGAP000290 | AGAP007037 | 0.484 |
| AGAP000290 | AGAP007454 | 0.439 |
| AGAP000290 | AGAP004977 | 0.526 |
| AGAP000290 | AGAP006910 | 0.510 |
| AGAP000290 | AGAP008654 | 0.513 |
| AGAP000290 | AGAP008368 | 0.459 |
| AGAP000290 | AGAP010816 | 0.475 |
| AGAP010730 | AGAP011780 | 0.437 |
| AGAP010730 | AGAP011792 | 0.437 |
| AGAP010730 | AGAP010731 | 0.606 |
| AGAP010730 | AGAP010968 | 0.616 |
| AGAP010730 | AGAP029770 | 0.558 |
| AGAP010730 | AGAP009217 | 0.450 |
| AGAP010730 | AGAP004855 | 0.596 |
| AGAP010730 | AGAP009844 | 0.643 |
| AGAP010730 | AGAP001648 | 0.429 |
| AGAP010730 | AGAP003246 | 0.503 |
| AGAP010730 | AGAP012037 | 0.500 |
| AGAP010730 | AGAP003250 | 0.450 |
| AGAP010730 | AGAP002270 | 0.436 |
| AGAP010730 | AGAP012034 | 0.576 |
| AGAP010730 | AGAP003689 | 0.472 |
| AGAP010730 | AGAP004719 | 0.466 |
| AGAP010730 | AGAP002813 | 0.490 |
| AGAP010730 | AGAP010530 | 0.494 |
| AGAP010730 | AGAP028728 | 0.549 |
| AGAP010730 | AGAP011785 | 0.440 |
| AGAP010730 | AGAP027997 | 0.454 |
| AGAP010730 | AGAP012616 | 0.488 |
| AGAP010730 | AGAP004977 | 0.499 |
| AGAP010730 | AGAP005625 | 0.531 |
| AGAP010730 | AGAP028725 | 0.570 |
| AGAP010730 | AGAP001377 | 0.521 |
| AGAP010730 | AGAP006911 | 0.441 |
| AGAP010730 | AGAP006910 | 0.535 |
| AGAP010730 | AGAP010815 | 0.483 |
| AGAP010730 | AGAP008654 | 0.495 |
| AGAP010730 | AGAP008364 | 0.486 |
| AGAP010730 | AGAP010816 | 0.544 |
| AGAP012591 | AGAP011782 | 0.536 |
| AGAP012591 | AGAP012021 | 0.437 |
| AGAP012591 | AGAP029074 | 0.468 |
| AGAP012591 | AGAP004980 | 0.469 |
| AGAP011793 | AGAP011794 | 0.456 |
| AGAP011793 | AGAP009273 | 0.489 |
| AGAP011793 | AGAP011040 | 0.555 |
| AGAP011793 | AGAP011719 | 0.590 |
| AGAP011793 | AGAP012502 | 0.536 |
| AGAP011793 | AGAP009316 | 0.469 |
| AGAP011793 | AGAP000940 | 0.476 |
| AGAP011793 | AGAP010708 | 0.506 |
| AGAP011793 | AGAP010830 | 0.436 |
| AGAP011794 | AGAP009000 | 0.433 |
| AGAP011794 | AGAP013089 | 0.549 |
| AGAP011794 | AGAP012502 | 0.441 |
| AGAP011794 | AGAP000940 | 0.433 |
| AGAP011794 | AGAP007408 | 0.574 |
| AGAP011794 | AGAP004980 | 0.443 |
| AGAP011780 | AGAP011792 | 0.530 |
| AGAP011780 | AGAP010968 | 0.430 |
| AGAP011780 | AGAP003251 | 0.437 |
| AGAP011780 | AGAP029770 | 0.485 |
| AGAP011780 | AGAP009214 | 0.563 |
| AGAP011780 | AGAP004855 | 0.484 |
| AGAP011780 | AGAP003250 | 0.539 |
| AGAP011780 | AGAP004148 | 0.513 |
| AGAP011780 | AGAP003057 | 0.508 |
| AGAP011780 | AGAP029769 | 0.520 |
| AGAP011780 | AGAP000572 | 0.467 |
| AGAP011780 | AGAP003689 | 0.450 |
| AGAP011780 | AGAP029074 | 0.429 |
| AGAP011780 | AGAP005625 | 0.513 |
| AGAP011780 | AGAP028725 | 0.447 |
| AGAP011780 | AGAP001377 | 0.518 |
| AGAP011780 | AGAP006911 | 0.462 |
| AGAP011787 | AGAP009217 | 0.515 |
| AGAP011787 | AGAP009849 | 0.440 |
| AGAP011787 | AGAP008654 | 0.511 |
| AGAP011787 | AGAP008368 | 0.434 |
| AGAP011787 | AGAP010816 | 0.459 |
| AGAP011787 | AGAP010812 | 0.453 |
| AGAP011789 | AGAP011792 | 0.472 |
| AGAP011789 | AGAP003251 | 0.472 |
| AGAP011789 | AGAP004855 | 0.489 |
| AGAP011789 | AGAP003250 | 0.430 |
| AGAP011789 | AGAP002813 | 0.443 |
| AGAP011789 | AGAP006348 | 0.456 |
| AGAP011789 | AGAP028725 | 0.460 |
| AGAP011789 | AGAP001377 | 0.497 |
| AGAP011789 | AGAP006911 | 0.521 |
| AGAP011792 | AGAP010731 | 0.472 |
| AGAP011792 | AGAP010968 | 0.516 |
| AGAP011792 | AGAP003251 | 0.473 |
| AGAP011792 | AGAP009844 | 0.450 |

| | | |
|---|---|---|
| AGAP011791 | AGAP005334 | 0.621 |
| AGAP011791 | AGAP006348 | 0.645 |
| AGAP011791 | AGAP004977 | 0.480 |
| AGAP011791 | AGAP005625 | 0.512 |
| AGAP011792 | AGAP002270 | 0.474 |
| AGAP011792 | AGAP003057 | 0.593 |
| AGAP011792 | AGAP000572 | 0.473 |
| AGAP011792 | AGAP004318 | 0.490 |
| AGAP011792 | AGAP004719 | 0.512 |
| AGAP011792 | AGAP009252 | 0.435 |
| AGAP011792 | AGAP005335 | 0.446 |
| AGAP011792 | AGAP005334 | 0.507 |
| AGAP011792 | AGAP006348 | 0.571 |
| AGAP011792 | AGAP007039 | 0.439 |
| AGAP011792 | AGAP004977 | 0.484 |
| AGAP011792 | AGAP005625 | 0.521 |
| AGAP011792 | AGAP028725 | 0.535 |
| AGAP011792 | AGAP001377 | 0.554 |
| AGAP011792 | AGAP001376 | 0.437 |
| AGAP011792 | AGAP006911 | 0.497 |
| AGAP011792 | AGAP006910 | 0.554 |
| AGAP011792 | AGAP009221 | 0.478 |
| AGAP011792 | AGAP008654 | 0.429 |
| AGAP011792 | AGAP008364 | 0.556 |
| AGAP010731 | AGAP010968 | 0.616 |
| AGAP010731 | AGAP029770 | 0.542 |
| AGAP010731 | AGAP004855 | 0.459 |
| AGAP010731 | AGAP009844 | 0.611 |
| AGAP010731 | AGAP001648 | 0.449 |
| AGAP010731 | AGAP003246 | 0.615 |
| AGAP010731 | AGAP003250 | 0.429 |
| AGAP010731 | AGAP002270 | 0.436 |
| AGAP010731 | AGAP012034 | 0.527 |
| AGAP010731 | AGAP004318 | 0.457 |
| AGAP010731 | AGAP003689 | 0.443 |
| AGAP010731 | AGAP004719 | 0.453 |
| AGAP010731 | AGAP010530 | 0.542 |
| AGAP010731 | AGAP028728 | 0.564 |
| AGAP010731 | AGAP005335 | 0.615 |
| AGAP010731 | AGAP005334 | 0.601 |
| AGAP010731 | AGAP006348 | 0.558 |
| AGAP010731 | AGAP007039 | 0.497 |
| AGAP010731 | AGAP012616 | 0.472 |
| AGAP010731 | AGAP004977 | 0.523 |
| AGAP010731 | AGAP005625 | 0.486 |
| AGAP010731 | AGAP001377 | 0.535 |
| AGAP010731 | AGAP006910 | 0.537 |
| AGAP010731 | AGAP010815 | 0.534 |
| AGAP010731 | AGAP008654 | 0.565 |
| AGAP010731 | AGAP008364 | 0.456 |
| AGAP010731 | AGAP010816 | 0.526 |
| AGAP010968 | AGAP003251 | 0.464 |
| AGAP010968 | AGAP029770 | 0.476 |
| AGAP010968 | AGAP009217 | 0.488 |
| AGAP010968 | AGAP004855 | 0.522 |
| AGAP010968 | AGAP009844 | 0.551 |
| AGAP010968 | AGAP003246 | 0.547 |
| AGAP010968 | AGAP003250 | 0.529 |
| AGAP010968 | AGAP003057 | 0.456 |
| AGAP010968 | AGAP012034 | 0.504 |
| AGAP010968 | AGAP004318 | 0.568 |
| AGAP010968 | AGAP004719 | 0.532 |
| AGAP010968 | AGAP002813 | 0.436 |
| AGAP010968 | AGAP010545 | 0.543 |
| AGAP010968 | AGAP009252 | 0.447 |
| AGAP010968 | AGAP010530 | 0.589 |
| AGAP010968 | AGAP028728 | 0.612 |
| AGAP010968 | AGAP011785 | 0.443 |
| AGAP010968 | AGAP005335 | 0.625 |
| AGAP010968 | AGAP005334 | 0.642 |
| AGAP010968 | AGAP006348 | 0.632 |
| AGAP010968 | AGAP007039 | 0.527 |

| | | |
|---|---|---|
| AGAP011790 | AGAP029770 | 0.460 |
| AGAP011790 | AGAP009217 | 0.471 |
| AGAP011790 | AGAP009844 | 0.494 |
| AGAP011790 | AGAP003246 | 0.440 |
| AGAP010968 | AGAP012616 | 0.537 |
| AGAP010968 | AGAP004977 | 0.513 |
| AGAP010968 | AGAP005625 | 0.615 |
| AGAP010968 | AGAP028725 | 0.481 |
| AGAP010968 | AGAP001377 | 0.591 |
| AGAP010968 | AGAP006911 | 0.453 |
| AGAP010968 | AGAP006910 | 0.488 |
| AGAP010968 | AGAP009221 | 0.435 |
| AGAP010968 | AGAP008654 | 0.446 |
| AGAP010968 | AGAP008364 | 0.450 |
| AGAP010968 | AGAP010816 | 0.446 |
| AGAP003251 | AGAP004855 | 0.510 |
| AGAP003251 | AGAP009844 | 0.430 |
| AGAP003251 | AGAP003250 | 0.650 |
| AGAP003251 | AGAP003057 | 0.434 |
| AGAP003251 | AGAP004318 | 0.492 |
| AGAP003251 | AGAP000573 | 0.489 |
| AGAP003251 | AGAP000315 | 0.488 |
| AGAP003251 | AGAP002813 | 0.441 |
| AGAP003251 | AGAP004977 | 0.450 |
| AGAP003251 | AGAP001377 | 0.481 |
| AGAP003251 | AGAP006911 | 0.484 |
| AGAP003251 | AGAP008364 | 0.431 |
| AGAP029770 | AGAP004855 | 0.495 |
| AGAP029770 | AGAP009844 | 0.449 |
| AGAP029770 | AGAP003246 | 0.478 |
| AGAP029770 | AGAP003250 | 0.497 |
| AGAP029770 | AGAP002270 | 0.548 |
| AGAP029770 | AGAP012034 | 0.494 |
| AGAP029770 | AGAP000573 | 0.493 |
| AGAP029770 | AGAP002813 | 0.453 |
| AGAP029770 | AGAP010530 | 0.444 |
| AGAP029770 | AGAP028728 | 0.478 |
| AGAP029770 | AGAP005335 | 0.459 |
| AGAP029770 | AGAP005334 | 0.541 |
| AGAP029770 | AGAP006348 | 0.487 |
| AGAP029770 | AGAP007039 | 0.472 |
| AGAP029770 | AGAP012616 | 0.450 |
| AGAP029770 | AGAP004977 | 0.459 |
| AGAP029770 | AGAP005625 | 0.441 |
| AGAP029770 | AGAP001377 | 0.431 |
| AGAP029770 | AGAP001376 | 0.448 |
| AGAP029770 | AGAP010815 | 0.477 |
| AGAP009214 | AGAP009215 | 0.457 |
| AGAP009214 | AGAP003250 | 0.440 |
| AGAP009214 | AGAP004148 | 0.488 |
| AGAP009214 | AGAP007692 | 0.433 |
| AGAP009214 | AGAP009213 | 0.587 |
| AGAP009217 | AGAP012034 | 0.446 |
| AGAP009217 | AGAP010545 | 0.491 |
| AGAP009217 | AGAP012502 | 0.506 |
| AGAP009217 | AGAP010530 | 0.469 |
| AGAP009217 | AGAP028728 | 0.446 |
| AGAP009217 | AGAP007457 | 0.444 |
| AGAP009217 | AGAP008368 | 0.445 |
| AGAP004855 | AGAP009844 | 0.500 |
| AGAP004855 | AGAP003246 | 0.440 |
| AGAP004855 | AGAP003250 | 0.497 |
| AGAP004855 | AGAP003057 | 0.429 |
| AGAP004855 | AGAP004318 | 0.431 |
| AGAP004855 | AGAP003689 | 0.430 |
| AGAP004855 | AGAP004719 | 0.447 |
| AGAP004855 | AGAP002813 | 0.472 |
| AGAP004855 | AGAP028728 | 0.445 |
| AGAP004855 | AGAP005334 | 0.439 |
| AGAP004855 | AGAP006348 | 0.498 |
| AGAP004855 | AGAP004977 | 0.534 |
| AGAP004855 | AGAP005625 | 0.472 |

| | | |
|---|---|---|
| AGAP010730 | AGAP005335 | 0.567 |
| AGAP010730 | AGAP005334 | 0.564 |
| AGAP010730 | AGAP006348 | 0.627 |
| AGAP010730 | AGAP007039 | 0.488 |
| AGAP004855 | AGAP006910 | 0.437 |
| AGAP004855 | AGAP010815 | 0.438 |
| AGAP004855 | AGAP008364 | 0.441 |
| AGAP010833 | AGAP012022 | 0.444 |
| AGAP010833 | AGAP028069 | 0.439 |
| AGAP009844 | AGAP001648 | 0.457 |
| AGAP009844 | AGAP003246 | 0.512 |
| AGAP009844 | AGAP013487 | 0.433 |
| AGAP009844 | AGAP003250 | 0.474 |
| AGAP009844 | AGAP003057 | 0.454 |
| AGAP009844 | AGAP012034 | 0.531 |
| AGAP009844 | AGAP004318 | 0.466 |
| AGAP009844 | AGAP004719 | 0.509 |
| AGAP009844 | AGAP002422 | 0.458 |
| AGAP009844 | AGAP002813 | 0.532 |
| AGAP009844 | AGAP010545 | 0.437 |
| AGAP009844 | AGAP010530 | 0.447 |
| AGAP009844 | AGAP028728 | 0.448 |
| AGAP009844 | AGAP005335 | 0.595 |
| AGAP009844 | AGAP005334 | 0.621 |
| AGAP009844 | AGAP006348 | 0.578 |
| AGAP009844 | AGAP004977 | 0.505 |
| AGAP009844 | AGAP005625 | 0.465 |
| AGAP009844 | AGAP028725 | 0.472 |
| AGAP009844 | AGAP001377 | 0.465 |
| AGAP009844 | AGAP006910 | 0.486 |
| AGAP009844 | AGAP010815 | 0.506 |
| AGAP009844 | AGAP008654 | 0.448 |
| AGAP009844 | AGAP010816 | 0.473 |
| AGAP009263 | AGAP007454 | 0.442 |
| AGAP009263 | AGAP010832 | 0.481 |
| AGAP001648 | AGAP002422 | 0.431 |
| AGAP001648 | AGAP005625 | 0.459 |
| AGAP001648 | AGAP010816 | 0.486 |
| AGAP009215 | AGAP011785 | 0.467 |
| AGAP009215 | AGAP007457 | 0.461 |
| AGAP003247 | AGAP004318 | 0.509 |
| AGAP003247 | AGAP010545 | 0.491 |
| AGAP003247 | AGAP008403 | 0.494 |
| AGAP003247 | AGAP009252 | 0.517 |
| AGAP003247 | AGAP028728 | 0.451 |
| AGAP003247 | AGAP005335 | 0.433 |
| AGAP003247 | AGAP012616 | 0.446 |
| AGAP003247 | AGAP003194 | 0.474 |
| AGAP003247 | AGAP010815 | 0.576 |
| AGAP003246 | AGAP003250 | 0.435 |
| AGAP003246 | AGAP012034 | 0.512 |
| AGAP003246 | AGAP004318 | 0.496 |
| AGAP003246 | AGAP004719 | 0.485 |
| AGAP003246 | AGAP010545 | 0.475 |
| AGAP003246 | AGAP010530 | 0.538 |
| AGAP003246 | AGAP028728 | 0.535 |
| AGAP003246 | AGAP005335 | 0.475 |
| AGAP003246 | AGAP005334 | 0.476 |
| AGAP003246 | AGAP006348 | 0.440 |
| AGAP003246 | AGAP007039 | 0.443 |
| AGAP003246 | AGAP007454 | 0.490 |
| AGAP003246 | AGAP012616 | 0.505 |
| AGAP003246 | AGAP004977 | 0.488 |
| AGAP003246 | AGAP010815 | 0.441 |
| AGAP003246 | AGAP008654 | 0.520 |
| AGAP003246 | AGAP008368 | 0.433 |
| AGAP003246 | AGAP010816 | 0.447 |
| AGAP012037 | AGAP012034 | 0.571 |
| AGAP012037 | AGAP005335 | 0.429 |
| AGAP012037 | AGAP005334 | 0.456 |
| AGAP012037 | AGAP010816 | 0.497 |
| AGAP013184 | AGAP003250 | 0.503 |

| | | |
|---|---|---|
| AGAP011792 | AGAP003247 | 0.476 |
| AGAP011792 | AGAP003246 | 0.449 |
| AGAP011792 | AGAP003250 | 0.564 |
| AGAP011792 | AGAP004148 | 0.446 |
| AGAP013184 | AGAP002813 | 0.502 |
| AGAP013184 | AGAP008091 | 0.461 |
| AGAP013184 | AGAP009252 | 0.456 |
| AGAP013184 | AGAP009273 | 0.560 |
| AGAP013487 | AGAP029074 | 0.544 |
| AGAP013487 | AGAP007408 | 0.477 |
| AGAP013487 | AGAP004980 | 0.487 |
| AGAP013487 | AGAP010815 | 0.463 |
| AGAP003249 | AGAP007454 | 0.489 |
| AGAP003250 | AGAP003057 | 0.478 |
| AGAP003250 | AGAP004318 | 0.566 |
| AGAP003250 | AGAP000573 | 0.572 |
| AGAP003250 | AGAP000315 | 0.463 |
| AGAP003250 | AGAP004719 | 0.497 |
| AGAP003250 | AGAP002813 | 0.479 |
| AGAP003250 | AGAP028728 | 0.451 |
| AGAP003250 | AGAP005334 | 0.449 |
| AGAP003250 | AGAP006348 | 0.458 |
| AGAP003250 | AGAP007039 | 0.446 |
| AGAP003250 | AGAP012616 | 0.505 |
| AGAP003250 | AGAP004977 | 0.538 |
| AGAP003250 | AGAP005625 | 0.477 |
| AGAP003250 | AGAP001377 | 0.463 |
| AGAP003250 | AGAP007692 | 0.510 |
| AGAP003250 | AGAP001376 | 0.446 |
| AGAP003250 | AGAP006911 | 0.548 |
| AGAP003250 | AGAP006910 | 0.437 |
| AGAP003250 | AGAP008364 | 0.523 |
| AGAP004149 | AGAP009000 | 0.470 |
| AGAP004149 | AGAP011040 | 0.430 |
| AGAP004149 | AGAP011782 | 0.456 |
| AGAP004149 | AGAP012502 | 0.489 |
| AGAP004149 | AGAP028069 | 0.513 |
| AGAP004149 | AGAP028183 | 0.566 |
| AGAP004149 | AGAP028229 | 0.446 |
| AGAP004149 | AGAP028102 | 0.465 |
| AGAP009211 | AGAP002811 | 0.429 |
| AGAP009211 | AGAP002625 | 0.537 |
| AGAP009211 | AGAP007280 | 0.460 |
| AGAP009211 | AGAP010818 | 0.446 |
| AGAP009211 | AGAP010832 | 0.571 |
| AGAP009211 | AGAP010831 | 0.543 |
| AGAP009220 | AGAP009252 | 0.439 |
| AGAP011325 | AGAP012022 | 0.471 |
| AGAP011325 | AGAP010830 | 0.444 |
| AGAP009849 | AGAP007037 | 0.451 |
| AGAP009849 | AGAP008654 | 0.464 |
| AGAP009849 | AGAP008368 | 0.478 |
| AGAP003686 | AGAP007691 | 0.467 |
| AGAP002270 | AGAP002813 | 0.444 |
| AGAP002270 | AGAP005334 | 0.538 |
| AGAP002270 | AGAP005625 | 0.437 |
| AGAP002270 | AGAP001376 | 0.458 |
| AGAP002270 | AGAP009221 | 0.449 |
| AGAP003057 | AGAP003689 | 0.458 |
| AGAP003057 | AGAP029093 | 0.490 |
| AGAP003057 | AGAP006348 | 0.454 |
| AGAP003057 | AGAP001377 | 0.469 |
| AGAP003057 | AGAP006911 | 0.436 |
| AGAP003057 | AGAP006910 | 0.433 |
| AGAP003057 | AGAP008654 | 0.457 |
| AGAP003057 | AGAP008364 | 0.461 |
| AGAP012034 | AGAP004719 | 0.429 |
| AGAP012034 | AGAP002813 | 0.493 |
| AGAP012034 | AGAP010545 | 0.450 |
| AGAP012034 | AGAP010530 | 0.501 |
| AGAP012034 | AGAP028728 | 0.551 |
| AGAP012034 | AGAP005335 | 0.518 |

| | | |
|---|---|---|
| AGAP010968 | AGAP027997 | 0.431 |
| AGAP010968 | AGAP007457 | 0.451 |
| AGAP010968 | AGAP007453 | 0.455 |
| AGAP012034 | AGAP012616 | 0.465 |
| AGAP012034 | AGAP028725 | 0.550 |
| AGAP012034 | AGAP007692 | 0.437 |
| AGAP012034 | AGAP010815 | 0.438 |
| AGAP012034 | AGAP008654 | 0.518 |
| AGAP012034 | AGAP010816 | 0.557 |
| AGAP028007 | AGAP012020 | 0.491 |
| AGAP028007 | AGAP029047 | 0.471 |
| AGAP028007 | AGAP007034 | 0.566 |
| AGAP028007 | AGAP005496 | 0.434 |
| AGAP028167 | AGAP002625 | 0.473 |
| AGAP028167 | AGAP010819 | 0.439 |
| AGAP028167 | AGAP010832 | 0.435 |
| AGAP004318 | AGAP004719 | 0.497 |
| AGAP004318 | AGAP010530 | 0.474 |
| AGAP004318 | AGAP028728 | 0.491 |
| AGAP004318 | AGAP005335 | 0.517 |
| AGAP004318 | AGAP005334 | 0.532 |
| AGAP004318 | AGAP007454 | 0.510 |
| AGAP004318 | AGAP012616 | 0.475 |
| AGAP004318 | AGAP004977 | 0.513 |
| AGAP004318 | AGAP005625 | 0.536 |
| AGAP004318 | AGAP001377 | 0.546 |
| AGAP004318 | AGAP007692 | 0.442 |
| AGAP004318 | AGAP006911 | 0.462 |
| AGAP004318 | AGAP009221 | 0.476 |
| AGAP004318 | AGAP008364 | 0.443 |
| AGAP000573 | AGAP000315 | 0.451 |
| AGAP000573 | AGAP028728 | 0.433 |
| AGAP000573 | AGAP007692 | 0.532 |
| AGAP000315 | AGAP003689 | 0.445 |
| AGAP000315 | AGAP002813 | 0.465 |
| AGAP000315 | AGAP027997 | 0.443 |
| AGAP000315 | AGAP007457 | 0.435 |
| AGAP003689 | AGAP006348 | 0.465 |
| AGAP003689 | AGAP027997 | 0.447 |
| AGAP003689 | AGAP008654 | 0.442 |
| AGAP003689 | AGAP008364 | 0.497 |
| AGAP003689 | AGAP010816 | 0.435 |
| AGAP004719 | AGAP002813 | 0.447 |
| AGAP004719 | AGAP010545 | 0.453 |
| AGAP004719 | AGAP009252 | 0.547 |
| AGAP004719 | AGAP005335 | 0.495 |
| AGAP004719 | AGAP005334 | 0.480 |
| AGAP004719 | AGAP006348 | 0.443 |
| AGAP004719 | AGAP004977 | 0.488 |
| AGAP004719 | AGAP001377 | 0.446 |
| AGAP004719 | AGAP006910 | 0.465 |
| AGAP002422 | AGAP029093 | 0.432 |
| AGAP002422 | AGAP029074 | 0.464 |
| AGAP008995 | AGAP009000 | 0.557 |
| AGAP008995 | AGAP008996 | 0.592 |
| AGAP008995 | AGAP001433 | 0.502 |
| AGAP008995 | AGAP008998 | 0.567 |
| AGAP008995 | AGAP002784 | 0.478 |
| AGAP008995 | AGAP011040 | 0.466 |
| AGAP008995 | AGAP011782 | 0.450 |
| AGAP008995 | AGAP012502 | 0.506 |
| AGAP008995 | AGAP009316 | 0.440 |
| AGAP008995 | AGAP000940 | 0.441 |
| AGAP008995 | AGAP029559 | 0.516 |
| AGAP008995 | AGAP007408 | 0.443 |
| AGAP008995 | AGAP002825 | 0.519 |
| AGAP008995 | AGAP004980 | 0.490 |
| AGAP009000 | AGAP008996 | 0.520 |
| AGAP009000 | AGAP008998 | 0.492 |
| AGAP009000 | AGAP002784 | 0.442 |
| AGAP009000 | AGAP011040 | 0.447 |
| AGAP009000 | AGAP012502 | 0.532 |
| AGAP009000 | AGAP012504 | 0.459 |
| AGAP004855 | AGAP028725 | 0.455 |
| AGAP004855 | AGAP001377 | 0.560 |
| AGAP004855 | AGAP006911 | 0.531 |
| AGAP009000 | AGAP000123 | 0.461 |
| AGAP009000 | AGAP004980 | 0.501 |
| AGAP009006 | AGAP001433 | 0.507 |
| AGAP009006 | AGAP002784 | 0.442 |
| AGAP009006 | AGAP012502 | 0.447 |
| AGAP009006 | AGAP000123 | 0.446 |
| AGAP008183 | AGAP008996 | 0.456 |
| AGAP008183 | AGAP008998 | 0.467 |
| AGAP008183 | AGAP007410 | 0.444 |
| AGAP008183 | AGAP029559 | 0.430 |
| AGAP008183 | AGAP002825 | 0.466 |
| AGAP008183 | AGAP004975 | 0.457 |
| AGAP008183 | AGAP001375 | 0.435 |
| AGAP013089 | AGAP011782 | 0.473 |
| AGAP013089 | AGAP007408 | 0.465 |
| AGAP013089 | AGAP004980 | 0.474 |
| AGAP013089 | AGAP006631 | 0.430 |
| AGAP013089 | AGAP010832 | 0.505 |
| AGAP013089 | AGAP010831 | 0.531 |
| AGAP008996 | AGAP001433 | 0.432 |
| AGAP008996 | AGAP012502 | 0.517 |
| AGAP008996 | AGAP029559 | 0.449 |
| AGAP008996 | AGAP002825 | 0.588 |
| AGAP008996 | AGAP004975 | 0.435 |
| AGAP008996 | AGAP004980 | 0.455 |
| AGAP008996 | AGAP001375 | 0.465 |
| AGAP008996 | AGAP008407 | 0.452 |
| AGAP001433 | AGAP002784 | 0.510 |
| AGAP001433 | AGAP012502 | 0.560 |
| AGAP001433 | AGAP002625 | 0.439 |
| AGAP001433 | AGAP010708 | 0.516 |
| AGAP001433 | AGAP000123 | 0.496 |
| AGAP001433 | AGAP004980 | 0.456 |
| AGAP001433 | AGAP010818 | 0.456 |
| AGAP002811 | AGAP012021 | 0.460 |
| AGAP002813 | AGAP009252 | 0.470 |
| AGAP002813 | AGAP005334 | 0.465 |
| AGAP002813 | AGAP006348 | 0.434 |
| AGAP002813 | AGAP004977 | 0.478 |
| AGAP002813 | AGAP028725 | 0.533 |
| AGAP008998 | AGAP007410 | 0.462 |
| AGAP008998 | AGAP007408 | 0.494 |
| AGAP002784 | AGAP029093 | 0.503 |
| AGAP002784 | AGAP012502 | 0.552 |
| AGAP002784 | AGAP010708 | 0.458 |
| AGAP010545 | AGAP009252 | 0.579 |
| AGAP010545 | AGAP010530 | 0.538 |
| AGAP010545 | AGAP028728 | 0.604 |
| AGAP010545 | AGAP005335 | 0.467 |
| AGAP010545 | AGAP012616 | 0.492 |
| AGAP010545 | AGAP004977 | 0.442 |
| AGAP010545 | AGAP008654 | 0.475 |
| AGAP010545 | AGAP008368 | 0.478 |
| AGAP003691 | AGAP009251 | 0.507 |
| AGAP003691 | AGAP009252 | 0.496 |
| AGAP029093 | AGAP009252 | 0.454 |
| AGAP029093 | AGAP011782 | 0.467 |
| AGAP029093 | AGAP029100 | 0.435 |
| AGAP029093 | AGAP028641 | 0.476 |
| AGAP029093 | AGAP000929 | 0.496 |
| AGAP029093 | AGAP027997 | 0.484 |
| AGAP029093 | AGAP007457 | 0.531 |
| AGAP029093 | AGAP001798 | 0.526 |
| AGAP029093 | AGAP028725 | 0.469 |
| AGAP029093 | AGAP007692 | 0.517 |
| AGAP008403 | AGAP009251 | 0.437 |
| AGAP009251 | AGAP009252 | 0.599 |
| AGAP009251 | AGAP028641 | 0.465 |
| AGAP009252 | AGAP028641 | 0.502 |
| AGAP009252 | AGAP010530 | 0.464 |
| AGAP013184 | AGAP012034 | 0.481 |
| AGAP013184 | AGAP000573 | 0.512 |
| AGAP013184 | AGAP000315 | 0.470 |
| AGAP009252 | AGAP006267 | 0.491 |
| AGAP009252 | AGAP012616 | 0.488 |
| AGAP009252 | AGAP004977 | 0.452 |
| AGAP009252 | AGAP005625 | 0.486 |
| AGAP009252 | AGAP028725 | 0.460 |
| AGAP009252 | AGAP008654 | 0.500 |
| AGAP009252 | AGAP008368 | 0.507 |
| AGAP009252 | AGAP010816 | 0.495 |
| AGAP009252 | AGAP010830 | 0.529 |
| AGAP011040 | AGAP011782 | 0.495 |
| AGAP011040 | AGAP011719 | 0.449 |
| AGAP011040 | AGAP012502 | 0.484 |
| AGAP011040 | AGAP009316 | 0.454 |
| AGAP011040 | AGAP004810 | 0.438 |
| AGAP011040 | AGAP010708 | 0.505 |
| AGAP011040 | AGAP002911 | 0.533 |
| AGAP011040 | AGAP002825 | 0.434 |
| AGAP011040 | AGAP004976 | 0.480 |
| AGAP011782 | AGAP029047 | 0.445 |
| AGAP011782 | AGAP000929 | 0.511 |
| AGAP011782 | AGAP007034 | 0.483 |
| AGAP011782 | AGAP006631 | 0.456 |
| AGAP011782 | AGAP010831 | 0.431 |
| AGAP029100 | AGAP028725 | 0.431 |
| AGAP029100 | AGAP007692 | 0.449 |
| AGAP029100 | AGAP006910 | 0.461 |
| AGAP029100 | AGAP003139 | 0.435 |
| AGAP011719 | AGAP012021 | 0.463 |
| AGAP011719 | AGAP012502 | 0.457 |
| AGAP011719 | AGAP009316 | 0.512 |
| AGAP011719 | AGAP010709 | 0.476 |
| AGAP011719 | AGAP000940 | 0.527 |
| AGAP011719 | AGAP010708 | 0.465 |
| AGAP012020 | AGAP028069 | 0.468 |
| AGAP012020 | AGAP028229 | 0.531 |
| AGAP012020 | AGAP028102 | 0.524 |
| AGAP012020 | AGAP000940 | 0.516 |
| AGAP012020 | AGAP007034 | 0.463 |
| AGAP012021 | AGAP012022 | 0.501 |
| AGAP012021 | AGAP012502 | 0.565 |
| AGAP012021 | AGAP028183 | 0.511 |
| AGAP012021 | AGAP012504 | 0.459 |
| AGAP012021 | AGAP004980 | 0.456 |
| AGAP012021 | AGAP010831 | 0.472 |
| AGAP012022 | AGAP028069 | 0.437 |
| AGAP012022 | AGAP028183 | 0.510 |
| AGAP012022 | AGAP012504 | 0.455 |
| AGAP012022 | AGAP004976 | 0.447 |
| AGAP012022 | AGAP004198 | 0.475 |
| AGAP012502 | AGAP004811 | 0.525 |
| AGAP012502 | AGAP009316 | 0.551 |
| AGAP012502 | AGAP004810 | 0.521 |
| AGAP012502 | AGAP002625 | 0.517 |
| AGAP012502 | AGAP006430 | 0.515 |
| AGAP012502 | AGAP029559 | 0.495 |
| AGAP012502 | AGAP010708 | 0.499 |
| AGAP012502 | AGAP002911 | 0.490 |
| AGAP012502 | AGAP002825 | 0.446 |
| AGAP012502 | AGAP004980 | 0.503 |
| AGAP012502 | AGAP004976 | 0.504 |
| AGAP012502 | AGAP004198 | 0.448 |
| AGAP012502 | AGAP010831 | 0.453 |
| AGAP028069 | AGAP028183 | 0.436 |
| AGAP028069 | AGAP028229 | 0.441 |
| AGAP028069 | AGAP028102 | 0.440 |
| AGAP028069 | AGAP005246 | 0.460 |
| AGAP028183 | AGAP000940 | 0.460 |
| AGAP028183 | AGAP010708 | 0.461 |
| AGAP028183 | AGAP002911 | 0.468 |
| AGAP028183 | AGAP007034 | 0.464 |
| AGAP012034 | AGAP005334 | 0.530 |
| AGAP012034 | AGAP006348 | 0.469 |
| AGAP012034 | AGAP007039 | 0.462 |
| AGAP028183 | AGAP004198 | 0.539 |
| AGAP028229 | AGAP028102 | 0.495 |
| AGAP028229 | AGAP028075 | 0.482 |
| AGAP028641 | AGAP011786 | 0.455 |
| AGAP028641 | AGAP027997 | 0.491 |
| AGAP029074 | AGAP007692 | 0.431 |
| AGAP028102 | AGAP028075 | 0.527 |
| AGAP010530 | AGAP028728 | 0.576 |
| AGAP010530 | AGAP005335 | 0.434 |
| AGAP010530 | AGAP005334 | 0.471 |
| AGAP010530 | AGAP007039 | 0.434 |
| AGAP010530 | AGAP012616 | 0.492 |
| AGAP010530 | AGAP004977 | 0.430 |
| AGAP010530 | AGAP009221 | 0.460 |
| AGAP010530 | AGAP008364 | 0.470 |
| AGAP028728 | AGAP005335 | 0.510 |
| AGAP028728 | AGAP005334 | 0.471 |
| AGAP028728 | AGAP006348 | 0.484 |
| AGAP028728 | AGAP007039 | 0.484 |
| AGAP028728 | AGAP012616 | 0.586 |
| AGAP028728 | AGAP004977 | 0.500 |
| AGAP028728 | AGAP028725 | 0.451 |
| AGAP028728 | AGAP008654 | 0.434 |
| AGAP011785 | AGAP005334 | 0.497 |
| AGAP011785 | AGAP012616 | 0.443 |
| AGAP011785 | AGAP005625 | 0.569 |
| AGAP011785 | AGAP009221 | 0.532 |
| AGAP029077 | AGAP007407 | 0.448 |
| AGAP029077 | AGAP010819 | 0.483 |
| AGAP004811 | AGAP004810 | 0.440 |
| AGAP009316 | AGAP000940 | 0.545 |
| AGAP009316 | AGAP010708 | 0.494 |
| AGAP009316 | AGAP004980 | 0.533 |
| AGAP009316 | AGAP010819 | 0.461 |
| AGAP009316 | AGAP010830 | 0.460 |
| AGAP005335 | AGAP005334 | 0.677 |
| AGAP005335 | AGAP006348 | 0.538 |
| AGAP005335 | AGAP007454 | 0.461 |
| AGAP005335 | AGAP004977 | 0.433 |
| AGAP005335 | AGAP005625 | 0.508 |
| AGAP005335 | AGAP028725 | 0.463 |
| AGAP005335 | AGAP010815 | 0.450 |
| AGAP005335 | AGAP008654 | 0.468 |
| AGAP005335 | AGAP010816 | 0.499 |
| AGAP029047 | AGAP006631 | 0.533 |
| AGAP006267 | AGAP007454 | 0.505 |
| AGAP006267 | AGAP012938 | 0.494 |
| AGAP000940 | AGAP010708 | 0.488 |
| AGAP000940 | AGAP002911 | 0.433 |
| AGAP000940 | AGAP007034 | 0.471 |
| AGAP000940 | AGAP010814 | 0.470 |
| AGAP002625 | AGAP010831 | 0.528 |
| AGAP006430 | AGAP001375 | 0.444 |
| AGAP007411 | AGAP007412 | 0.442 |
| AGAP005334 | AGAP006348 | 0.605 |
| AGAP005334 | AGAP004977 | 0.443 |
| AGAP005334 | AGAP005625 | 0.623 |
| AGAP005334 | AGAP028725 | 0.621 |
| AGAP005334 | AGAP001377 | 0.429 |
| AGAP005334 | AGAP001376 | 0.437 |
| AGAP005334 | AGAP009221 | 0.561 |
| AGAP005334 | AGAP010815 | 0.510 |
| AGAP005334 | AGAP010816 | 0.555 |
| AGAP007407 | AGAP007455 | 0.439 |
| AGAP007407 | AGAP007454 | 0.489 |
| AGAP007410 | AGAP007408 | 0.527 |
| AGAP029559 | AGAP002825 | 0.500 |
| AGAP029559 | AGAP004980 | 0.533 |
| AGAP010708 | AGAP002911 | 0.459 |
| AGAP010708 | AGAP004980 | 0.474 |

| | | | | | | | | | | | |
|---|---|---|---|---|---|---|---|---|---|---|---|
| AGAP009000 | AGAP029559 | 0.541 | AGAP009252 | AGAP028728 | 0.482 | AGAP028183 | AGAP002825 | 0.455 | AGAP010708 | AGAP004198 | 0.488 |
| AGAP010708 | AGAP010819 | 0.528 | AGAP007280 | AGAP010818 | 0.456 | AGAP006348 | AGAP004977 | 0.478 | AGAP028725 | AGAP010816 | 0.476 |
| AGAP010708 | AGAP010818 | 0.540 | AGAP007280 | AGAP010830 | 0.501 | AGAP006348 | AGAP005625 | 0.529 | AGAP012938 | AGAP009212 | 0.447 |
| AGAP010708 | AGAP010814 | 0.504 | AGAP005625 | AGAP028725 | 0.591 | AGAP006348 | AGAP028725 | 0.561 | AGAP012938 | AGAP003139 | 0.595 |
| AGAP010708 | AGAP010830 | 0.475 | AGAP005625 | AGAP001377 | 0.523 | AGAP006348 | AGAP001377 | 0.542 | AGAP012938 | AGAP008366 | 0.438 |
| AGAP007408 | AGAP009213 | 0.479 | AGAP005625 | AGAP006910 | 0.440 | AGAP006348 | AGAP006911 | 0.472 | AGAP006909 | AGAP006910 | 0.430 |
| AGAP007408 | AGAP007693 | 0.505 | AGAP005625 | AGAP009221 | 0.593 | AGAP006348 | AGAP006910 | 0.466 | AGAP006909 | AGAP009670 | 0.469 |
| AGAP007408 | AGAP010830 | 0.438 | AGAP005625 | AGAP010815 | 0.437 | AGAP006348 | AGAP010815 | 0.445 | AGAP001377 | AGAP006911 | 0.553 |
| AGAP002911 | AGAP004980 | 0.467 | AGAP005625 | AGAP010816 | 0.504 | AGAP006348 | AGAP008654 | 0.443 | AGAP007692 | AGAP010815 | 0.519 |
| AGAP000929 | AGAP004980 | 0.483 | AGAP028725 | AGAP001377 | 0.430 | AGAP007455 | AGAP007454 | 0.650 | AGAP001376 | AGAP009221 | 0.475 |
| AGAP007455 | AGAP007456 | 0.642 | AGAP006327 | AGAP006416 | 0.459 | AGAP012616 | AGAP005625 | 0.437 | AGAP009221 | AGAP009212 | 0.471 |
| AGAP007455 | AGAP007453 | 0.656 | AGAP007454 | AGAP007456 | 0.634 | AGAP012616 | AGAP007692 | 0.440 | AGAP009221 | AGAP010815 | 0.446 |
| AGAP007455 | AGAP004978 | 0.451 | AGAP007454 | AGAP007453 | 0.661 | AGAP012616 | AGAP009221 | 0.446 | AGAP007693 | AGAP003194 | 0.442 |
| AGAP007034 | AGAP004975 | 0.445 | AGAP007454 | AGAP004978 | 0.589 | AGAP012616 | AGAP010815 | 0.440 | AGAP010815 | AGAP010816 | 0.487 |
| AGAP007034 | AGAP004980 | 0.442 | AGAP007454 | AGAP010815 | 0.453 | AGAP004977 | AGAP005625 | 0.453 | AGAP010819 | AGAP010818 | 0.659 |
| AGAP007034 | AGAP006631 | 0.484 | AGAP007456 | AGAP007453 | 0.596 | AGAP004977 | AGAP006910 | 0.430 | AGAP010819 | AGAP010832 | 0.498 |
| AGAP007034 | AGAP010818 | 0.499 | AGAP007456 | AGAP004978 | 0.432 | AGAP004977 | AGAP008654 | 0.432 | AGAP010819 | AGAP010814 | 0.529 |
| AGAP005496 | AGAP004980 | 0.437 | AGAP002825 | AGAP004975 | 0.509 | AGAP004977 | AGAP008364 | 0.453 | AGAP010819 | AGAP010830 | 0.672 |
| AGAP007045 | AGAP008368 | 0.464 | AGAP002825 | AGAP004980 | 0.490 | AGAP004977 | AGAP010816 | 0.489 | AGAP010818 | AGAP010814 | 0.562 |
| AGAP007037 | AGAP008654 | 0.513 | AGAP002825 | AGAP001375 | 0.478 | AGAP004980 | AGAP007692 | 0.439 | AGAP010818 | AGAP010830 | 0.637 |
| AGAP007037 | AGAP008368 | 0.573 | AGAP002825 | AGAP004198 | 0.440 | AGAP004980 | AGAP007691 | 0.440 | AGAP008654 | AGAP008368 | 0.535 |
| AGAP007039 | AGAP012616 | 0.476 | AGAP006258 | AGAP004975 | 0.497 | AGAP004980 | AGAP007693 | 0.460 | AGAP008654 | AGAP010816 | 0.526 |
| AGAP007039 | AGAP004977 | 0.493 | AGAP006258 | AGAP004981 | 0.454 | AGAP004980 | AGAP010818 | 0.467 | AGAP008654 | AGAP010812 | 0.430 |
| AGAP007039 | AGAP006910 | 0.475 | AGAP006258 | AGAP004980 | 0.479 | AGAP004980 | AGAP010831 | 0.482 | AGAP010832 | AGAP008366 | 0.509 |
| AGAP007039 | AGAP010815 | 0.446 | AGAP006258 | AGAP007693 | 0.431 | AGAP004976 | AGAP004198 | 0.497 | AGAP010832 | AGAP010831 | 0.677 |
| AGAP027997 | AGAP007457 | 0.461 | AGAP004975 | AGAP004980 | 0.510 | AGAP004976 | AGAP010831 | 0.497 | AGAP008366 | AGAP010831 | 0.439 |
| AGAP027997 | AGAP004980 | 0.471 | AGAP004975 | AGAP001375 | 0.494 | AGAP007280 | AGAP010819 | 0.447 | AGAP010816 | AGAP010812 | 0.484 |
| AGAP027997 | AGAP008368 | 0.444 | AGAP012616 | AGAP004977 | 0.596 | AGAP006910 | AGAP008364 | 0.483 | AGAP010814 | AGAP010830 | 0.631 |

**Table S17: Node table of AgMelGCN2.0** Table presents node-associated data, including AGAP number, gene name, immune family membership, phenotype in CLIP-RNAi screen (this study), memberships in network community and network core, as well as four network centrality measures (strength, eigenvector, closeness, and betweenness).

| AGAP | Label | name | family | subfamily | phenotype | community | core | strength | eigenvector | closeness | betweenness |
|---|---|---|---|---|---|---|---|---|---|---|---|
| AGAP000123 | CTL | CTLSE2 | CTL | CTL | not a clip | 1 | 0 | 1.856 | 0.000049 | 25.43811 | 0 |
| AGAP001433 | cSP_D | CLIPD3 | cSP | cSP_D | positive_A | 1 | 0 | 5.409 | 0.00037 | 33.74858 | 0.00961 |
| AGAP002784 | cSP_D | CLIPD8 | cSP | cSP_D | no phenotype | 1 | 0 | 3.835 | 0.00081 | 33.68684 | 0.014416 |
| AGAP002811 | cSP_D | CLIPD4 | cSP | cSP_D | no phenotype | 1 | 0 | 0.889 | 0.000016 | 24.33237 | 0 |
| AGAP002815 | cSPH_A | CLIPA15 | cSPH | cSPH_A | no phenotype | 1 | 0 | 5.604 | 0.000374 | 33.75459 | 0.003506 |
| AGAP000929 | CTL | CTLSE1 | CTL | CTL | not a clip | 1 | 0 | 1.49 | 0.000723 | 31.0577 | 0.001299 |
| AGAP000940 | CTL | CTL7 | CTL | CTL | not a clip | 1 | 0 | 5.26 | 0.000115 | 30.39393 | 0.008117 |
| AGAP001375 | SRPN | SRPN12 | SRPN | SRPN | not a clip | 1 | 0 | 2.316 | 0.000041 | 24.9128 | 0.000195 |
| AGAP003686 | cSP_B | CLIPB47 | cSP | cSP_B | no phenotype | 1 | 0 | 0.467 | 0.000004 | 19.79513 | 0 |
| AGAP004149 | cSP_B | CLIPB41 | cSP | cSP_B | no phenotype | 1 | 0 | 3.835 | 0.00017 | 31.12575 | 0.040974 |
| AGAP006954 | cSPH_A | CLIPA10 | cSPH | cSPH_A | no phenotype | 1 | 0 | 7.461 | 0.000453 | 35.45673 | 0.02039 |
| AGAP008183 | cSP_D | CLIPD2 | cSP | cSP_D | no phenotype | 1 | 0 | 3.602 | 0.000067 | 25.88951 | 0.000844 |
| AGAP008995 | cSP_D | CLIPD12 | cSP | cSP_D | no phenotype | 1 | 0 | 8.189 | 0.000464 | 35.04082 | 0.004675 |
| AGAP002625 | CTL | CTL9 | CTL | CTL | not a clip | 1 | 0 | 2.494 | 0.000159 | 29.88545 | 0.006039 |
| AGAP008996 | cSP_D | CLIPD22 | cSP | cSP_D | no phenotype | 1 | 0 | 6.464 | 0.000357 | 33.1015 | 0.016169 |
| AGAP008998 | cSP_D | CLIPD7 | cSP | cSP_D | no phenotype | 1 | 0 | 3.495 | 0.000099 | 28.97886 | 0.001169 |
| AGAP009000 | cSP_D | CLIPD13 | cSP | cSP_D | no phenotype | 1 | 0 | 6.907 | 0.000411 | 34.41841 | 0.007468 |
| AGAP009006 | cSPH_D | CLIPD14 | cSPH | cSPH_D | no phenotype | 1 | 0 | 1.842 | 0.000151 | 27.66281 | 0 |
| AGAP002825 | PPO | PPO1 | PPO | PPO | not a clip | 1 | 0 | 6.361 | 0.000334 | 32.78054 | 0.002792 |
| AGAP002911 | CTL | CTLMA9 | CTL | CTL | not a clip | 1 | 0 | 2.85 | 0.000263 | 31.04089 | 0.000714 |
| AGAP009211 | cSP_B | CLIPB42 | cSP | cSP_B | not tested | 1 | 0 | 2.986 | 0.000118 | 28.3568 | 0.004026 |
| AGAP003194 | SRPN | SRPN8 | SRPN | SRPN | not a clip | 1 | 0 | 0.916 | 0.001499 | 26.36818 | 0.001753 |
| AGAP009263 | cSPH_B | CLIPB16 | cSPH | cSPH_B | no phenotype | 1 | 0 | 1.398 | 0.00256 | 28.10365 | 0.00513 |
| AGAP010833 | cSP_B | CLIPB14 | cSP | cSP_B | positive_A | 1 | 0 | 0.883 | 0.000002 | 20.8387 | 0 |
| AGAP011040 | cSP_E | CLIPE19 | cSP | cSP_E | no phenotype | 1 | 0 | 6.17 | 0.000272 | 32.87134 | 0.002532 |
| AGAP011325 | cSP_B | CLIPB45 | cSP | cSP_B | not tested | 1 | 0 | 0.915 | 0.000122 | 25.82005 | 0.003312 |
| AGAP011719 | cSP_E | CLIPE21 | cSP | cSP_E | no phenotype | 1 | 0 | 3.939 | 0.00019 | 30.42929 | 0.011688 |
| AGAP011782 | cSPH_E | CLIPE2 | cSPH | cSPH_E | not tested | 1 | 0 | 5.661 | 0.000671 | 34.25887 | 0.032987 |
| AGAP011783 | cSPH_A | CLIPA13 | cSPH | cSPH_A | negative_M | 1 | 0 | 1.475 | 0.004217 | 29.79542 | 0.018896 |
| AGAP011793 | cSPH_A | CLIPA31 | cSPH | cSPH_A | not tested | 1 | 0 | 4.513 | 0.000356 | 32.70979 | 0.01513 |
| AGAP011794 | cSPH_A | CLIPA32 | cSPH | cSPH_A | not tested | 1 | 0 | 3.329 | 0.000279 | 31.59638 | 0.002597 |
| AGAP012020 | cSP_E | CLIPE22 | cSP | cSP_E | no phenotype | 1 | 0 | 2.993 | 0.00002 | 26.57377 | 0.011429 |
| AGAP012021 | cSPH_E | CLIPE23 | cSPH | cSPH_E | not tested | 1 | 0 | 5.331 | 0.000335 | 34.02823 | 0.031558 |
| AGAP004198 | SRPN | SRPN19 | SRPN | SRPN | not a clip | 1 | 0 | 2.887 | 0.00015 | 29.55783 | 0.000195 |
| AGAP012022 | cSP_E | CLIPE24 | cSP | cSP_E | not tested | 1 | 0 | 3.74 | 0.000056 | 28.01932 | 0.013442 |
| AGAP012502 | cSP_E | CLIPE25 | cSP | cSP_E | no phenotype | 1 | 0 | 14.17 | 0.003006 | 41.98184 | 0.20039 |
| AGAP004810 | CTL | CTL3 | CTL | CTL | not a clip | 1 | 0 | 1.399 | 0.000138 | 28.12771 | 0 |
| AGAP004811 | CTL | CTL1 | CTL | CTL | not a clip | 1 | 0 | 0.965 | 0.00013 | 27.88145 | 0 |
| AGAP012504 | cSP_E | CLIPE33 | cSP | cSP_E | not tested | 1 | 0 | 1.843 | 0.000639 | 28.9328 | 0.006429 |
| AGAP004975 | PPO | PPO3 | PPO | PPO | not a clip | 1 | 0 | 3.347 | 0.000215 | 30.43384 | 0.008182 |
| AGAP004976 | PPO | PPO8 | PPO | PPO | not a clip | 1 | 0 | 2.425 | 0.000153 | 29.70799 | 0.001234 |
| AGAP004980 | PPO | PPO7 | PPO | PPO | not a clip | 1 | 0 | 13.373 | 0.002952 | 41.48613 | 0.134026 |
| AGAP004981 | PPO | PPO4 | PPO | PPO | not a clip | 1 | 0 | 0.454 | 0.000055 | 21.94864 | 0 |
| AGAP005246 | SRPN | SRPN10 | SRPN | SRPN | not a clip | 1 | 0 | 0.46 | 0 | 18.89171 | 0 |
| AGAP005496 | LRIM | LRIM12 | LRIM | LRIM | not a clip | 1 | 0 | 0.871 | 0.000109 | 26.83182 | 0 |
| AGAP006258 | PPO | PPO2 | PPO | PPO | not a clip | 1 | 0 | 2.306 | 0.001521 | 30.80015 | 0.015455 |
| AGAP006430 | CTL | CTLGA2 | CTL | CTL | not a clip | 1 | 0 | 0.959 | 0.000124 | 27.93169 | 0 |
| AGAP006631 | ModSP | SP214 | SP | ModSP | not a clip | 1 | 0 | 1.903 | 0.000041 | 26.10293 | 0.000325 |
| AGAP012591 | cSPH_A | CLIPA3 | cSPH | cSPH_A | positive_A | 1 | 0 | 1.91 | 0.000323 | 30.06625 | 0.004221 |
| AGAP007034 | LRIM | LRIM11 | LRIM | LRIM | not a clip | 1 | 0 | 4.317 | 0.000169 | 31.84561 | 0.016948 |
| AGAP007280 | ModSP | SP212 | SP | ModSP | not a clip | 1 | 0 | 1.864 | 0.000164 | 27.33992 | 0 |
| AGAP007408 | CTL | CTLMA8 | CTL | CTL | not a clip | 1 | 0 | 4.92 | 0.000624 | 32.84511 | 0.027727 |
| AGAP007410 | CTL | CTLMA5 | CTL | CTL | not a clip | 1 | 0 | 1.433 | 0.000032 | 24.95262 | 0.001104 |
| AGAP007691 | SRPN | SRPN18 | SRPN | SRPN | not a clip | 1 | 0 | 0.907 | 0.000103 | 26.33292 | 0.011364 |
| AGAP007693 | SRPN | SRPN7 | SRPN | SRPN | not a clip | 1 | 0 | 1.838 | 0.000236 | 28.71122 | 0.004675 |
| AGAP013089 | cSP_D | CLIPD20 | cSP | cSP_D | no phenotype | 1 | 0 | 3.427 | 0.000255 | 31.54562 | 0.00461 |

| | | | | | | | | | | | |
|---|---|---|---|---|---|---|---|---|---|---|---|
| AGAP013487 | cSP_B | CLIPB3a | cSP | cSP_B | no phenotype | 1 | 0 | 2.404 | 0.010996 | 34.72804 | 0.021218 |
| AGAP008366 | TEP | TEP2 | TEP | TEP | not a clip | 1 | 0 | 1.386 | 0.00009 | 25.93208 | 0.006558 |
| AGAP028007 | cSP_C | CLIPC13 | cSP | cSP_C | no phenotype | 1 | 0 | 2.506 | 0.000194 | 27.83284 | 0.017662 |
| AGAP008407 | TEP | TEP13 | TEP | TEP | not a clip | 1 | 0 | 0.452 | 0.000013 | 22.49774 | 0 |
| AGAP028069 | cSPH_E | CLIPE26 | cSPH | cSPH_E | not tested | 1 | 0 | 3.634 | 0.000013 | 25.63682 | 0.013442 |
| AGAP028075 | cSP_E | CLIPE9 | cSP | cSP_E | not tested | 1 | 0 | 1.009 | 0.000001 | 18.98176 | 0 |
| AGAP028102 | cSPH_E | CLIPE34 | cSPH | cSPH_E | not tested | 1 | 0 | 2.451 | 0.000008 | 24.13159 | 0.011364 |
| AGAP028167 | cSP_C | CLIPC14 | cSP | cSP_C | positive_M | 1 | 0 | 1.347 | 0.000075 | 25.3845 | 0.000195 |
| AGAP028183 | cSPH_E | CLIPE27 | cSPH | cSPH_E | not tested | 1 | 0 | 4.87 | 0.000082 | 30.06201 | 0.007987 |
| AGAP028229 | cSPH_E | CLIPE28 | cSPH | cSPH_E | not tested | 1 | 0 | 2.395 | 0.000008 | 23.92304 | 0 |
| AGAP029074 | cSPH_E | CLIPE31 | cSPH | cSPH_E | not tested | 1 | 0 | 2.786 | 0.004699 | 30.6942 | 0.013961 |
| AGAP029077 | cSPH_E | CLIPE8 | cSPH | cSPH_E | not tested | 1 | 0 | 0.931 | 0.000055 | 24.26781 | 0.00539 |
| AGAP009316 | CTL | CTL10 | CTL | CTL | not a clip | 1 | 0 | 4.919 | 0.000464 | 34.3738 | 0.007013 |
| AGAP010708 | CTL | CTLMA7 | CTL | CTL | not a clip | 1 | 0 | 7.86 | 0.000513 | 35.69756 | 0.014156 |
| AGAP010709 | CTL | CTL2 | CTL | CTL | not a clip | 1 | 0 | 0.476 | 0.000007 | 21.72545 | 0 |
| AGAP010814 | TEP | TEP6 | TEP | TEP | not a clip | 1 | 0 | 2.696 | 0.000225 | 30.37802 | 0.004026 |
| AGAP010818 | TEP | TEP11 | TEP | TEP | not a clip | 1 | 0 | 4.722 | 0.00036 | 34.06709 | 0.022565 |
| AGAP010819 | TEP | TEP10 | TEP | TEP | not a clip | 1 | 0 | 4.716 | 0.000325 | 32.47158 | 0.019545 |
| AGAP010830 | TEP | TEP9 | TEP | TEP | not a clip | 1 | 0 | 5.223 | 0.00342 | 36.24271 | 0.072532 |
| AGAP010831 | TEP | TEP8 | TEP | TEP | not a clip | 1 | 0 | 5.053 | 0.000377 | 33.92372 | 0.02539 |
| AGAP010832 | TEP | TEP19 | TEP | TEP | not a clip | 1 | 0 | 4.168 | 0.001689 | 32.76003 | 0.03289 |
| AGAP029047 | CTL | CTL5 | CTL | CTL | not a clip | 1 | 0 | 1.449 | 0.000033 | 24.59888 | 0 |
| AGAP029559 | CTL | CTLMA6 | CTL | CTL | not a clip | 1 | 0 | 4.058 | 0.000329 | 32.24865 | 0 |
| AGAP000572 | cSP_C | CLIPC10 | cSP | cSP_C | no phenotype | 2 | 0 | 0.94 | 0.009715 | 27.86501 | 0 |
| AGAP000573 | cSP_C | CLIPC4 | cSP | cSP_C | positive_A | 2 | 0 | 3.482 | 0.025579 | 33.26683 | 0.002338 |
| AGAP001376 | SRPN | SRPN17 | SRPN | SRPN | not a clip | 2 | 0 | 2.701 | 0.028461 | 29.49602 | 0 |
| AGAP001377 | SRPN | SRPN11 | SRPN | SRPN | not a clip | 2 | 1 | 11.044 | 0.136834 | 38.49581 | 0.001818 |
| AGAP002270 | cSPH_B | CLIPB7 | cSPH | cSPH_B | no phenotype | 2 | 0 | 4.22 | 0.049636 | 31.66474 | 0 |
| AGAP002813 | cSP_D | CLIPD6 | cSP | cSP_D | no phenotype | 2 | 0 | 8.967 | 0.102475 | 36.53888 | 0.002792 |
| AGAP003057 | cSP_B | CLIPB8 | cSP | cSP_B | positive_M | 2 | 0 | 8.438 | 0.089949 | 37.68571 | 0.009156 |
| AGAP003246 | cSP_B | CLIPB2 | cSP | cSP_B | no phenotype | 2 | 1 | 14.665 | 0.180023 | 40.41954 | 0.007532 |
| AGAP003247 | cSP_B | CLIPB19 | cSP | cSP_B | no phenotype | 2 | 0 | 4.867 | 0.039876 | 34.49753 | 0.017857 |
| AGAP003250 | cSP_B | CLIPB4 | cSP | cSP_B | positive_M | 2 | 1 | 17.655 | 0.170539 | 42.33712 | 0.040714 |
| AGAP003251 | cSP_B | CLIPB1 | cSP | cSP_B | no phenotype | 2 | 0 | 8.946 | 0.085075 | 35.57884 | 0.00026 |
| AGAP003689 | cSPH_C | CLIPC7 | cSPH | cSPH_C | no phenotype | 2 | 0 | 5.954 | 0.065648 | 34.54148 | 0.000519 |
| AGAP004148 | cSP_B | CLIPB5 | cSP | cSP_B | no phenotype | 2 | 0 | 1.447 | 0.010077 | 28.61984 | 0 |
| AGAP004318 | cSP_C | CLIPC3 | cSP | cSP_C | positive_M | 2 | 1 | 11.852 | 0.133638 | 39.74442 | 0.017143 |
| AGAP004719 | cSP_C | CLIPC9 | cSP | cSP_C | positive_M | 2 | 1 | 11.481 | 0.15036 | 39.64835 | 0.007029 |
| AGAP004855 | cSP_B | CLIPB13 | cSP | cSP_B | positive_M | 2 | 1 | 13.976 | 0.162786 | 39.17605 | 0.000779 |
| AGAP004977 | PPO | PPO6 | PPO | PPO | not a clip | 2 | 1 | 14.895 | 0.185392 | 40.20945 | 0.001169 |
| AGAP005334 | CTL | CTLMA2 | CTL | CTL | not a clip | 2 | 1 | 17.285 | 0.205414 | 42.46863 | 0.01513 |
| AGAP005335 | CTL | CTL4 | CTL | CTL | not a clip | 2 | 1 | 14.228 | 0.181018 | 40.71092 | 0.006071 |
| AGAP005625 | ModSP | SP213 | SP | ModSP | not a clip | 2 | 1 | 14.163 | 0.16263 | 41.50993 | 0.020974 |
| AGAP006348 | LRIM | LRIM1 | LRIM | LRIM | not a clip | 2 | 1 | 15.798 | 0.198071 | 41.10028 | 0.00039 |
| AGAP006910 | SRPN | SRPN3 | SRPN | SRPN | not a clip | 2 | 0 | 9.16 | 0.113343 | 37.08358 | 0.007987 |
| AGAP006911 | SRPN | SRPN2 | SRPN | SRPN | not a clip | 2 | 0 | 6.379 | 0.071514 | 34.05005 | 0 |
| AGAP007039 | LRIM | LRIM4 | LRIM | LRIM | not a clip | 2 | 0 | 7.48 | 0.100243 | 34.81456 | 0 |
| AGAP007692 | SRPN | SRPN14 | SRPN | SRPN | not a clip | 2 | 0 | 5.149 | 0.027357 | 37.99852 | 0.049675 |
| AGAP008364 | TEP | TEP15 | TEP | TEP | not a clip | 2 | 0 | 7.04 | 0.087654 | 34.91586 | 0.00026 |
| AGAP009221 | SRPN | SRPN5 | SRPN | SRPN | not a clip | 2 | 0 | 5.822 | 0.052946 | 33.92141 | 0.011883 |
| AGAP009844 | cSP_B | CLIPB15 | cSP | cSP_B | no phenotype | 2 | 1 | 17.568 | 0.210221 | 42.62997 | 0.017192 |
| AGAP010530 | cSPH_E | CLIPE4 | cSPH | cSPH_E | no phenotype | 2 | 0 | 10.729 | 0.129563 | 39.86685 | 0.011558 |
| AGAP010730 | cSPH_A | CLIPA28 | cSPH | cSPH_A | positive_M | 2 | 1 | 21.308 | 0.235324 | 45.68076 | 0.020065 |
| AGAP010731 | cSPH_A | CLIPA8 | cSPH | cSPH_A | positive_M | 2 | 1 | 17.458 | 0.214655 | 42.20676 | 0.004416 |
| AGAP010815 | TEP | TEP1 | TEP | TEP | not a clip | 2 | 0 | 8.989 | 0.095337 | 37.90545 | 0.02026 |
| AGAP010968 | cSPH_A | CLIPA9 | cSPH | cSPH_A | no phenotype | 2 | 1 | 22.369 | 0.243679 | 47.1486 | 0.064042 |
| AGAP011780 | cSPH_A | CLIPA4 | cSPH | cSPH_A | negative_A | 2 | 0 | 10.668 | 0.085201 | 37.58117 | 0.046039 |
| AGAP011781 | cSPH_A | CLIPA12 | cSPH | cSPH_A | no phenotype | 2 | 0 | 2.354 | 0.016251 | 28.97309 | 0.000779 |
| AGAP011785 | cSPH_E | CLIPE6 | cSPH | cSPH_E | no phenotype | 2 | 0 | 3.391 | 0.037994 | 31.48271 | 0.003961 |
| AGAP011788 | cSPH_A | CLIPA14 | cSPH | cSPH_A | negative_M | 2 | 0 | 7.994 | 0.0856 | 35.06385 | 0.001234 |
| AGAP011789 | cSPH_A | CLIPA6 | cSPH | cSPH_A | no phenotype | 2 | 0 | 5.346 | 0.057946 | 32.25655 | 0 |
| AGAP011791 | cSPH_A | CLIPA1 | cSPH | cSPH_A | no phenotype | 2 | 1 | 18.544 | 0.220723 | 44.13242 | 0.02474 |

| | | | | | | | | | | | |
|---|---|---|---|---|---|---|---|---|---|---|---|
| AGAP011792 | cSPH_A | CLIPA7 | cSPH | cSPH_A | negative_M | 2 | 1 | 17.318 | 0.176375 | 42.65034 | 0.035519 |
| AGAP012034 | cSP_C | CLIPC12 | cSP | cSP_C | positive_M | 2 | 0 | 13.919 | 0.162489 | 40.90828 | 0.010584 |
| AGAP012037 | cSP_B | CLIPB20 | cSP | cSP_B | positive_M | 2 | 0 | 3.423 | 0.051168 | 32.27647 | 0 |
| AGAP012616 | PPO | PPO5 | PPO | PPO | not a clip | 2 | 0 | 9.679 | 0.105006 | 38.31529 | 0.000714 |
| AGAP028725 | cSPH_A | SPCLIP1 | cSPH | cSPH_A | positive_M | 2 | 0 | 11.089 | 0.13005 | 40.05926 | 0.014545 |
| AGAP028728 | cSP_E | CLIPE5 | cSP | cSP_E | no phenotype | 2 | 1 | 13.927 | 0.170779 | 41.35602 | 0.001623 |
| AGAP029100 | cSPH_E | CLIPE20 | cSPH | cSPH_E | not tested | 2 | 0 | 2.667 | 0.010617 | 31.1262 | 0.009416 |
| AGAP029769 | cSP_B | CLIPB9 | cSP | cSP_B | positive_M | 2 | 0 | 0.52 | 0.003493 | 25.49838 | 0 |
| AGAP029770 | cSP_B | CLIPB10 | cSP | cSP_B | positive_M_A | 2 | 1 | 12.948 | 0.15665 | 38.20197 | 0.002338 |
| AGAP006909 | SRPN | SRPN1 | SRPN | SRPN | not a clip | 3 | 0 | 1.339 | 0.010795 | 27.02869 | 0.011364 |
| AGAP007033 | LRIM | APL1C | LRIM | LRIM | not a clip | 3 | 1 | 18.743 | 0.199976 | 43.26772 | 0.043766 |
| AGAP007035 | LRIM | APL1B | LRIM | LRIM | not a clip | 3 | 0 | 1.392 | 0.001181 | 25.14584 | 0.011948 |
| AGAP007036 | LRIM | APL1A | LRIM | LRIM | not a clip | 3 | 0 | 0.497 | 0.000046 | 19.39181 | 0 |
| AGAP007037 | LRIM | LRIM3 | LRIM | LRIM | not a clip | 3 | 0 | 2.471 | 0.021636 | 30.24947 | 0 |
| AGAP007045 | LRIM | LRIM15 | LRIM | LRIM | not a clip | 3 | 0 | 1.376 | 0.009037 | 28.73283 | 0 |
| AGAP008368 | TEP | TEP14 | TEP | TEP | not a clip | 3 | 0 | 7.189 | 0.047531 | 36.94004 | 0.014675 |
| AGAP008654 | TEP | TEP12 | TEP | TEP | not a clip | 3 | 0 | 13.05 | 0.136237 | 40.07611 | 0.014756 |
| AGAP009670 | SRPN | SRPN4 | SRPN | SRPN | not a clip | 3 | 0 | 0.469 | 0.000399 | 20.03214 | 0 |
| AGAP010812 | TEP | TEP4 | TEP | TEP | not a clip | 3 | 0 | 1.823 | 0.017741 | 28.79948 | 0 |
| AGAP010816 | TEP | TEP3 | TEP | TEP | not a clip | 3 | 0 | 11.362 | 0.128425 | 39.36788 | 0.012792 |
| AGAP000290 | cSPH_A | CLIPA27 | cSPH | cSPH_A | no phenotype | 3 | 1 | 14.993 | 0.159405 | 41.59043 | 0.034594 |
| AGAP001648 | cSP_B | CLIPB17 | cSP | cSP_B | positive_M | 3 | 0 | 3.643 | 0.047363 | 31.75679 | 0 |
| AGAP001964 | cSPH_A | CLIPA26 | cSPH | cSPH_A | no phenotype | 3 | 0 | 3.313 | 0.01646 | 32.75825 | 0.011234 |
| AGAP003245 | cSPH_A | CLIPA19 | cSPH | cSPH_A | negative_A | 3 | 0 | 6.048 | 0.039633 | 35.21769 | 0.035195 |
| AGAP009217 | cSPH_B | CLIPB12 | cSPH | cSPH_B | negative_M | 3 | 0 | 6.14 | 0.064068 | 39.27742 | 0.138571 |
| AGAP009849 | cSP_B | CLIPB46 | cSP | cSP_B | no phenotype | 3 | 0 | 2.816 | 0.022482 | 30.24349 | 0 |
| AGAP010545 | cSP_E | CLIPE10 | cSP | cSP_E | no phenotype | 3 | 0 | 9.837 | 0.110494 | 39.77044 | 0.012403 |
| AGAP011787 | cSPH_A | CLIPA5 | cSPH | cSPH_A | negative_M | 3 | 0 | 4.612 | 0.028861 | 32.97984 | 0.017922 |
| AGAP011790 | cSPH_A | CLIPA2 | cSPH | cSPH_A | negative_M | 3 | 1 | 12.922 | 0.162113 | 41.06493 | 0.006185 |
| AGAP001798 | ModSP | SP217 | ModSP | ModSP | not a clip | 4 | 0 | 0.526 | 0.000619 | 26.33126 | 0 |
| AGAP007457 | LRIM | LRIM7 | LRIM | LRIM | not a clip | 4 | 0 | 2.783 | 0.013341 | 33.39569 | 0.006494 |
| AGAP009213 | SRPN | SRPN16 | SRPN | SRPN | not a clip | 4 | 0 | 1.066 | 0.000539 | 27.21082 | 0.005325 |
| AGAP027997 | LRIM | LRIM5 | LRIM | LRIM | not a clip | 4 | 0 | 5.562 | 0.029758 | 38.33764 | 0.040032 |
| AGAP000315 | cSP_C | CLIPC6 | cSP | cSP_C | no phenotype | 4 | 0 | 3.66 | 0.018767 | 31.72341 | 0.001169 |
| AGAP002422 | cSP_D | CLIPD1 | cSP | cSP_D | no phenotype | 4 | 0 | 1.785 | 0.009881 | 30.12996 | 0.000584 |
| AGAP003691 | cSPH_E | CLIPE12 | cSPH | cSPH_E | no phenotype | 4 | 0 | 1.003 | 0.003251 | 27.45088 | 0 |
| AGAP008091 | cSPH_E | CLIPE1 | cSPH | cSPH_E | no phenotype | 4 | 0 | 0.461 | 0.000786 | 23.09149 | 0 |
| AGAP008403 | cSP_E | CLIPE15 | cSP | cSP_E | no phenotype | 4 | 0 | 0.931 | 0.001694 | 24.47263 | 0 |
| AGAP009214 | cSP_B | CLIPB11 | cSP | cSP_B | no phenotype | 4 | 0 | 2.968 | 0.011128 | 31.44803 | 0.012662 |
| AGAP009215 | cSP_B | CLIPB18 | cSP | cSP_B | no phenotype | 4 | 0 | 1.385 | 0.002285 | 25.90156 | 0.000325 |
| AGAP009220 | cSP_B | CLIPB44 | cSP | cSP_B | no phenotype | 4 | 0 | 0.439 | 0.002732 | 26.00186 | 0 |
| AGAP009251 | cSPH_E | CLIPE16 | cSPH | cSPH_E | not tested | 4 | 0 | 2.008 | 0.004099 | 29.82648 | 0.003377 |
| AGAP009252 | cSPH_E | CLIPE17 | cSPH | cSPH_E | no phenotype | 4 | 0 | 11.295 | 0.07893 | 41.42287 | 0.119562 |
| AGAP009273 | cSP_E | CLIPE18 | cSP | cSP_E | no phenotype | 4 | 0 | 1.049 | 0.000968 | 28.42584 | 0.007922 |
| AGAP011786 | cSPH_E | CLIPE7 | cSPH | cSPH_E | no phenotype | 4 | 0 | 0.455 | 0.000179 | 22.31619 | 0 |
| AGAP013184 | cSPH_B | CLIPB36 | cSPH | cSPH_B | no phenotype | 4 | 0 | 3.945 | 0.021619 | 33.46403 | 0.021169 |
| AGAP028641 | cSPH_E | CLIPE29 | cSPH | cSPH_E | not tested | 4 | 0 | 2.389 | 0.004993 | 31.50454 | 0.012987 |
| AGAP029093 | cSPH_E | CLIPE14 | cSPH | cSPH_E | not tested | 4 | 0 | 6.28 | 0.014918 | 37.79605 | 0.063052 |
| AGAP003139 | SRPN | SRPN9 | SRPN | SRPN | not a clip | 5 | 0 | 1.03 | 0.000376 | 23.55756 | 0.000909 |
| AGAP004978 | PPO | PPO9 | PPO | PPO | not a clip | 5 | 0 | 1.472 | 0.001498 | 24.85014 | 0 |
| AGAP006267 | CTL | CTL6 | CTL | CTL | not a clip | 5 | 0 | 1.49 | 0.004221 | 29.59323 | 0.013701 |
| AGAP007407 | CTL | CTLMA4 | CTL | CTL | not a clip | 5 | 0 | 1.376 | 0.001198 | 24.96659 | 0.003831 |
| AGAP007453 | LRIM | LRIM9 | LRIM | LRIM | not a clip | 5 | 0 | 2.368 | 0.010468 | 30.73028 | 0.009351 |
| AGAP007454 | LRIM | LRIM8A | LRIM | LRIM | not a clip | 5 | 0 | 6.812 | 0.029013 | 33.89002 | 0.053474 |
| AGAP007455 | LRIM | LRIM10 | LRIM | LRIM | not a clip | 5 | 0 | 2.838 | 0.00223 | 26.90514 | 0 |
| AGAP007456 | LRIM | LRIM8B | LRIM | LRIM | not a clip | 5 | 0 | 2.304 | 0.002106 | 26.27177 | 0 |
| AGAP009212 | SRPN | SRPN6 | SRPN | SRPN | not a clip | 5 | 0 | 0.918 | 0.001975 | 24.13852 | 0.001494 |
| AGAP012938 | SRPN | SRPN_x | SRPN | SRPN | not a clip | 5 | 0 | 1.974 | 0.000255 | 24.93539 | 0.006753 |
| AGAP003249 | cSP_B | CLIPB3b | cSP | cSP_B | no phenotype | 5 | 0 | 0.489 | 0.001119 | 23.28545 | 0 |

**Table S18: Node enrichment analysis in AgMelGCN2.0.** The significance of under (negative) or over (positive) enrichment in each community within the network was calculated based on the cumulative distribution function (CDF) of the hypergeometric distribution (P< 0.05). k= number of successes; s= sample size; M= Number of successes in the population; N= population size.

| Community | immunoregulatory vs. All | | | | | | | CLIP vs. All | | | | | | | immunoregulatory vs. CLIP | | | | | | |
|---|---|---|---|---|---|---|---|---|---|---|---|---|---|---|---|---|---|---|---|---|---|
| | k | s | M | N | fold enrichment | expected | hypergeometric p-value | k | s | M | N | fold enrichment | expected | hypergeometric p-value | k | s | M | N | fold enrichment | expected | hypergeometric p-value |
| 1 | 5 | 79 | 26 | 177 | -2.32 | 11.60452 | 0.0036 | 40 | 79 | 98 | 177 | -1.09 | 43.74011 | 0.1622 | 5 | 40 | 26 | 98 | -2.12 | 10.6122 | 0.0074 |
| 2 | 16 | 48 | 26 | 177 | 2.27 | 7.050847 | 6.49E-05 | 33 | 48 | 98 | 177 | 1.24 | 26.57627 | 0.0212 | 16 | 33 | 26 | 98 | 1.83 | 8.7551 | 0.0007 |
| 3 | 5 | 20 | 26 | 177 | 1.70 | 2.937853 | 0.1472 | 9 | 20 | 98 | 177 | -1.23 | 11.07345 | 0.2257 | 5 | 9 | 26 | 98 | 2.09 | 2.38776 | 0.0529 |
| 4 | 0 | 19 | 26 | 177 | 0 | 2.79096 | 0.0408 | 15 | 19 | 98 | 177 | 1.43 | 10.51977 | 0.0235 | 0 | 15 | 26 | 98 | 0 | 3.97959 | 0.0063 |
| 5 | 0 | 11 | 26 | 177 | 0 | 1.615819 | 0.1647 | 1 | 11 | 98 | 177 | -6.09 | 6.090395 | 0.0015 | 0 | 1 | 26 | 98 | 0 | 0.26531 | 0.7347 |

| Community | non-phenotypic vs. All | | | | | | | non-phenotypic vs. CLIP | | | | | | |
|---|---|---|---|---|---|---|---|---|---|---|---|---|---|---|
| | k | s | M | N | fold enrichment | expected | hypergeometric p-value | k | s | M | N | fold enrichment | expected | hypergeometric p-value |
| 1 | 20 | 79 | 53 | 177 | -1.18 | 23.65537 | 0.1487 | 20 | 40 | 53 | 98 | -1.08 | 21.63265 | 0.3200 |
| 2 | 16 | 48 | 53 | 177 | 1.11 | 14.37288 | 0.3354 | 16 | 33 | 53 | 98 | -1.12 | 17.84694 | 0.2815 |
| 3 | 4 | 20 | 53 | 177 | -1.50 | 5.988701 | 0.2240 | 4 | 9 | 53 | 98 | -1.22 | 4.867347 | 0.3962 |
| 4 | 12 | 19 | 53 | 177 | 2.11 | 5.689266 | 0.0016 | 12 | 15 | 53 | 98 | 1.48 | 8.112245 | 0.0259 |
| 5 | 1 | 11 | 53 | 177 | -3.29 | 3.293785 | 0.1061 | 1 | 1 | 53 | 98 | 1.85 | 0.540816 | 0.5408 |

**Table S19: Node enrichment analysis in AgGCN1.0.** The significance of under (negative) or over (positive) enrichment in each community within the network was calculated based on the cumulative distribution function (CDF) of the hypergeometric distribution ($P < 0.05$). k= number of successes; s= sample size; M= Number of successes in the population; N= population size.

| Community | immunoregulatory vs. All | | | | | | | CLIP vs. All | | | | | | | immunoregulatory vs. CLIP | | | | | | |
|---|---|---|---|---|---|---|---|---|---|---|---|---|---|---|---|---|---|---|---|---|---|
| | k | s | M | N | fold enrichment | expected | hypergeometric p value | k | s | M | N | fold enrichment | expected | hypergeometric p-value | k | s | M | N | fold enrichment | expected | hypergeometric p-value |
| 1 | 2 | 2787 | 26 | 11794 | -3.07 | 6.1440 | 0.0361 | 4 | 2787 | 85 | 11794 | -5.02 | 20.0861 | 2.30E-06 | 2 | 4 | 26 | 85 | 1.63 | 1.2235 | 3.58E-01 |
| 2 | 4 | 1936 | 26 | 11794 | -1.07 | 4.2679 | 0.5722 | 24 | 1936 | 85 | 11794 | 1.72 | 13.9529 | 0.0042 | 4 | 24 | 26 | 85 | -1.84 | 7.3412 | 0.0655 |
| 3 | 1 | 1605 | 26 | 11794 | -3.54 | 3.5382 | 0.1133 | 9 | 1605 | 85 | 11794 | -1.29 | 11.5673 | 0.2631 | 1 | 9 | 26 | 85 | -2.75 | 2.7529 | 0.1705 |
| 4 | 0 | 1548 | 26 | 11794 | 0 | 3.4126 | 0.0257 | 0 | 1548 | 85 | 11794 | 0 | 11.1565 | 6.11E-06 | 0 | 0 | 26 | 85 | | | |
| 5 | 2 | 899 | 26 | 11794 | 1.01 | 1.9819 | 0.6000 | 5 | 899 | 85 | 11794 | -1.30 | 6.4791 | 0.3627 | 2 | 5 | 26 | 85 | 1.31 | 1.5294 | 0.4866 |
| 6 | 0 | 771 | 26 | 11794 | 0 | 1.6997 | 0.1721 | 2 | 771 | 85 | 11794 | -2.78 | 5.5566 | 0.0772 | 0 | 2 | 26 | 85 | 0 | 0.6118 | 0.4793 |
| 7 | 16 | 489 | 26 | 11794 | 14.84 | 1.0780 | 2.22E-16 | 35 | 489 | 85 | 11794 | 9.93 | 3.5242 | 1.66E-26 | 16 | 35 | 26 | 85 | 1.49 | 10.7059 | 1.11E-02 |
| 8 | 0 | 468 | 26 | 11794 | 0 | 1.0317 | 0.3486 | 0 | 468 | 85 | 11794 | 0 | 3.3729 | 0.0316 | 0 | 0 | 26 | 85 | | | |
| 9 | 1 | 430 | 26 | 11794 | 1.05 | 0.9479 | 0.6197 | 2 | 430 | 85 | 11794 | -1.55 | 3.0990 | 0.3961 | 1 | 2 | 26 | 85 | 1.63 | 0.6118 | 0.5207 |
| 10 | 0 | 216 | 26 | 11794 | 0 | 0.4762 | 0.6181 | 0 | 216 | 85 | 11794 | 0 | 1.5567 | 0.2066 | 0 | 0 | 26 | 85 | | | |
| 11 | 0 | 214 | 26 | 11794 | 0 | 0.4718 | 0.6209 | 1 | 214 | 85 | 11794 | -1.54 | 1.5423 | 0.5415 | 0 | 1 | 26 | 85 | 0 | 0.3059 | 0.6941 |
| 12 | 0 | 127 | 26 | 11794 | 0 | 0.2800 | 0.7544 | 2 | 127 | 85 | 11794 | 2.19 | 0.9153 | 0.2328 | 0 | 2 | 26 | 85 | 0 | 0.6118 | 0.4793 |
| 13 | 0 | 119 | 26 | 11794 | 0 | 0.2623 | 0.7680 | 0 | 119 | 85 | 11794 | 0 | 0.8576 | 0.4210 | 0 | 0 | 26 | 85 | | | |
| 14 | 0 | 32 | 26 | 11794 | 0 | 0.0705 | 0.9317 | 0 | 32 | 85 | 11794 | 0 | 0.2306 | 0.7931 | 0 | 0 | 26 | 85 | | | |
| 15 | 0 | 28 | 26 | 11794 | 0 | 0.0617 | 0.9400 | 0 | 28 | 85 | 11794 | 0 | 0.2018 | 0.8165 | 0 | 0 | 26 | 85 | | | |
| 16 | 0 | 125 | 26 | 11794 | 0 | 0.2756 | 0.7578 | 1 | 125 | 85 | 11794 | 1.11 | 0.9009 | 0.5971 | 0 | 1 | 26 | 85 | 0 | 0.3059 | 0.6941 |

| Community | non-phenotypic vs. All | | | | | | | non-phenotypic vs. CLIP | | | | | | |
|---|---|---|---|---|---|---|---|---|---|---|---|---|---|---|
| | k | s | M | N | fold enrichment | expected | hypergeometric p value | k | s | M | N | fold enrichment | expected | hypergeometric p-value |
| 1 | 2 | 2787 | 50 | 11794 | -5.91 | 11.8153 | 0.0002 | 2 | 4 | 50 | 85 | -1.18 | 2.3529 | 0.5475 |
| 2 | 14 | 1936 | 50 | 11794 | 1.71 | 8.2076 | 0.0270 | 14 | 24 | 50 | 85 | -1.01 | 14.1176 | 0.5717 |
| 3 | 7 | 1605 | 50 | 11794 | 1.03 | 6.8043 | 0.5300 | 7 | 9 | 50 | 85 | 1.32 | 5.2941 | 0.1961 |
| 4 | 0 | 1548 | 50 | 11794 | 0 | 6.5627 | 0.0009 | 0 | 0 | 50 | 85 | | | |
| 5 | 3 | 899 | 50 | 11794 | -1.27 | 3.8113 | 0.4643 | 3 | 5 | 50 | 85 | 1.02 | 2.9412 | 0.6659 |
| 6 | 2 | 771 | 50 | 11794 | -1.63 | 3.2686 | 0.3565 | 2 | 2 | 50 | 85 | 1.70 | 1.1765 | 0.3431 |
| 7 | 18 | 489 | 50 | 11794 | 8.68 | 2.0731 | 5.07927E-13 | 18 | 35 | 50 | 85 | -1.14 | 20.5882 | 0.1748 |
| 8 | 0 | 468 | 50 | 11794 | 0 | 1.9841 | 0.1315 | 0 | 0 | 50 | 85 | | | |
| 9 | 1 | 430 | 50 | 11794 | -1.82 | 1.8230 | 0.4510 | 1 | 2 | 50 | 85 | -1.18 | 1.1765 | 0.6569 |
| 10 | 0 | 216 | 50 | 11794 | 0 | 0.9157 | 0.3961 | 0 | 0 | 50 | 85 | | | |
| 11 | 1 | 214 | 50 | 11794 | 1.10 | 0.9072 | 0.6005 | 1 | 1 | 50 | 85 | 1.70 | 0.5882 | 0.5882 |
| 12 | 2 | 127 | 50 | 11794 | 3.71 | 0.5384 | 0.1010 | 2 | 2 | 50 | 85 | 1.70 | 1.1765 | 0.3431 |
| 13 | 0 | 119 | 50 | 11794 | 0 | 0.5045 | 0.6016 | 0 | 0 | 50 | 85 | | | |
| 14 | 0 | 32 | 50 | 11794 | 0 | 0.1357 | 0.8727 | 0 | 0 | 50 | 85 | | | |
| 15 | 0 | 28 | 50 | 11794 | 0 | 0.1187 | 0.8877 | 0 | 0 | 50 | 85 | | | |
| 16 | 0 | 125 | 50 | 11794 | 0 | 0.5299 | 0.5863 | 0 | 1 | 50 | 85 | 0 | 0.5882 | 0.4118 |
